# Beyond Correlation: Optimal Transport Metrics For Characterizing Representational Stability and Remapping in Neurons Encoding Spatial Memory

**DOI:** 10.1101/2023.07.11.548592

**Authors:** Andrew Aoun, Oliver Shetler, Radha Raghuraman, Gustavo A. Rodriguez, S. Abid Hussaini

**Affiliations:** Taub Institute for Research on Alzheimer’s Disease and the Aging Brain, Columbia University Medical Center, New York, NY 10032, USA; Department of Pathology and Cell Biology, Columbia University Medical Center, New York, NY 10032, USA

**Keywords:** remapping, stability, place cell, grid cell, activity maps, optimal transport, spatial coding, spatiotemporal

## Abstract

Spatial representations in the entorhinal cortex (EC) and hippocampus (HPC) are fundamental to cognitive functions like navigation and memory. These representations, embodied in spatial field maps, dynamically remap in response to environmental changes. However, current methods, such as Pearson’s correlation coefficient, struggle to capture the complexity of these remapping events, especially when fields do not overlap, or transformations are non-linear. This limitation hinders our understanding and quantification of remapping, a key aspect of spatial memory function. To address this, we propose a family of metrics based on the Earth Mover’s Distance (EMD) as a versatile framework for characterizing remapping. Applied to both normalized and unnormalized distributions, the EMD provides a granular, noise-resistant, and rate-robust description of remapping. This approach enables the identification of specific cell types and the characterization of remapping in various scenarios, including disease models. Furthermore, the EMD’s properties can be manipulated to identify spatially tuned cell types and to explore remapping as it relates to alternate information forms such as spatiotemporal coding. By employing approximations of the EMD, we present a feasible, lightweight approach that complements traditional methods. Our findings underscore the potential of the EMD as a powerful tool for enhancing our understanding of remapping in the brain and its implications for spatial navigation, memory studies and beyond.

## 1 INTRODUCTION

The entorhinal cortex (EC) and hippocampus (HPC) have been shown to play a crucial role in spatial navigation and memory (Frank et al. (2000), Chrobak et al. (2000), Fyhn et al. (2004), Buzsáki and Moser (2013), Lever et al.(2002)). Cells in the EC-HPC circuit encode a neural representation of the spatial environment and its associated contexts (Fyhn et al. (2004), Hafting et al. (2005), Moser et al. (2017)). This encoding results in a variety of cell behaviors with different characteristics including, but not limited to, place cells, grid cells, object cells and other vector or feature based coding (O’Keefe and Dostrovsky (1971), Iwase et al. (2020), Lisman (2007), Steffenach et al. (2005), Moser et al. (2008), Solstad et al. (2008), Høydal et al. (2019), Hardcastle et al. (2017), Tsao et al. et al. (2013)). However, the spatial map generated from the coordinated activity of these cell types is not necessarily stable. In fact, multiple studies demonstrated that place cells are able to remap their activity flexibly and in response to small changes in their environment (Anderson and Jeffery (2003), Colgin et al. (2008), Wilson and McNaughton (1993)).

Evidence suggests this remapping of firing activity patterns enables dynamic encoding of different spatial representations, a mechanism that underlies episodic memory (Leutgeb et al. (2005b), Leutgeb et al. (2004), Ferbinteanu and Shapiro (2003)). It is therefore of particular interest to be able to quantify the degree of remapping occurring in different spatial cell types and to be able to characterize the spatiotemporal changes in a given firing rate map. Such firing rate maps also need to be quantified outside the EC-HPC circuit, such as in visual areas, and are not necessarily restricted to position as a dimension.

Currently, spatial remapping has been segregated under two categories thought to represent distinct environmental changes; these are rate remapping and global remapping (Leutgeb et al. (2005b), Colgin et al. (2008)). Rate remapping is observed when testing animals in the same location but changing contextual cues within the environment (e.g. object color) and is denoted by a change in firing rate unaccompanied by a shift in place field location (Leutgeb et al. (2005b)). Global remapping, however, can involve both firing rate changes and shifts in firing fields and can occur both when testing animals in different locations and with certain salient cues (Wood et al. (2000), Kentros et al. (1998), Bostock et al. (1991)). As such the boundaries between the mechanisms that give rise to global and rate remapping are not strictly delineated with respect to the degree of change necessary to trigger them. The underlying processes do however differ in that rate remapping supports continuous information streams and has been observed to occur gradually in response to environmental changes such as morphing of different arena shapes (Leutgeb et al. (2005a), Wills et al. (2005)), while global remapping is an abrupt process where all fields for a given cell remap entirely without intermediate steps (Leutgeb et al. (2005b), Leutgeb et al. (2005a))

Although global remapping is all-or-none within a cell, this is not necessarily the case for the broader population. The presence of partial remapping suggests that global remapping is used to transition between stable and unstable states thus supporting continuous-like information streams through population responses (Wills et al. (2005), Tsodyks (1999)). These differing subsets of reference frames within global remapping need to be characterized to understand how these transitions between states occur and the environmental influences driving them. Additionally, global remapping seems to be a product of both a rate component and a spatial component where, in the former, firing rate is altered and, in the latter, firing place fields can be translated, rotated, scaled or otherwise reshaped. To better understand the specifics of these spatial transformations we need to be able to thoroughly characterize the transitions between states. This requires disentangling the different components involved in remapping. There are three main components that contribute to the spatial maps involved in remapping studies. These are a rate component, a temporal component and a spatial component. The first is well defined in remapping studies however, given that rate coding is not the only information coding schema present in EC and HPC spatial cells, it is important to be able to describe remapping as it relates to alternate information forms such as spatiotemporal coding.

The main methods to identify rate and global remapping are based on Pearson’s r correlation applied to spatial bins on a firing rate map (Leutgeb et al. (2005b), Wills et al. (2005), Hussaini et al. (2011)). This approach is sufficient to identify linear relationships in transitions between firing rate maps on different experimental sessions or conditions. However, Pearson’s r is vulnerable to outliers and cannot effectively capture non-linear transitions nor can it allow for a segregation of remapping types beyond that of pure rate remapping or joint rate-spatial remapping. This correlation approach compares spatial bins at the same position across different ratemaps for a given cell and is most informative when only linear rate changes are occurring. Pearson’s r can quantify simple rate changes but is unable to quantify global remapping beyond its presence or lack thereof. Therefore the profile for global remapping incorporates both a change in firing rate and any type of non-linear shift in the spatial map density that cannot easily be described by a correlation metric. Pearson’s r also requires distributions to be the same in size, owing to its bin to bin approach, which restricts the information that can be carried in a sample of the rate map. This can result in spurious correlations for fields with different sizes and arenas with different shapes.The lack of explicit characterization of the varied transformations observed in spatial remapping is particularly problematic in disease models where seemingly random distortions in fields are seen (Jun et al. (2020), Fu et al. (2017), Ridler et al. (2020), Mably et al. (2017)). We therefore believe a more rigorous approach, focused on spatiotemporal distances that capture non-linear rate transformations, is necessary to further probe the complexities of remapping clearly visible in spatial navigation and memory studies.

One such method that can complement Pearson’s r correlations for rate remapping and extend our ability to identify and describe both spatial and temporal transformations in global remapping is the Earth Mover’s Distance (EMD). The EMD is a distance computed on a pair of distributions (Vasserstein (1969), Panaretos and Zemel (2019)), often applied in computer science for image analysis tasks (Rubner et al. (2000)). This distance when computed using unnormalized distributions can also be referred to as the discrete Wasserstein distance; for normalized distributions it reduces to the Wasserstein distance, but will be described as the ‘normalized EMD’ in this paper to avoid confusing the EMD and Wasserstein distance as wholly separate metrics. The EMD has been shown to be a highly robust spike train distance metric when quantifying temporal similarity patterns in spike trains with varying rate profiles, enabling us to probe the pure temporal component of remapping (Grossberger et al. (2018),Sihn and Kim (2019), Sotomayor-Gómez et al. (2023)). However, the rate component remains the primary source of evidence in current remapping studies while the spatial component, as well as the joint spatio-temporal component, remain poorly quantified.

This study aims to address the challenges in characterizing the remapping process of spatial cell types observed in navigation studies, memory studies, and beyond, with particular focus on non-linear transformations and transitions between states. Current methods, such as Pearson’s r correlation, have limitations in distinguishing simple rate changes from broader whole field changes in rate and space. Therefore, we propose a more rigorous and encompassing approach by employing the Earth Mover’s Distance (EMD). The EMD quantifies the minimum optimal transport distance between two 2D firing rate distributions capturing non-linear non-overlapping transformations and changes in shape and dispersion of fields. We aim to explore how the EMD can enhance our understanding of mechanisms underlying spatial cell remapping and provide a valuable metric for quantifying remapping across different experimental trials/sessions. We consider the use of such a metric for neurodegenerative or otherwise impaired remapping studies by exploring spatial sensitivity and noise robustness. We investigate how generalized approximations of the EMD could be used to manipulate spatial sensitivity properties and identify various cell types associated with points, areas, or other mapped stimuli, such as object and trace cells in the lateral entorhinal cortex (LEC) or point-driven attention mechanisms in visual areas. We also consider the feasibility of applying the EMD given the computational cost by comparing the sliced EMD approximation, which allows for efficient applications of the normalized and unnormalized 2D EMD by computing the average of many 1D EMD values along random image slices (Bonneel et al. (2015)). This approximation is used in all computations in this paper apart from the single-point Wasserstein generalizations (see methods). We further assess the feasibility of applying the EMD given the nature of stability and remapping data where rate effects are varied, and rate changes commonplace. The EMD’s performance is also assessed against the non-linear spearman rank correlation coefficient, and using real life data examples. Through this comprehensive analysis, we underscore the superiority of the EMD over traditional metrics, highlighting its unique ability to explain complex spatiotemporal transformations, effectively distinguish various global remapping patterns, and maintain robustness in the face of noise, rate fluctuations, and other potential distortions.

## 2 METHODS

### 2.1 Synthetic Fields

Place cells and grid cells were modeled as gaussian blobs with fixed standard deviation. Place cell centroids were restricted to bins inside the square map. Place cells were modeled as (17,17) ratemaps with standard deviation varying between 1 and 3 for different figures where standard deviation was consistent. For elliptical fields, two standard deviation parameters are used for the y and x direction. These were set to 1 and 3 respectively. To model remapping, we shift the location of the centroid on a wider map (N*3, N*3) and slice the relevant portion to center the field as needed.

Grid cells consisted of multiple place fields with kernel size and standard deviation parameters. They were organized in a hexagonal pattern across a wider grid (N*8 + kernel * 2) with gapN bins separating place field edges both horizontally and vertically. Slices were taken across this wider grid to obtain (17,17) rate maps of grid ‘cells’. To model remapping, we shift the initial sliced grid map by N*N pairs (from 0 to N) on a wider map (N*8 + kernel*2) and slice the relevant portion to shift the grid phase as needed.

In generating heatmaps, while place field centroids are shifted across the map (left/right/up/down) and tested against a fixed field at the center, grid fields are shifted right by 0 to N and down by 0 to N creating stepwise slices across the wider grid. These slices are tested against the initial sliced grid map. Therefore grid cell examples are not shifted around the absolute center of the grid map but rather moving away from the top left corner. Given that the wider grid pattern is consistent and symmetrical, the information provided by the results should be no different than if a single point at the center of the grid map was chosen to shift around.

Both wider and tighter spaced grid modules were considered by varying standard deviation and gapN parameters. Grid field centroids were not necessarily inside the (17,17) ratemap at all slices taken and a part of the field excluding the centroid could be included in the slice.

All fields were interpolated to (257, 257) for plotting only. Additional examples of synthetic fields without interpolation are provided in the supplementary documents (S1-S3).

### 2.2 EMD

The Earth Mover’s Distance (EMD) is a measure of dissimilarity between two arbitrary un-normalized distributions defined over a metric space with a distance metric *d*(*x, y*). Intuitively, EMD can be thought of as the minimum cost required to transform one distribution into another, where the cost is proportional to the amount of “earth” moved and the distance it is moved. The general formula for the EMD is:

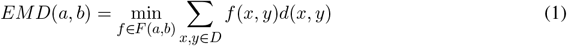

In this equation, *f* (*x, y*) represents the flow from element *x* to element *y*, and d(*x, y*) denotes the ground distance between × and y. F(a, b) is the set of all feasible flows satisfying the following constraints:

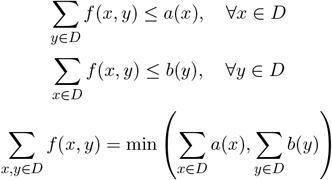

These constraints ensure that the total flow from each element in *a* does not exceed its value, the total flow to each element in b does not exceed its value, and the total flow between the two distributions is equal to the smaller sum of either distribution.

To compute the EMD, one must solve a transportation problem, which is an instance of a minimum-cost flow problem. In the onedimensional case, the EMD has a closed-form solution that is much simpler to compute compared to the multi-dimensional case.

Let P(x) and Q(x) be two one-dimensional distributions defined over the same domain, with cumulative distribution functions *F*_P_(x) and *F*_Q_(x) . The EMD between them can be computed as:

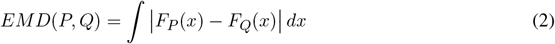

In the discrete case, it can be calculated as:

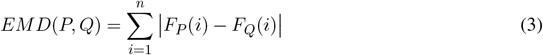

where *F*_*P*_ (*i*) and *F*_*Q*_(*i*) are the cumulative sums of the respective distributions up to index 1. The onedimensional EMD has a closed-form solution because the optimal transport plan is unique and easy to find. In contrast, the multidimensional EMD, which extends the onedimensional EMD algorithm to multiple-dimensional signals such as images or videos, does not have a closed-form solution. The optimal transport plan is more complicated, and the problem becomes a linear optimization problem. Common approaches to computing the multi-dimensional EMD include the Hungarian algorithm or other linear programming techniques, which can solve the problem in polynomial time.

The algorithmic complexity of the multi-dimensional EMD depends on the chosen method for solving the linear optimization problem (Figure S11). For the Hungarian algorithm, the complexity is *O* (*n*^3^), where *n* is the number of elements in each distribution. The 1D Wasserstein however has a closed form solution with runtime O(n). With the python package scipy’s optimized implementation this results in a more efficient runtime than the Pearson’s r correlaton function (Figure S11). For two dimensional distributions, the complexity increases to *O*((*mn*)^3^), for a *m* × *n* distribution. For example, rate-maps of size 16 × 16 (*n* = 16^2^ = 256) would require 16^6^ = 16, 777, 216 operations and a 32 × 32 ratemap would require 1, 073, 741, 824 operations. The computational cost can be reduced in practice by using approximations. In this paper, we use the Sliced Earth Mover’s Distance (Sliced EMD) (Bonneel et al. (2015)).

The Sliced EMD is an efficient algorithm to estimate the EMD between multi-dimensional distributions, such as 2D distributions, by leveraging the closed-form solution for the 1D case. The main idea behind the sliced EMD is to project the multidimensional distributions onto multiple one-dimensional lines (slices) and then compute the 1D EMD on each of these slices. The average of the 1D EMDs across all slices provides an approximation of the multi-dimensional EMD.

The sliced EMD algorithm does the following steps:

- Choose a set of random directions (slices) in the 2D space.
- Project the 2D distributions onto each of these slices.
- For each slice, sort the projections of both distributions.
- Calculate the 1D EMD between the sorted projections using the closed-form solution.
- Average the 1D EMDs across all slices to obtain the sliced EMD.

The algorithmic complexity of the sliced EMD is determined by the number of slices (*L*), the number of points in each distribution (*N*), and the sorting complexity. Since sorting has an average complexity of *O*(*N* log *N*), the total complexity of the sliced EMD algorithm is *O*(*LN* log *N*). For a 16 × 16 ratemap and *L* = 10^3^ projections would be 2, 048, 000. For a 32 × 32 ratemap, the complexity scales to 10, 240, 000. The percent complexity of the Sliced EMD compared to the EMD computed with the Hungarian Algorithm for 16 *×* 16 and 32 *×* 32 ratemaps respectively is 1.22% and 0.95% respectively.

Compared to the Hungarian algorithm and other linear programming techniques, the sliced EMD provides a much more computationally efficient approximation for multi-dimensional EMD, especially when the number of points in the distributions is large. While the accuracy of the sliced EMD may not be as high as the exact EMD computed using the Hungarian algorithm, it often provides a very good balance between computational efficiency and accuracy, making it suitable for various applications where an exact EMD calculation would be too computationally expensive. For two dimensional distributions, the sliced EMD generally converges with between one hundred and ten thousand projections (Figure S11). Even at the upper bound of ten thousand projections, the sliced EMD is substantially more efficient than Optimal Transport techniques. We used the sliced EMD approximation with 10**4 slices to reproduce near-theoretical EMD scores on all figures in our analysis except for field localization figures (10**2).

### 2.3 Single point Wasserstein

In this paper, we primarily use the Sliced Earth Mover’s Distance (EMD) to compare two distributions. However, there is a special use case where the EMD complexity can be further reduced. This use case arises when comparing a normalized ratemap with a distribution that has all its mass concentrated at a single point.

In this case, the EMD formula simplifies to the sum of the distances between the point of interest and each point in the distribution, weighted by the normalized proportion of mass at that point in the distribution. Specifically, let *P* be a normalized ratemap and *Q* be a distribution that has all its mass concentrated at a single point *q*. The EMD between *P* and *Q* is given by:

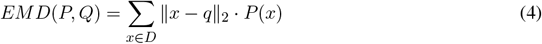

Here, ∥*x - q*∥_2_ denotes the Euclidean distance between x and q, and P(x) represents the proportion of mass at point *×* in the normalized ratemap P. This formula reduces the complexity of computing the EMD from *O* (*n*^3^*∖* log *n*) to *O*(*n*), where *n* is the number of elements in the distribution (Figure S11).

This simplified EMD formula is useful for object remapping, where one is interested in comparing a ratemap to a distribution that represents a single object. For example, it can be used to compare the ratemap of a rodent’s environment with a distribution that represents the location of a reward or danger zone.

### 2.4 Polymorphisms

Here, we outline specific polymorphisms of the EMD that we compute for several use cases.

#### 2.4.1Whole map EMD

The whole map EMD is computed with two ratemaps. For this use case, the Sliced EMD is used with between 10^2^ and 10^4^ projections.

#### 2.4.2 Field EMD

The field EMD is a special use case where only firing fields are considered. In this case, rates are imputed to zero outside of the firing field while rates within the firing field are retained. The Sliced EMD is then used to estimate the EMD on these masked ratemaps.

#### 2.4.3 Binary EMD

The binary EMD is computed using ratemaps where one is imputed for the bins inside the firing field and zero is imputed for those outside the firing field.

#### 2.4.4 Centroid distance

The centroid distance refers to the euclidean distance between two points (e.g. field centroid-object or field centroid-field centroid)

### 2.5 Reference quantiles

Optimal Transport (OT) metrics, encompassing both Earth Mover’s Distance (EMD) and Wasserstein Distances, provide a measure of the dissimilarity between two probability distributions. However, due to their unbounded nature and susceptibility to various external factors, it is often useful to standardize these metrics for meaningful comparisons. To do this, we propose the following steps to transform raw OT values into a relative scale, in terms of their position within a reference distribution.

- Choose a counterfactual distribution to standardize the OT metrics. For example, the counterfactual could be OT metrics from a sampling of mis-matched neurons within the same ensemble.
- When the data is hierarchical (e.g. neurons from different brain regions), it is advisable to compute quantiles separately for each subgroup to ensure meaningful comparisons within the hierarchy.
- For each observed OT metric value, compute the quantile q as follows. Let N be the total number of samples in the counter-factual distribution. Let n be the number of samples less than the observed value. Let q=n/N. This results in a quantile value between 0 and 1 for each observed OT metric.

OT quantiles can be used to assess neural stability and remapping in two main ways. First, the quantiles themselves can be interpreted as one-tailed p-values, gauging the likelihood that each neuron’s observed stability level deviates significantly from what could be expected by chance. The second major way of using OT quantiles is to consider them as a standard scale for group-based hypothesis testing. For example, differences in the representational stability of neurons from two or more groups may be assessed using an appropriate hypothesis test or regression model.

### 2.6 Firing blob extraction

Firing fields were estimated as in (Fyhn et al. (2007)). The peak firing rate was chosen as any bin in the rate map with the largest value (rate). Firing fields were determined as a contiguous region where the firing rate was above 20% of the peak rate. Therefore the top 80% of bins were considered and a blob extraction procedure was then used to extract contiguous regions.

### 2.7 Correlation measures

Pearson’s r correlation coefficients were computed using the python package Scipy. Correlations were computed between the raw and normalized 2D gaussian distributions by comparing the value at each bin in the NxN ratemap. Spearman-rank correlation coefficients were computed in the same way using the spearman function instead.

### 2.8 Data processing

Pre-processing steps for medial lateral entohinbal cortex (MEC) and HPC recordings of mouse models in the lab. Processing steps are also provided for an open-source place cell dataset from the HPC of mice.

#### 2.8.1 MEC & HPC examples

Ratemap dimensions were set to 32 by 32. Firing rates were determined by dividing spike number and time for each bin of the two smoothed maps. Cell recordings were done in rectangular arenas with some HPC cells tested in alternating sessions of circular and rectangular arenas. Spatial information scores were computed as per the Skaggs’ formula which quantifies the information about animal location carried in a spike as bits per spike ((Skaggs et al. (1992)). computation of grid and border scores was done as described .previously ((Bonnevie et al. (2013), Langston et al. (2010)). The largest borders score of the 4 available was chosen.

#### 2.8.2 Place cell dataset

Place cells from hippocampal CA1 two-photon microscopy recordings in mice running across a virtual linear track were obtained from a published dataset (Grijseels et al. (2021a)). Deconvolved spike trains from Suite2P outputs were used directly as cumulative spike counts and converted to 1D firing rates. Δ*F/F*_0_ fluorescence values were computed and preprocessed in the way described by the authors (Grijseels et al. (2021a)). The authors included a normalization step involving 15 second intervals. As such we averaged activity within 15 second intervals (112 frames) to better demonstrate the peak firing rate trend. Given that our current EMD use is not adapted to support negative values, this helped reduced the count of cells that had to be rejected given negative fluorescence. No other filtering was applied and this resulted in 752 unique cells (including spatial and non-spatial cells).

### 2.9 Code

All analyses were done in python using custom code on Jupyter notebooks. All plots were made using matplotlib. Sliced Wasserstein distances on whole maps and field restricted maps as well as binary EMD distances were computed using the python optimal transport package (ot). Map to point distances were computed using custom single-point Wasserstein functions. All functions are integrated as part of our custom analysis toolkit Neuroscikit. A prototype to analyze remapping under different EMD metrics for whole map to whole map cases, object/quadrant cases, field to field cases and specific session grouping cases is provided and being developed as part of this toolkit.

## 3 RESULTS

In order to understand why EMD outperforms Pearson’s r, we need to explore remapping concepts with fine grained control of spatial fields. We therefore opted to synthetically generate examples of field maps where manipulations would allow us to vary 1) field size (stdev) 2) field noise (stdev) 3) field shape (ellipse or circle) 4) field count (1,2 or grid) and, most importantly, 5) field location (x,y) (see methods). We did this by modeling fields as gaussian blobs with centroid coordinates (x,y) on a square map of size N and a standard deviation parameter to vary the field intensity. When necessary and wherever specified, we added normally or uniformly distributed noise across the ratemap. These synthetic fields allow us to approximate certain cell types and different firing field situations that come about in experimentally recorded data. We use these synthetic fields to demonstrate how EMD outperforms Pearson’s r in single, dual and multi-field cases.

### 3.1 EMD captures linear and non-linear transformations resulting in non-overlapping fields

While single field maps can be thought of as a modeling of place cells, the insights derived from the EMD in such situations are applicable to any type of single field map. Therefore, to first compare the Earth Mover’s Distance (EMD) metric to the Pearson’s r correlation coefficient, we considered the simplest remapping case which involves two maps with one identical field in each, but at different locations. This would approximate the simplest case of spatial remapping where every aspect of a field is unchanged except the position on a map. More importantly however, this manipulation allows us to consider remapping transformations that result in non-overlapping fields, a case that Pearson’s r would quantify as no correlation (*p* = 0). Such cases are critical to quantify since remapping resulting in non-overlap still has a biological significance and/or a driver beyond that of having remapped or not. These non-overlapping transformations are often seen in cases of global remapping, and particularly in studies making use of changing arena shapes. In fact, this can be thought of as an analogue to testing remapping across two non-identical arenas where a cell field may move to a now non-overlapping region outside of the initial arena shape (e.g. circular then square arena).

Our synthetic place fields were modeled as 2D-gaussian distributions (*σ* = 1) with a single centroid (center bin) in the middle of a square map. To allow for a single point that is at the true center of the square map, we used N = 17 bins for height and width as opposed to the more traditional (16,16) ratemaps. While one ratemap had its field, and centroid, fixed to the middle of the map, the other was translated across the square such that the centroid had visited every possible bin in the (NxN) ratemap. At every point in a bin, the remapping scores were computed between the fixed and translated rate map. Weights from the ratemap were normalized resulting in a normalized EMD score (Wasserstein distance). As such, we tested remapping across all possible transformations resulting in a field, at least partially, still in the map but not necessarily overlapping with the fixed field. In doing so, we demonstrate that the EMD metric shows greater sensitivity to spatial transformations than the Pearson’s r correlation coefficient (Figure 1). Specifically, the EMD metric is able to capture all possible linear and non-linear transformations resulting in either overlapping or non-overlapping receptive fields. We show that, for a given pair of identical receptive fields, circular or elliptical, the EMD score is non-zero for all possible place field centroids whereas, as one place field from the pair is gradually translated outside the field of the other, the Pearson’s r coefficient accelerates to 0 (Figure 1). In fact, Pearson’s r is unable to quantify any remapping that results in wholly non-overlapping receptive fields (r = 0) while the EMD is non-zero at all possible centroids. Therefore the coverage of information provided by Pearson’s r is restricted to remapping that results in overlap between fields and varies depending on the size and placement of the field in a map.

**Figure 1.**
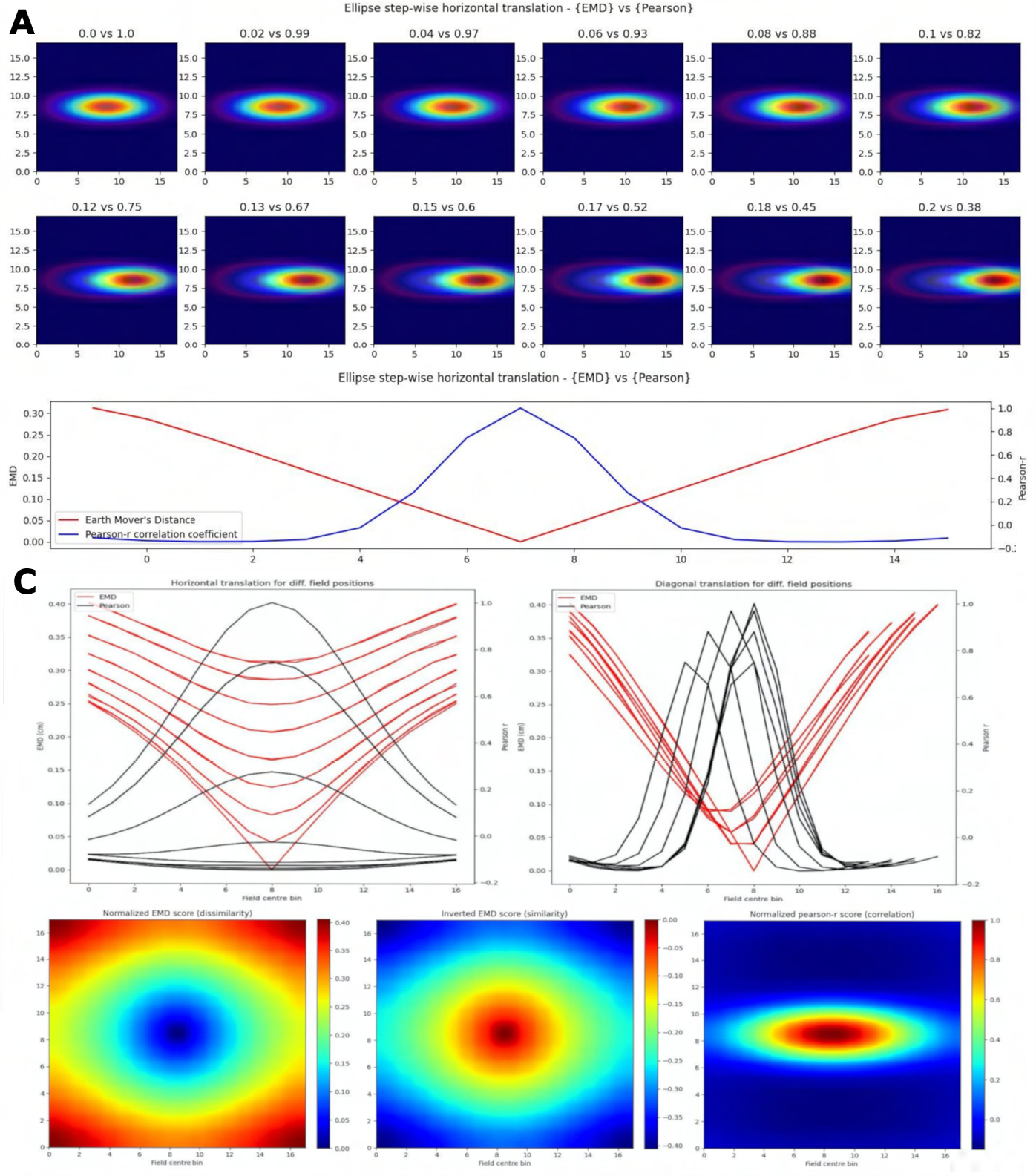
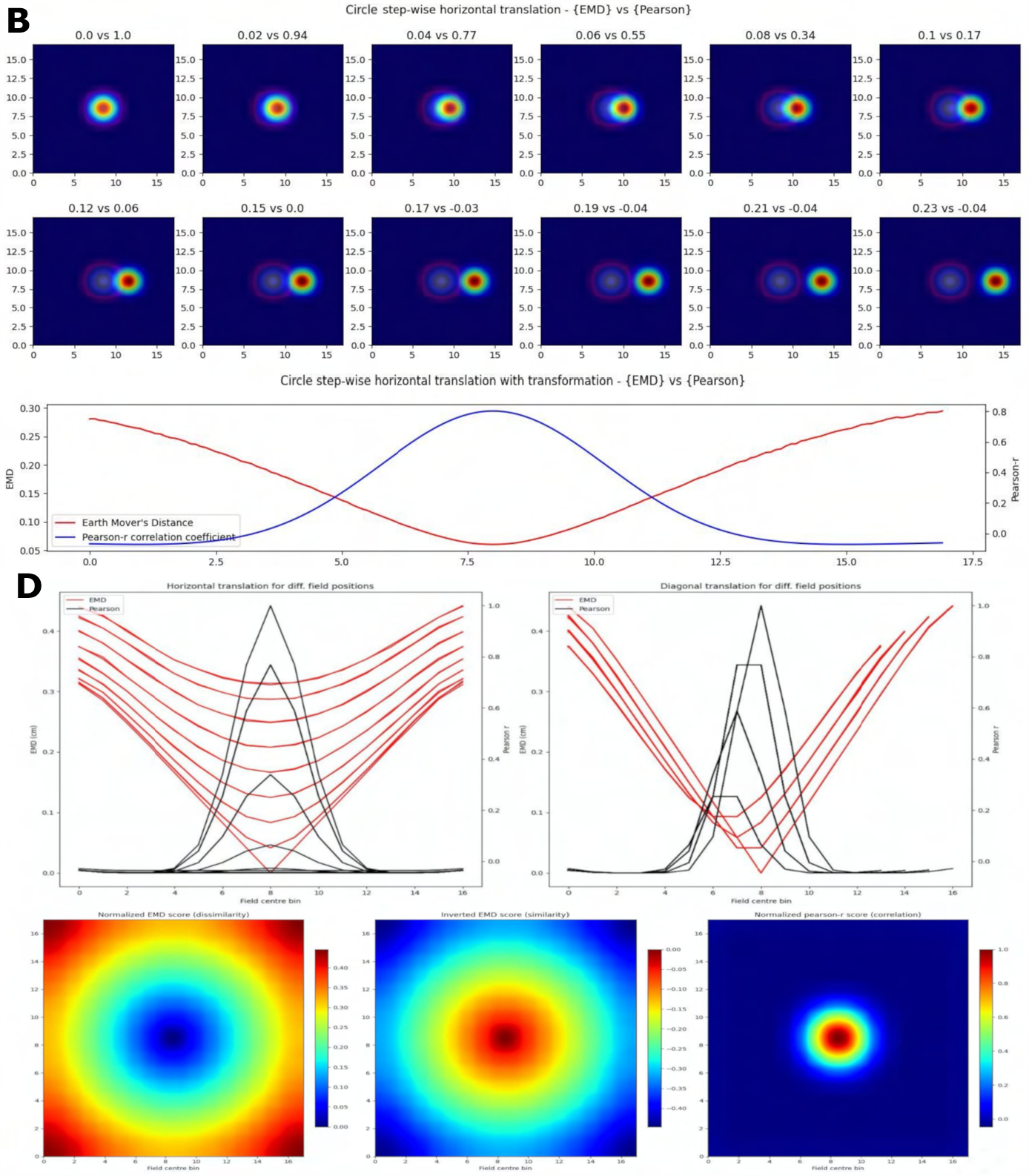
Identical place field translation. Stepwise horizontal linear translation of identical, overlapping place fields (*N* = 17, *σ* = 3) moving from the center to the right (**A, C**). EMD score is shown on the left while Pearson’s r is shown on the right (top panel - EMD vs Pearson). 12 steps are shown and scores are rounded for display. Scores from remapping tested at all possible centroids along a single row on the rate map (bottom panel). EMD and Pearson’s r scores tested at all possible centroids in the rate map (N*N) (**B, D**). Scores for horizontal and diagonal translations along the rate map are shown for all rows (*N* = 17) (top panel). Heatmap showing the gradient of scores for both raw and inverted EMD (left and center) and for Pearson’s r (right) (bottom panel).

Experimentally however, single field cells are not the only observation and various cell types exhibit multiple fields. One such prominent example is the observed pattern in grid cells where multiple fields are laid out in a hexagonal pattern. Combinations of these grid cells coordinate as grid modules with various orientation, spacing and other properties that enable advanced spatial processing using the overlap of multiple grid cells (Hafting et al. (2005), Fyhn et al. (2007)). As such, we then considered the case of overlapping grid modules, both tight and wide in their spacing. In the wide case, we modeled *n* = 3 grid fields with *σ* = 2 and 10 bins of separation. In the tight case, we consider *N* = 3 grid fields with *σ* = 1 and only 6 bins of separation. Given that the map size was *N* = 17, and the requirement to not overlap fields in a grid cell, the decrease in bin separation from 10 to 6 is non-trivial and leads to significantly less spacing between grid fields. In both cases we follow a similar procedure as above and shift the grid module horizontally and vertically for N*N pairs of positions creating a map of EMD and Pearson’s r scores (Figure 2). We demonstrate that EMD also surpasses Pearson’s r in its sensitivity to phase transitions. Specifically, we see that both EMD and Pearson’s r suitably capture the transition from in phase to out of phase but EMD provides a more specific quantification of this boundary. For ease of comparison, we inverted the color scheme of the EMD heatmaps so that hotter colors represent similarity and colder colors represent dissimilarity in a way aligned with how Pearson’s r shows up on comparable heatmaps. In doing so we see that the inverted EMD heatmap demonstrates a broader range of EMD values which allow for a narrower determination of the phase crossing boundary (thinner shaded region at boundary crossing). Additionally, Pearson’s r also results in 0 correlation scores when grid modules are completely non-overlapping, EMD however can still quantify these regions of no overlap. Therefore EMD also offers more spatial sensitivity in the case of grid fields.

**Figure 2.**
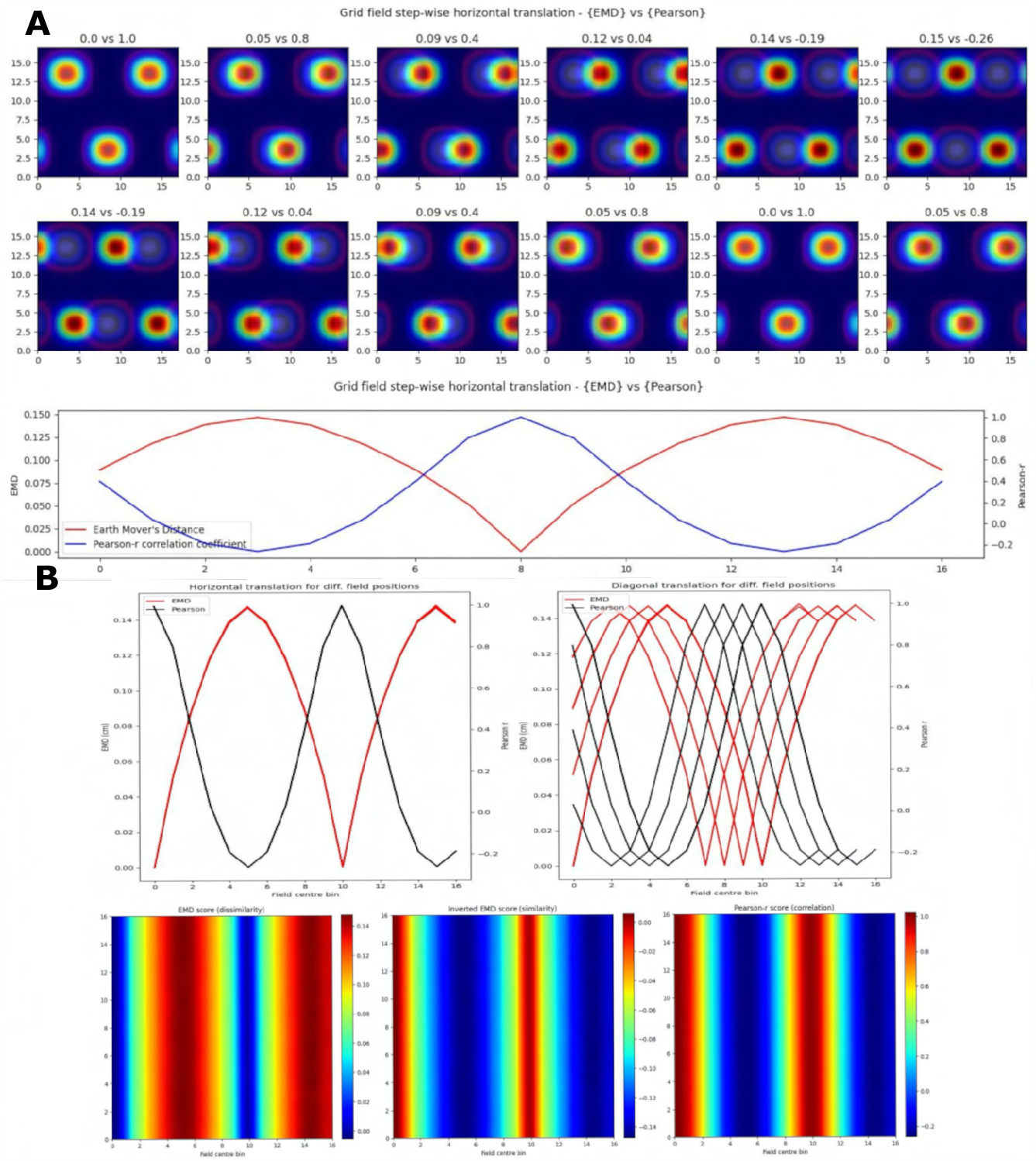
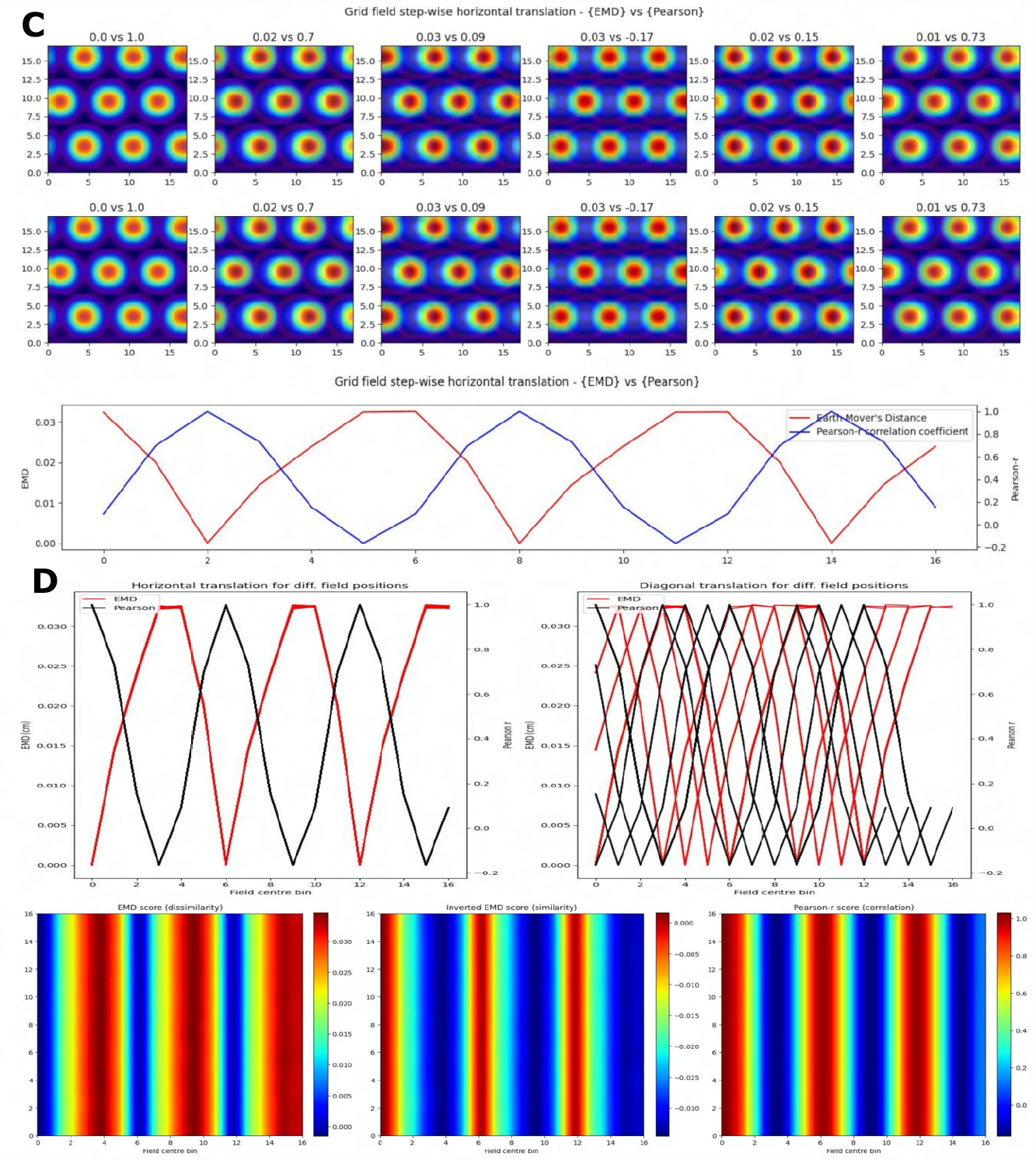
Identical grid field translation. Stepwise horizontal linear translation of identical, overlapping grid fields (*N* = 3, *σ* = 1) moving from the top left corner to the right and/or downwards on a rate map (*N* = 17) (**A, C**). EMD score is shown on the left while Pearson’s r is shown on the right. 12 steps are shown and scores are rounded for display (top panel - EMD vs Pearson). Grid maps were sliced from a larger map with sufficient fields and bins to support N*N steps. Initial grid maps were chosen by taking a slice from the wider map. Scores from remapping tested across N*N different shifts from the initial grid map (0 to N combinations) (bottom panel). EMD and Pearson’s r scores tested at N*N different centroid positions on the wider grid (**B, D**). Scores for horizontal and diagonal translations along the rate map are shown for all rows (*N* = 17) (top panel). Heatmap showing the gradient of scores for both raw and inverted EMD (left and center) and for Pearson’s r (right) (bottom panel).

The EMD is especially useful in cases of no overlap where global remapping transformations struggle to be defined with Pearson’s r. The application of the EMD on place cell transformations resulting in non-overlap demonstrates this (Figure 1, Figure S4). We see reflective properties in that all four corners of the square rate map show the same EMD score gradient. Similarly, the top, bottom, left and right (N,S,W,E) locations show highly similar EMD values. This suggests that the EMD can facilitate the characterization of complex non-linear transformations involving rotations of fields, both overlapping and non-overlapping. This can also be extended to object fields, object trace fields, border fields and any other point or area driven field remapping. Additionally, given that the distribution of EMD scores extends outside the area covered by the gaussian field, this metric may be able to describe remapping and distortions of the underlying field, either in the form of simple scaling or complex degeneration of fields. The increased spatial sensitivity demonstrated by the EMD is also reflected in the elliptical and circular nature of place fields being captured in the underlying EMD distribution (Figure 1, Figure S4). EMD is equally sensitive and informative in the case of non identical place fields, where field shape is varied, as seen with the circle-ellipse pair and the rotated ellipse-ellipse pair further suggesting that the EMD may be used to capture complex changes in the field shape, size and distribution on a rate map (Figure S4).

These complex changes and field distortions are commonplace in neurodegenerative studies such as the observed progressive degeneration of spatial maps in AD mouse models during spatial memory and navigation tasks (Jun et al. (2020), Fu et al. (2017), Ridler et al. (2020), Mably et al. (2017)). Despite the impairments to the underlying spatial map, the remapping of such a map still needs to be quantified to assess differences in stability, learning and memory in these disease states. Additionally, inactivation studies, such as those involving muscimol to inactivate medial septum (MS) or hippocampal input to EC resulting in impaired grid cells and grid cell disappearance respectively (Brandon et al. (2011), Bonnevie et al. (2013)), are commonplace and assessing the stability of spatial maps post inactivation is important. Optogenetic methods have also shown similar findings with altered place and grid cell behavior (Miao et al. (2015), Miao et al. (2017)). To confirm the benefit of using the EMD in cases of impaired cell maps, we considered two more remapping scenarios involving no translations to the field centroid but rather in-place transformations on the field itself resulting in noisy or degenerate spatial maps.

### 3.2 EMD is more stable to noise and field degeneration

In the first case, we considered a scaling remapping of the place field where a gradual increase in the standard deviation of the gaussian field on a fixed (33x33) map results in an increasingly large field (*σ* range = 0 *-* 6). We found that while Pearson’s r was poor at quantifying spatial scaling and insensitive to minor changes, the EMD maintained its spatial sensitivity, and symmetry, allowing for a quantification of both scaling up and scaling down of place fields relative to a single fixed field (*σ* = 3) (Figure 3). The EMD is therefore sensitive across a broader range of standard deviations than Pearson’s r which can only really distinguish large shifts in scaling magnitudes. Pearson’s r is also skewed such that larger standard deviations are less distinguishable than smaller ones while the EMD is robust in both directions. This susceptibility to field size restricts the information Pearson’s r can provide without additional testing. For example, Pearson’s r would indicate greater stability for larger field sizes, but this might simply be due to the increase in correlation bins. Larger field sizes have been associated with disease models such as in AD mice (Cacucci et al. (2008)). Therefore applying Pearson’s r directly would not necessarily be indicative of true stability and would require that the field sizes be normalized before making comparisons, as was done in a previous study (Hussaini et al. (2011)). EMD doesn’t suffer from this problem and has been shown in figure 3A to outperform Pearson’s r.

**Figure 3.**
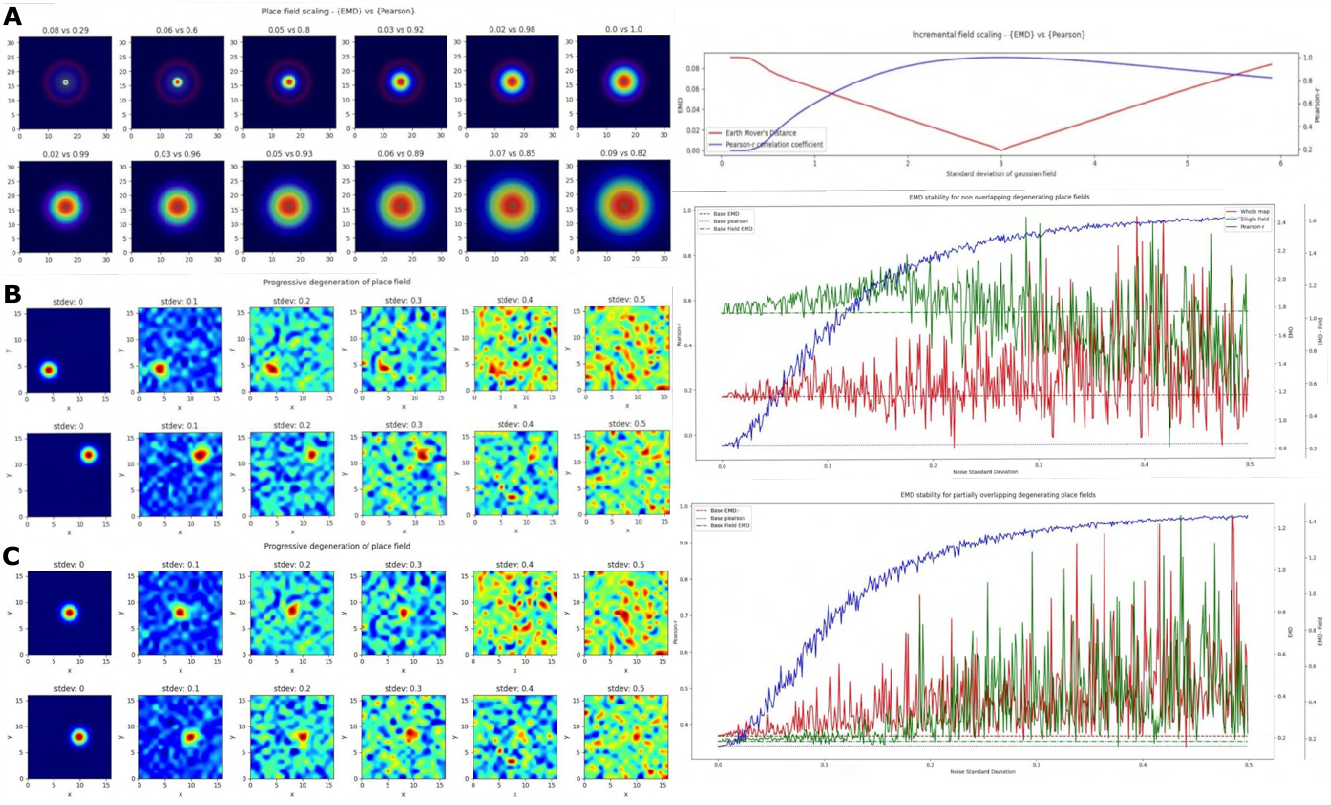
Incremental field degeneration. Stepwise nonlinear translation of overlapping fields (*N* = 33) relative to a fixed field at *σ* = 3. 12 steps are shown with 6 scaling down and 6 scaling up relative to the fixed field (**A**). EMD score is shown on the left while Pearson’s r is shown on the right (left panel). Scores from remapping tested across a range of standard deviations for the scaling field (right panel). Incremental field degeneration for a pair of fields, non-overlapping and overlapping (**B**,**C**). Left panels show the stepwise degradation in the rate map due to randomly sampled normally distributed noise with varying standard deviations. Noise standard deviations are shown above the rate map plots. The distribution plots show the computed remapping score between the pair of fields for the overlapping and non-overlapping cases. Both cases have values for the EMD (red), field EMD (green) and Pearson’s r scores (blue) displayed (right panels).

In the next case, we considered a more complex ‘remapping’ of place fields (17 by 17 pixels) involving progressive degeneration of spatial maps as commonly seen in spatial memory studies of disease models (Jun et al. (2020),Fu et al. (2017)). To model this progressive degeneration of a spatial map and the increased ‘noise’ associated with this degeneration, we added 17 by 17 pixel noise maps in a stepwise manner. The noise fields were sampled from a normal distribution (*μ* = 0) with increasing standard deviation. We did this for two pairs of place fields (*σ*= 1), non-overlapping and partially overlapping place fields (Figure 3). For each we computed the whole map EMD, the Pearson’s r correlation coefficient and a field-restricted EMD following blob extraction based on 20% of the peak firing rate as in (Fyhn et al. (2007)). In the former non-overlapping case, we see significant deviation of the Pearson’s r correlation coefficient from the baseline remapping value of *-*0.0454 computed at no noise. The value is close to 0 since there is no overlap between the fields and bins outside the field are close to 0. We see however that as we increase the added noise, Pearson’s r score accelerates away from the true score and begins to plateau around *r* = 0.8 to 1, solely due to extraneous noise. The distribution of Pearson’s r scores evolving asymptotically is also evident in the overlapping case but with a larger base correlation given the initial overlap (base: 0.339, *μ*: 0.814, *er*: 0.190).

The whole map EMD on the other hand remains more stable relative to the initial no-noise score and deviates fewer times and at larger standard deviations. This trend is seen in the overlapping (base: 0.208, *μ*: 0.369, *er*: 0.171) and non-overlapping (base: 1.17, *μ*: 1.29, *er*: 0.247) cases. We also find that the whole map EMD is more stable at larger standard deviations than the field EMD. In the overlapping case the field EMD shows a greater standard deviation than the whole map EMD despite a similar base and mean (base: 0.183, *μ*: 0.369, *er*: 0.222). This contrasts with the non-overlapping case where the field EMD shows a smaller standard deviation than the whole map EMD (base: 1.04, *μ*: 1.03, *er*: 0.202). These differences in the field EMD and whole map EMD can be understood through the interference of added noise on the detection of firing fields based on peak rate (see discussion). Even so, we find that the EMD metrics are overall more robust to noise and stay closer to the baseline score for larger deviations of noise. As such the EMD scores provide a more stable metric to describe remapping stability than Pearson’s r.

Given that Pearson’s r is so sensitive to outliers and these ratemaps were un-smoothed and un-normalized post added noise, we repeated the procedure by smoothing ratemaps with a gaussian kernel (size= 5, *σ* = 1) and normalizing post smoothing (Figure S5, S6). In doing so, we demonstrate that the observed trend holds and the normalized EMD (Wasserstein distance) again proves to be a more robust metric. We first find that the non-linear trend of Pearson’s r scores across noise standard deviations seen in the unnormalized case remains but shifts to more linear in the smoothed-normalized case. We observe this for both the non-overlapping and partially overlapping scenarios. EMD on the other hand is again a more stable choice and deviates less from the baseline score for overlapping and non overlapping fields. The whole map EMD scores also have noticeably fewer outliers in that there are fewer large peaks away from baseline. We see similar trends in a third test with normalized but unsmoothed ratemaps where Pearson’s r is unchanged from the unnormalized case owing to the inherent normalization procedure of computing Pearson’s r. Therefore EMD is less sensitive to degeneration for both unnormalized and normalized ratemaps, with or without smoothing, emphasizing the use for it in identifying regions of low remapping for particularly noisy or disease-driven degenerate spatial maps We therefore demonstrate that the EMD computed between two ratemaps is robust to noise and degeneration while Pearson’s r is not. The emphasis however is on a robust distance between two non-identical ratemaps (two different cells/sessions) in the presence of increasing noise. While remapping stability is often assessed between different cells or a single cell across different sessions, this is not the only case that can benefit from informed stability analyses. In fact, the EMD should also maintain this noise robustness in a within-cell or within-session comparison as opposed to an across cell/session comparison. This is an important property in order to confidently localize a field on a map and describe regions of high or low remapping as they relate to spatial locations of different objects, points or even other centroids/fields on that same map.

### 3.3 EMD distributions extend field localization and capture remapping relative to a point

To test such within-session stimulus/location-relative stability, we opted to compare the performance of the EMD to Pearson’s r as well as to established field localization methods based on peak firing rates (Fyhn et al. (2007)). Specifically, we sought to understand if the EMD could be used to locate fields and, by extension, to identify specific regions of interest in stability analyses. To do this, we use an adjusted EMD metric computed between the true whole map and a pseudo-map where all the density has been placed in a single bin (single point Wasserstein distance). This process is repeated across all possible bins in the 17x17 spatial map such that a map of EMD scores should be lowest (least dissimilar/most similar) at the region of lowest remapping. In the case of place fields or other single field cells, the point of lowest remapping is presumed to be at or near the centroid of the field. We do this for a rate map with low and high standard deviations of added noise. Correlations are computed between the whole map and point map while field extraction is done on the whole map. To account for border and numerical issues that reflect the scores, we pad the ratemap with N=2 bins (Figure S8). To reduce the influence of outliers on Pearson’s r and firing rates, we smooth the ratemaps after adding noise (Figure S8). We note that smoothing primarily recovers Pearson’s r’s accuracy but not the EMD which remains robust without smoothing (Figure S8). We also find that, as the amount of noise is increased, the EMD is more robust to noise than Pearson’s r and the peak firing rates, with the spatial bins holding the top 20% of EMD scores being less spread out and less sparse than the spatial bins holding the top 20% of Pearson’s r scores or firing rates (Figure 4, FigureS7, Figure S8). This is particularly evident on noise heavy maps where Pearson’s r and peak firing rates lose their specificity towards the region of lowest remapping while EMD retains it. That is, the scores become less specific and therefore less sensitive as a tool to determine the true centroid of a field, locate that field on a map and quantify any associated changes.

**Figure 4.**
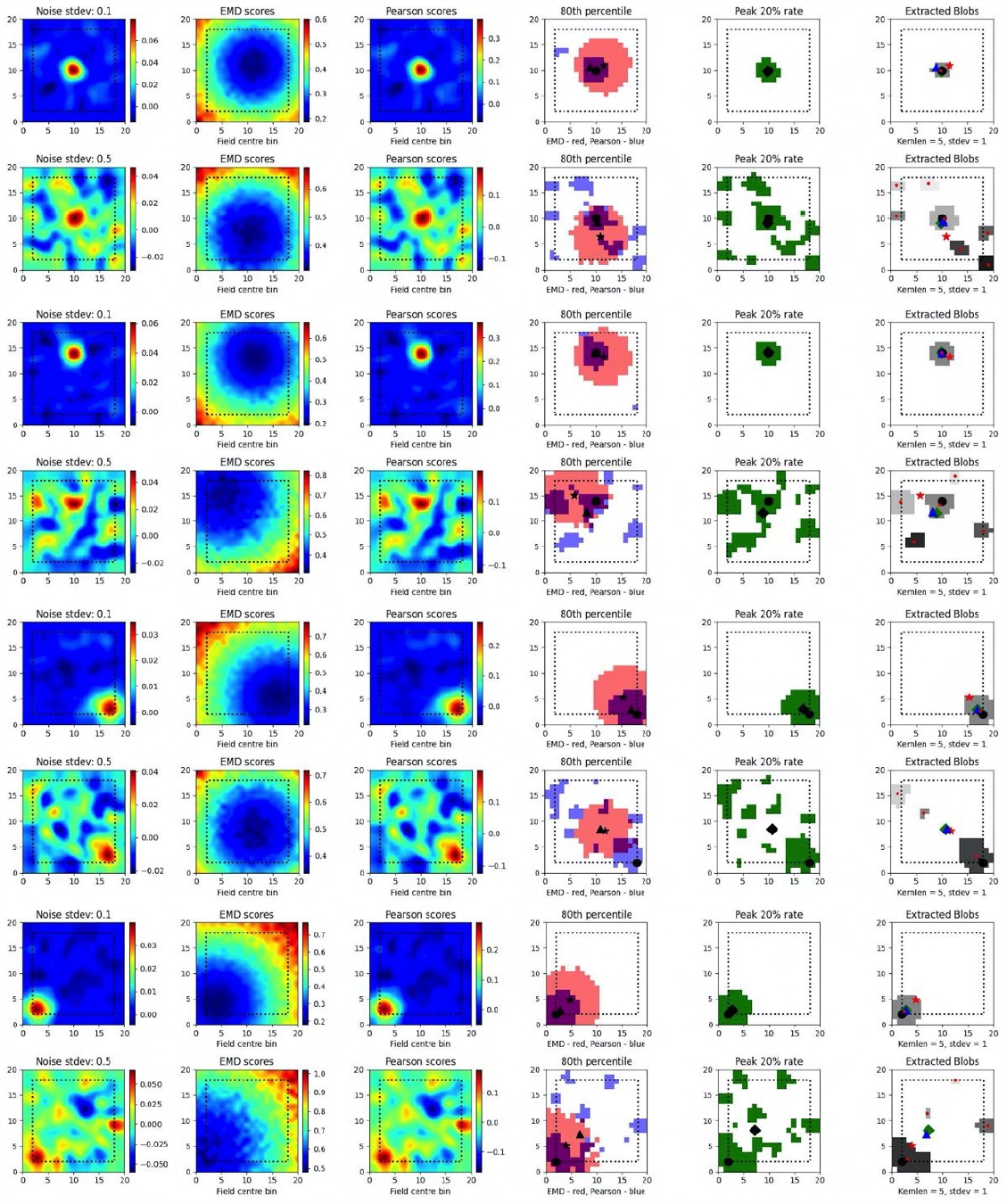
Single field localization. Field localization plots across two different noise levels (rows: low noise 0.1 and high noise 0.5). For each row in a plot, the first column shows the ratemap post added noise with padding, smoothing and normalizing. The second column shows the EMD distribution on the padded rate map with scores being relative to a point map with all the density placed in the bin at which the EMD score is found. The third column shows the same map to point computation for Pearson’s r scores. The fourth column shows the 80th percentile scores for the EMD (red) and Pearson’s r distributions (blue). The fifth column shows the top 20% firing rates in the cell. The last column (sixth) holds the extracted blobs (fields) from the padded ratemap with the centroid of each blob shown in red. The circle represents the true field centroid. The star represents the centroid computed on the peak EMD scores. The triangle is the centroid computed from the peak Pearson’s r scores. The diamond is the centroid from the peak firing rates. The red dots are the centroids of a given field

In fact, with greater noise, the quality is sufficiently degraded to result in multiple fields being detected despite there being a single place field on the original map. These often smaller fields that are outside the region encompassed by the top 20% of EMD scores would be removed experimentally if they were below a certain area threshold. However it is not uncommon for noise blobs (noise fields) to cross this threshold and the number/shape can often vary with ratemap smoothing parameters that are part of the field detection process (Grijseels et al. (2021b)) (Figure S9, Figure S10). As such blob detection algorithms based on a percentage threshold of the peak firing rate may be more robust than other existing methods but can result in too many fields being detected (Figure S9, Figure S10). In the case of single fields, the EMD is a more robust metric to measure remapping relative to a point than the Pearson’s r score and may complement field localization by filtering/validating extracted fields in cases of noisy rate maps.

In the case of dual fields however, the top 20% of EMD values span a contiguous region that includes the area spanned by both fields. While EMD is still robust here, we can see that it does not capture the separate nature of each field at high noise (Figure 5). In fact, while the top 20% of EMD scores are robust to field shape at low standard deviations, this is not the case for larger standard deviations where the map is sufficiently distorted for the EMD to miss the dual field relationship. Despite this we still observe that the relative positioning of the highest EMD region is more consistent and less degenerate at higher standard deviations (Figure 5). We see that the EMD peak regions are dependent on the placement of fields on the map. With multiple fields, the lowest EMD region will be located at a weighted average of all fields, which is the lowest point of remapping, and requires the least amount of work in order to shift all the density to that point as opposed to any other point on the rate map.

**Figure 5.**
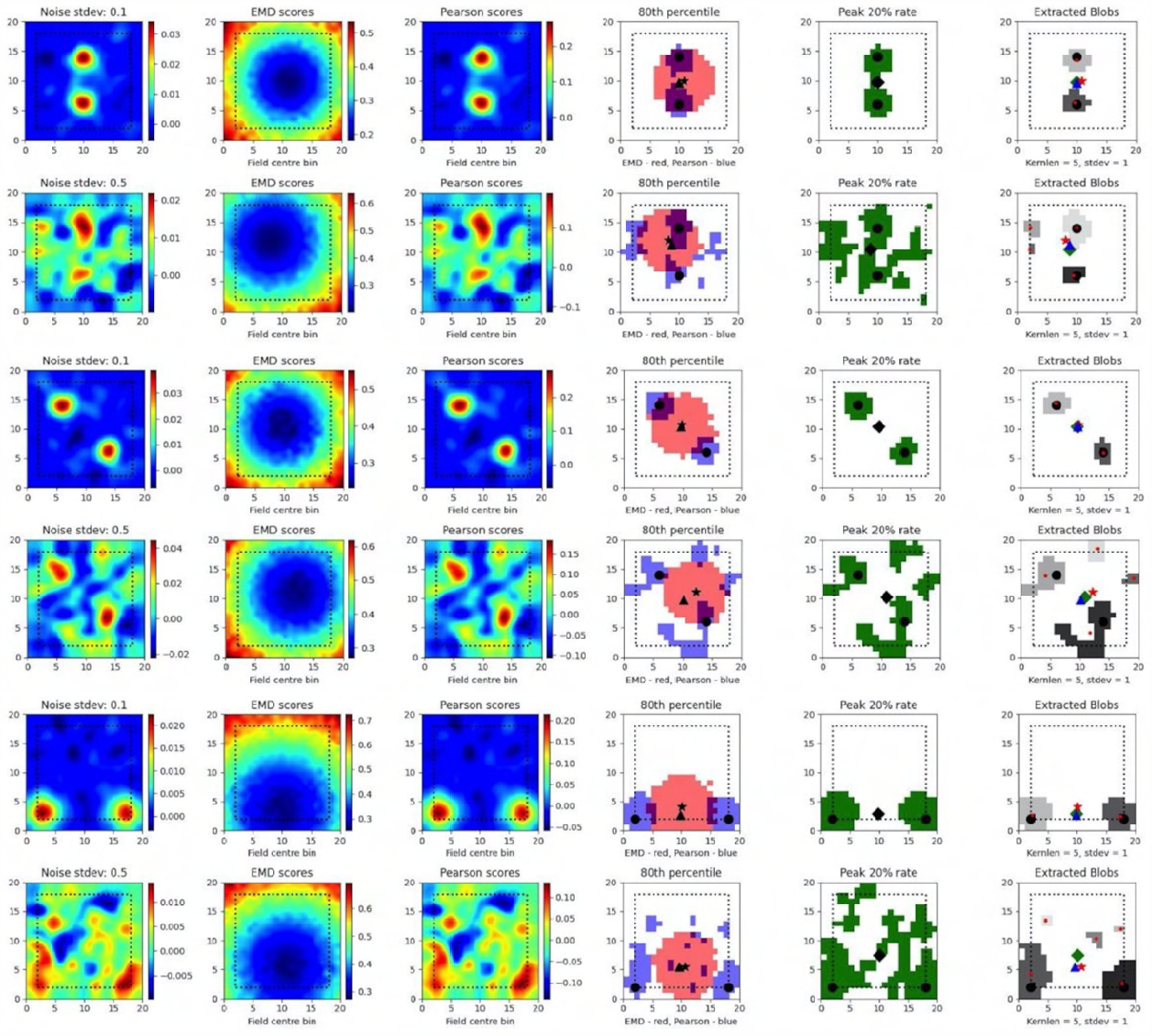
Dual field localization. Field localization plots across two different noise levels (rows: low noise 0.1 and high noise 0.5). For each row in a plot, the first column shows the ratemap post added noise with padding, smoothing and normalizing. The second column shows the EMD distribution on the padded rate map with scores being relative to a point map with all the density placed in the bin at which the EMD score is found. The third column shows the same map to point computation for Pearson’s r scores. The fourth column shows the 80th percentile scores for the EMD (red) and Pearson’s r distributions (blue). The fifth column shows the top 20% firing rates in the cell. The last column (sixth) holds the extracted blobs (fields) from the padded ratemap with the centroid of each blob shown in red. The circle represents the true field centroid. The star represents the centroid computed on the peak EMD scores. The triangle is the centroid computed from the peak Pearson’s r scores. The diamond is the centroid from the peak firing rates. The red dots are the centroids of a given field

As such, the map-to-point EMD approach is still informative in the case of multiple fields and the distribution of EMD values generated by comparing remapping at every possible point allows for a localization of remapping regions. This can be used to identify the region, or quantile, of remapping relative to a known stimulus region (e.g. object location) and could even help filter out detected fields that fall far outside of the contiguous regions of lowest remapping. We also see that this distribution is sustained through increasing noise providing a robust metric to assess spatially correlated and/or driven remapping.

### 3.4 EMD captures object or trace cells and other position/rotation driven remapping

The noise robustness of the EMD using the map-to-point approach, coupled with the symmetrical properties shown and a quantile reference distribution, lends itself to the investigation of specific entorhinal and hippocampal cell types. Specifically, the EMD map-to-point approach involving a quantile reference distribution can be used to track remapping relative to a stimulus point (object location), both current and past (object and object-trace), relative to a stimulus region, both general (N,S,E,W) and specific (single point or multi-location), and relative to current position, both discrete (place cell with single preferred x,y location) and continuous (stepwise sliding window). With the findings of object, location and stimulus encoding playing a central role in EC-HPC circuitry, it is critical that we are able to quantify such remapping to understand its role in episodic memory and changes associated with impairments in this role (Wilson et al. (2013a), Chao et al. (2016), Wilson et al. (2013b), Tsao et al. (2013), Wang et al. (2018)). Additionally, point driven remapping is especially important given theories of EC functioning and mounting evidence for relative temporal tracking of stimulus locations and identities in LEC (Wilson et al. (2013a), Tsao et al. (2013), Wang et al. (2018)). Such point remapping is also found in visual areas where remapping of pointers is suggested to underlie an attention mechanism (Cavanagh et al. (2010)). This evidence further supports the notion of attention-modulated stimulus tracking in the LEC where relative changes in cell responses need thorough characterization to understand their functional role.

In the case of object location tracing, we can provide both relative and specific information. In the relative case, we make use of a distribution of map-to-point EMD quantiles generated by computing distances between a cell field and a series of randomly sampled locations. We can then compute distances to known object locations and describe, relative to the reference distribution, which one leads to the lowest quantile. For more specific tracking involving a distance threshold from the object, we can use a combination of the whole map-to-point map EMD and centroid distances (euclidean distance between field centers). Specifically, for a given NxN ratemap, we compute the EMD between the true rate map and a pseudo rate map where a single bin (object location) holds all the density. We can do this relative to the current session/trial or a previous session/trial. We consider a model experiment setup where an object is shifted across four possible locations and a synthetic object ‘cell’ traces the current location of the object with the placement of its field (*N* = 17, *σ* = 1) (Figure 6). At each object ‘session’, we compute the centroid distance between the field and all 4 possible locations as well as map-to-point EMD values.

**Figure 6.**
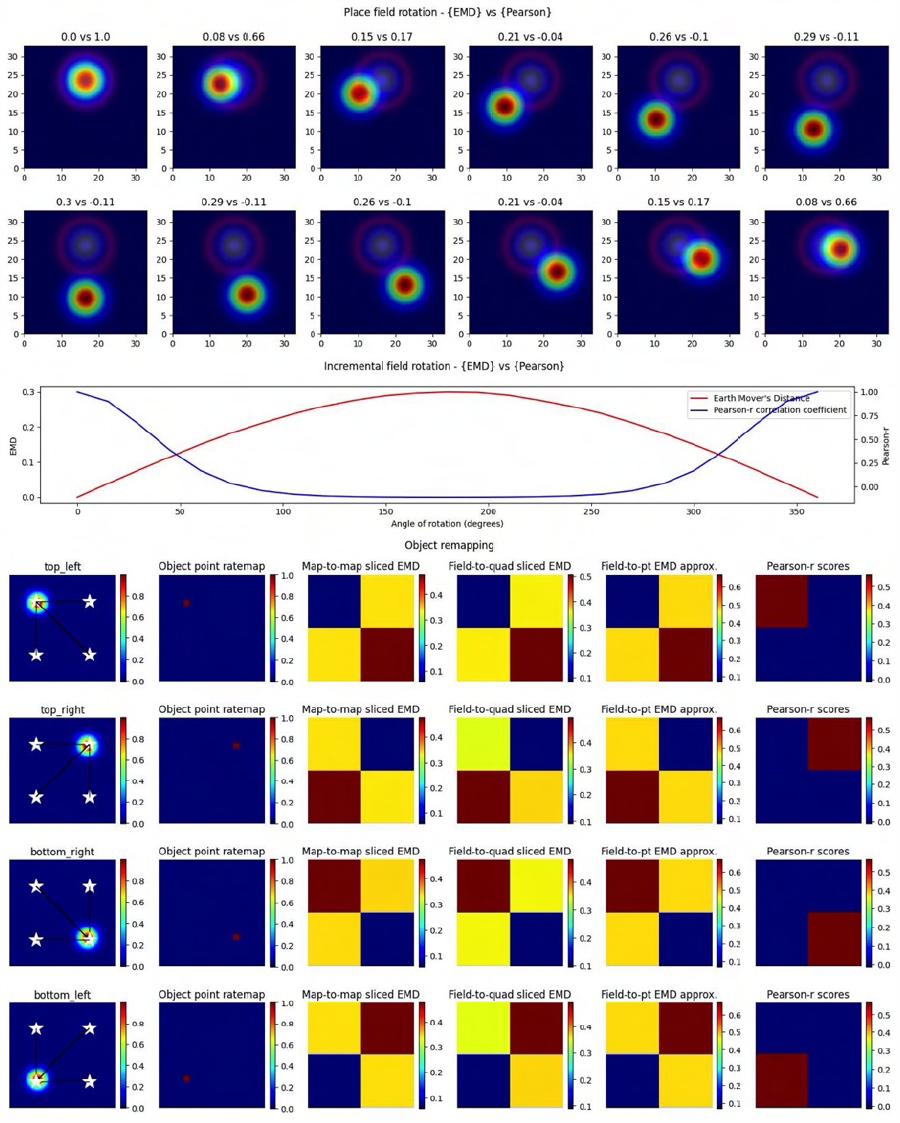
Complex non-linear field remapping. Stepwise nonlinear translation of overlapping fields (*N* = 33) relative to a fixed field at *a* = 3. 12 steps are shown with 6 scaled down and 6 scaled up relative to the fixed field (A). EMD score is shown on the left while Pearson’s r is shown on the right (top panel). Scores from remapping tested across a range of rotation angles (bottom panel). Four corner point driven remapping with top left, top right, bottom right and bottom left tested. Fields were positioned so as to be fully encompassed by the rate map area. The first column shows the field location, four possible object/point/stimulus locations (stars), and distances from the field centroid to each of the four positions. The second column shows the whole map to whole map EMD scores with the full rate map and a pointmap (1 at object location, 0 everywhere else). The third shows a field restricted EMD between a field and a quadrant of multiple bins. The fourth column shows an approximation to the whole map sliced EMD using only the field and the single point object point (single point Wasserstein). The last column holds the Pearson’s r scores. Heatmaps demonstrate the scores in the four possible corners.

We leave the EMD as a distance instead of a quantile since our setup guarantees the object location is at the minimum quantile and we only test 4 points. We see that these values are at their lowest when the field is overlapping with the object location and identical when at opposing yet equidistant locations to the current field. Centroid distances enable us to impose distance requirements and provide more specific localization. In practice, they also provide additional information regarding field shape/dispersion such as when the EMD quantile may be lowest at one object location while the centroid may be closer to another location. Thus the single point EMD (Wasserstein) can be used, along with centroid distances, to determine the object location that requires the least amount of work in moving the field. This demonstrates that we can quantify remapping towards the current object location on a trial, previous trial location or even future trial location (if we suspect the subject’s cell of predicting the change). In doing so, the EMD enables us to identify object, object-trace and/or other types of cell tracking mechanisms.

Importantly however, we again notice the reflective property of the EMD where two possible object positions share the same EMD value despite having different centers. The location of the objects on the map and the vector transformations from the field centroids towards these different locations allow us to differentiate between them despite the similarity in EMD scores. Therefore, a substantial shift in centroid location associated with a low remapping value can be indicative of rotational remapping for a map/field. We can see that this property is not present with Pearson’s r scores which are unable to quantify any rotational remapping or object tracing resulting in non-overlap with r=0 at all points outside of current object location (*N* = 33, *σ*= 3 for rotating field) (Figure 6). Identifying rotational remapping and describing the angle and direction of a rotation is important to understand how these rotations come about and the specific influences that may be driving them or that they may be driven towards. These rotations, either whole map or for a specific field on a map, have been observed experimentally in different contexts including, but not limited to, grid cell rotational realignment in entorhinal cortex (Fyhn et al. (2007)) and place cell cue-driven rotations (Fenton et al. (2000), Muller and Kubie(1987)).

EMD is even more suitable to quantify these rotations and other stimulus driven remappings as it can be computationally simplified for the point map case. Specifically, we show that the properties seen in the whole map to single point map EMD computation are held in cases where the object location is poorly defined or when we are restricted to a field. In the former case, we computed the EMD between only the field on a map and a general quadrant encompassing 1/4 of the square arena and including the current object location. In the latter, we computed a simplified, normalized EMD (single point Wasserstein) between only the field on a map and a singular point rather than the entire point map. In both cases the reflective property that supports complex remapping involving rotations and/or tracing is maintained. Therefore the concept of a whole-map to point-map approach can be generalized to any combination of whole-map/field-map to point-map/quadrant-map/single-point remapping.

The generalization of the whole map - point map EMD into a whole map - single point Wasserstein reduces computation time and provides a simpler and more flexible way to apply the EMD computation. This is highly useful for less specific remapping cases involving no object locations but broader environment differences spanning multiple spatial bins. For example, EMD can also be used to characterize rotational remapping of border cells or remapping towards/driven by entire quadrants/regions of an arena/environment. EMD is also unrestricted by non-overlapping rotational remapping. If we consider remapping in the form of field rotation for a pair of overlapping place fields, we see that Pearson’s r cannot effectively describe rotations. We show that the angle of rotation or presence of a different angle of rotation cannot effectively be distinguished. We find that Pearson’s r is close to 0 for around 100 to 250 degrees of rotation and is blown up below and above those limits. EMD on the other hand is much smoother across all tested angles demonstrating its effectiveness both for rotational remapping that results in either overlapping or non-overlapping fields (Figure 6).

### 3.5 EMD on spatial maps is robust to rate changes

The feasibility of the EMD is not only supported by the various simplifications of its computation (map to point, sliced map, field to field), but also by the rate robustness seen with the EMD in previous studies (Grossberger et al. (2018), Sihn and Kim (2019)) and in our results. Given that firing rate can vary within a session and even more so across sessions separated by days, the EMD benefits from having rate robustness properties which allow for a characterization of spatiotemporal changes alongside the traditional rate remapping quantifications. In the previous study, EMD was shown to be rate robust for temporal coding by varying the total number of spikes. The authors demonstrated that the variability of the EMD across different firing ratios of spike trains was negligible whereas the distance variability for varying degrees of temporal similarity was not, thus allowing for a rate-robust pure-temporal spike train distance (Grossberger et al. (2018), Sihn and Kim (2019)). We also see this rate-robustness in our data where EMD values do not vary significantly with different firing rate ratios using normalized EMD distances. Additionally, it is common practice in field comparison studies to discard low firing rates to reduce spurious correlations caused by noise. While this can help reduce noise effects, there is an inherent loss of information associated with the change in computed firing distribution. Therefore having a rate robust metric that can include all spikes with minimal impact from noise is especially important.

As such, we propose the use of a binary EMD to describe field dispersion regardless of the rate distribution. We demonstrate this rate robustness using the binary EMD generalization alongside normalized and unnormalized whole-map and field-map EMD distances (Figure 7). To do this, we used the scaling of place fields as a type of rate transformation. We can view this transformation as an increase in field intensity where larger intensities indicate higher firing rates while smaller intensities indicate lower firing rate. For a fixed field, scaling its activity across sessions, we computed three measures of EMD remapping involving whole map to map EMD, field to field EMD and a binary EMD that imputes 0 outside a field and 1 inside a field thereby silencing rate effects and varying solely with density dispersion. In practice, one could compute two separate ‘binary’ metrics where one is field restricted and imputes 0 or 1 as described while another spike density EMD would use all raw spike positions (x,y for a given spike) directly without distributing them into a weighted (NxN) firing rate map (whole map dispersion). Since we synthesized our data for these simulated cases, we can only apply the binary EMD and not the whole map spike density EMD. We repeat the selected EMD measures on our rate maps for normalized (Wasserstein) and unnormalized (EMD) intensity changes (place field scaling) (Figure 7). Given that the binary EMD uses raw spikes rather than firing rate maps, we find that the scores are the same in both the normalized and unnormalized case. As such, the binary EMD provides a rate robust metric to quantify the dispersion of rate maps and/or the dispersion of the underlying field in the ratemap, regardless of the rate distribution across the map/field. This binary EMD approach can thus be used to describe remapping without the influence of the rate distribution.

**Figure 7.**
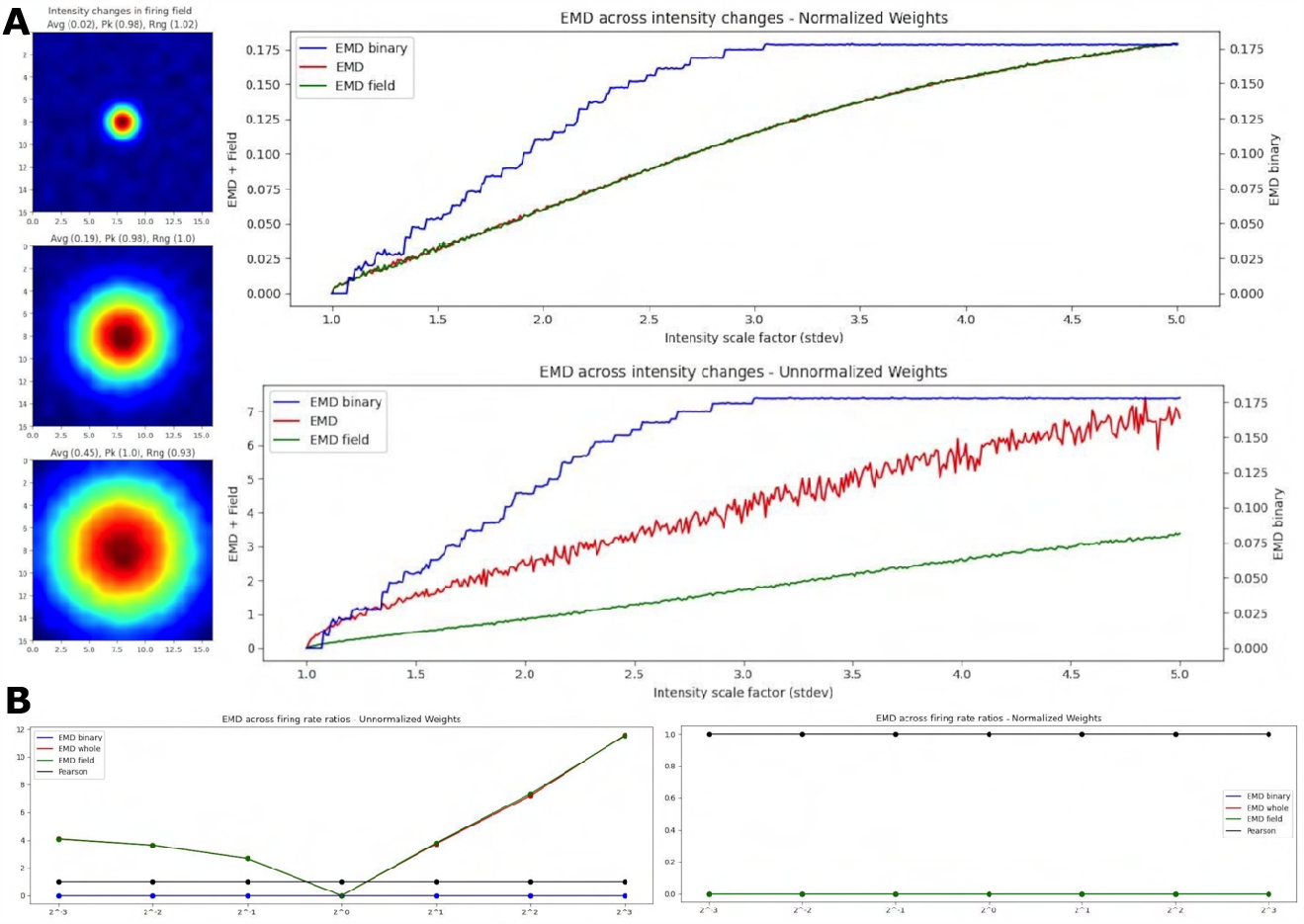
EMD robustness to intensity changes. Incremental field scaling (increasing intensity) across a (17,17) ratemap with a field at the center (**A**). Field is scaled from = 1 to *σ* = 6 (left). Intensity changes are considered using whole map EMD (red), field restricted EMD (green) and a binary EMD (blue) using raw spike positions. Normalized (top right panel) and unnormalized (bottom right panel) weights are shown for both. EMD scores tested against different firing rate ratios for two identical fields (**B**). Ratios greater than 1 and less than 1 were tested.

In the case of EMD measures including rate influences, we test both the whole map EMD and the field restricted EMD. In the normalized case, we see that the field EMD (Wasserstein) and the whole map EMD (Wasserstein) show similar gradients/rates of change across intensity scaling factors (standard deviations). This is not the case when unnormalized where, as we increase the intensity, the whole map EMD and field EMD increase their separation. When compared to the normalized case, this is reflective of the use of raw firing rate scores which, at larger standard deviations, result in larger differences between whole map EMD and field EMD. However, the use of the normalized and unnormalized EMD results in similarly evolving trends for the distance metric enabling a rate robust interpretation of the results. Therefore, when comparing two identical spatial maps with varying firing rate ratios, the raw EMD value itself is highly stable if using the binary EMD or the normalized EMD but varies when using unnormalized rates (Figure 7). These differences therefore allow the EMD to be manipulated in such a way that rate effects can be separated from spatiotemporal changes and the underlying shifts in shape and dispersion described regardless of intensity changes. As such, we can investigate remapping both with and without rate effects. We can characterize remapping for reduced rate effects (normalized EMD == Wasserstein distance) and raw rate effects (unnormalized EMD) and compare these to binary or spike density EMD scores to separately quantify the contribution to remapping provided by rate and/or spatiotemporal changes.

### 3.6 EMD outperforms linear and non-linear metrics

While Pearson’s r was the primary metric tested alongside EMD, owing to its popular use in remapping studies, it is not the only plausible metric to apply. As such, we also thought it relevant to compare the EMD and Wasserstein metric to another common metric. Since Pearson’s r is a strong linear metric, we opted for a non-linear metric found in the Spearman rank correlation coefficient. We repeated all figures with the spearman rank coefficient instead of the Pearson’s r coefficient and provide a sample of key figures in the supplementary documents (S12 - S18). We find that the EMD remains the superior choice as an overall more sensitive and robust metric. Specifically, we see that Spearman- *ρ* actually outperforms Pearson’s r in its spatial sensitivity but still cannot effectively capture all transformations resulting in non-overlap. While in the case of two overlapping circular fields of identical shape Spearman- *ρ* is fully symmetric like the EMD, it does not maintain this property across different field shapes with the overlapping ellipse pair (same angle of rotation) only allowing for a distinction of left and right as opposed to all four corners.

In fact, despite being a non-linear metric, Spearman- *ρ* cannot effectively quantify all types of complex, non-linear transformations. This is further seen in its inability to quantify overlapping identical fields across different intensity scales. Spearman- *ρ* is however capable of describing transformations such as rotations or, depending on the field shape, single point/quadrant remapping. This however is highly restricted by noise, as is the case with Pearson’s r. We find that Spearman- *ρ* is not robust to noise and quickly degenerates away from the baseline, no-noise Spearman score. Spearman- *ρ* is therefore similarly susceptible to Pearson’s r despite the added nonlinear sensitivity. Overall however, the EMD’s non-linear selectivity is more encompassing, more robust and more effective at quantifying and characterizing the spatial transformations underlying remapping.

### 3.7 EMD applied to real spatial data

The effectiveness of the EMD spans a broad range of spatial transformations seen in a variety of contexts, and with different brain regions. Synthetic data alone however cannot recreate all the intricacies present in spatiotemporal representations. As such, we offer examples of the EMD’s increased spatial sensitivity using activity maps from recorded cells. We offer a combination of individual examples that depict specific cell types, or representative maps, as well as an open-source place cell dataset from the hippocampus (HPC) (Grijseels et al. (2021a)).

In our individual examples, we share cells from both the MEC and HPC of AD mouse models (Figure 8). We also provide additional examples from the HPC of AD mouse models and the MEC of non-AD models (Figure S19-S20). We compute the remapping distance, and correlation, between the spatial map of matched cells on a first, reference session, and a subsequent, shifting session. We shift the center of the latter session map to all possible bins in the ratemap and recompute stability measures to assess how the gradient of EMD and correlation values support quantifying remapping. Given this approach, we anticipate a smooth gradient of EMD values as the second session is shifted further or closer from its center. As we shift the second map, we add constant values of 0. However, for MEC examples only, we also shift with wrapping such that no aspect of the map is replaced with 0s (Figure 8A). This is because of the grid symmetry in MEC where it is more appropriate to treat maps as part of a wider grid module. We include both low noise and high noise maps and provide a selection of high grid, high border and high spatial information score examples (Figure 8, Figure S19-S20).

**Figure 8.**
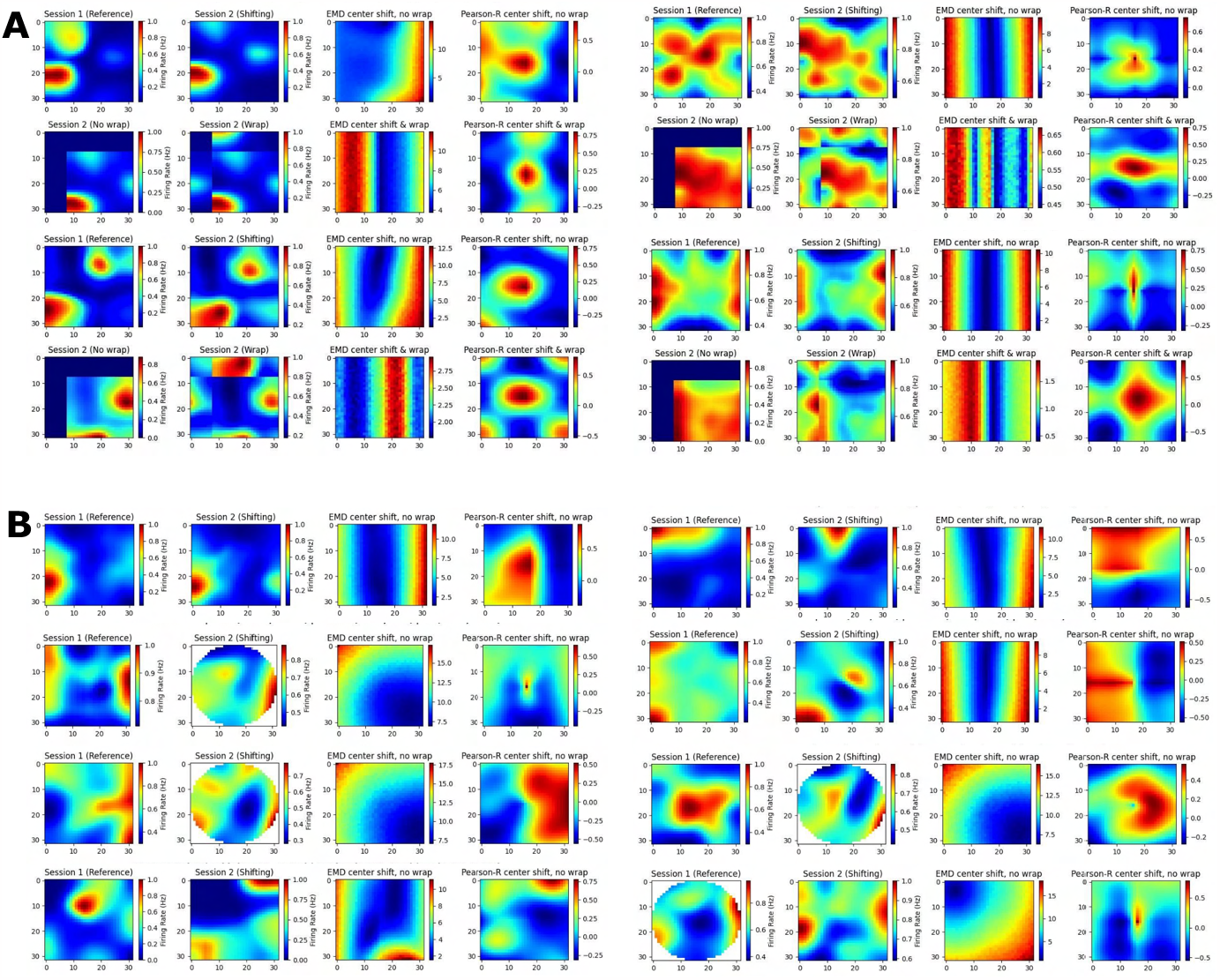
Individual cell examples. Examples of ratemaps from the MEC and HPC of AD mouse models for a reference session and a shifting session. Gradient of Pearson’s r scores tested at all possible map shift centers (N*N) is shown to the far right of each cell example with the EMD gradient immediately to the left of it. For each MEC example, the top row demonstrates a shift with no wrap (0 padding) while the bottom row demonstrates a shift with wrapping (**A**). For each HPC example, only the no wrap row is provided, and examples of matched cells across circular to rectangular arena transitions are included (**B**).

In doing so we find multiple strengths of the EMD discussed earlier. Primarily, we observe that the EMD is much more effective at describing remapping and is not greatly influenced by these small shifts, as evidenced by the smooth gradients of EMD values. Pearson’s r on the other hand shows its lack of spatial sensitivity. For high noise examples, we see that spurious correlations can cover the entire map. For low noise examples, as well as sparse firing, we see that 0 correlation scores can cover significant regions of the map and prevent the true degree of similarity from appropriately being quantified. In fact, when map noise is particularly widespread or firing is especially sparse, Pearson’s r gradients can be fragmented and misleading in the information they carry. Correlation in practice is therefore too ‘brittle’ and non robust to minor spatial shifts. Given the focus of correlation on bin to bin similarity, we show how these minor shifts, commonplace in experimental data, can result in similar maps obtaining a low correlation score. On the other hand we also show how two dissimilar maps can incorrectly be classed as highly correlated given widespread noise in the ratemap. The distributional aspect of the EMD approach rescues these effects.

Additionally, we see that Pearson’s r suffers from this bin to bin approach in experiments involving changing arena shapes. We apply the same approach to cells recorded from the HPC of AD mouse models tested on alternating rectangular and circular arenas (Figure 8B). We note that the EMD can be computed between two distributions no matter the size difference while Pearson’s r requires a bin to bin comparison and is therefore restricted by the smaller distribution’s size. As such, while correlation values can be computed for every possible center point, certain post-shift maps involve fewer bins with which to compute a correlation. Therefore we find a smooth gradient for EMD values while Pearson’s r struggles to accurately capture the similarity between the two distributions, particularly for center shift bins in the corners of the map (outside circular arena).

Examples so far have focused on 2D spatial maps, often with multiple fields. Pearson’s weaknesses however are not restricted to 2D spatial maps, nor to multi-field cells, and also extend into relative decoding. Specifically, results with synthetic data suggest that Pearson’s cannot effectively support similarly between a reference point, or set of reference points. In a further example, we test the EMD on place cells obtained from head-fixed two-photon recordings in mice running across a VR linear track (Figure 9, Figure 9S). We use the deconvolved suite2p spike outputs directly to compute 1D rates in each bin for each cell in the dataset (Figure 9). We do not filter out specific cells based on noise, firing rate, spatial information or any other metric. Instead we offer the full dataset of linear ratemaps as a noise heavy dataset including both place cells and non-spatial cells in the hippocampus. For each cell, we compute the remapping distance and correlation score relative to a pseudo-map where all the density is placed in 1 bin, across a window of bins or distributed with a gaussian template.

**Figure 9.**
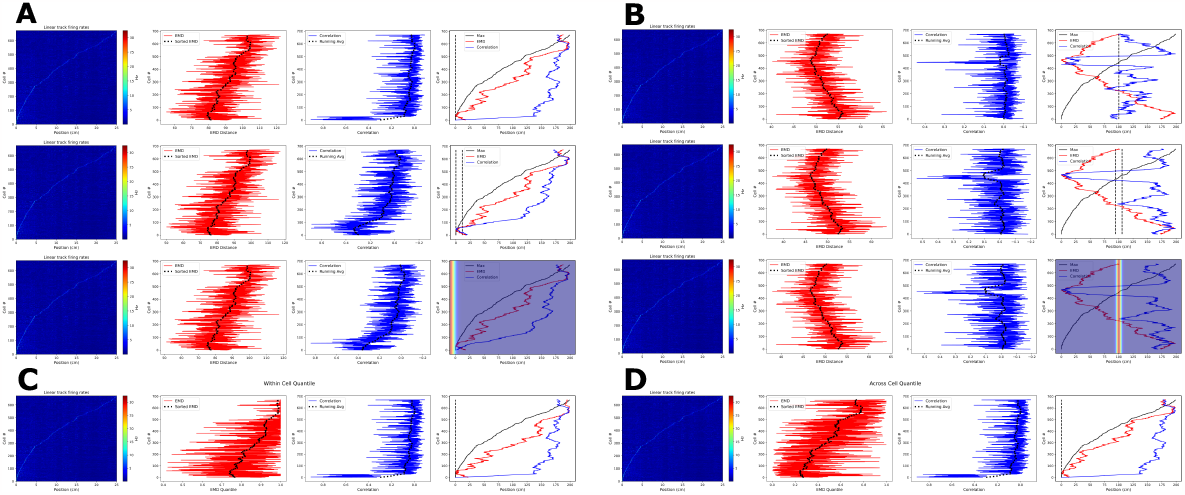
Place cell population decoding. Place cell population decoding using a reference template and a population of 1D rates across a 200cm linear track. The first plot in a row shows a population map of linear firing rates. The second plot shows the distribution of EMD values, and the running average. The third plot shows the population of correlation values, and the running average. The last plot shows the peak firing rate trend overlaid with the running averages of each score distribution. Two sets of examples are provided with reference templates highest in density at the start (**A**) and in the center (**B**). In each set, the first row of plots uses a reference map with all the activity in a single bin. The second row uses a map with the activity spread out across 10 bins. The last row uses a 1D gaussian template with sigma = 5. Reference points, and windows, are plotted with a dashed black line in the final plot of each row. Reference gaussians are overlaid in the final plot of each row. Population decoding with quantiles instead of distances. Quantiles are computed using the distribution of EMD distances from all other cells (**C**). Quantiles are computed using the distribution of EMD distances from a given cell relative to random reference locations (**D**).

We consider two reference locations: the start and middle positions of the linear track. In the former case, given that we sort cells based on the peak rate, we expect a successful measure of relative similarity (or dissimilarity) to capture the same trend as the sorted peak firing rates. In the latter case, given the reference point is in the middle, we expect a successful measure to decrease in similarity (or increase in dissimilarity) as we move in either direction away from the center point. Therefore here we expect a somewhat ‘v’ shaped trend that peaks at the middle position. We find that the EMD demonstrates this expected behavior on both cases while the bin to bin approach of Pearson’s r is unable to quantify relative differences in stability between each cell map and the reference map. We also find that the EMD score is robust to the size of the window, and does not vary greatly if a gaussian template is used instead. Pearson’s r on the other hand shows more spurious increases in similarity as the size of the window is increased.

We note that, in a preprocessing step to generate 1D linear firing rates, we average deconvolved spikes across multiple frames to reduce the number of bins. This creates a bin parameter to which correlation and not EMD is vulnerable. This is again owing to the distributional aspect of EMD which allows for stability despite these hyperparameters while correlation’s bin to bin approach does not. Given that the authors of the dataset used Δ *F/F*_0_ values in their analysis, we repeated these results with the fluorescence traces and found that the same set of results holds (Figure S21).

We also note that, in practice, EMD quantiles are superior to EMD distances and enable for comparisons across animals, contexts or other groups where behavioral effects can influence raw EMD values. We therefore repeat this analysis using the single point EMD with two sets of reference quantiles (Figure 9). In the first, we demonstrate quantiles using across cell reference distributions. That is, for a given cell map to reference map distance, the quantile describes how many of the other cell distances are larger. This would be suitable for a simple threshold technique (e.g. top10% = *q<* 0.1). This however describes across cell references for a within cell analysis (relative to a reference map). If stability were being assessed across cells (e.g. from one session to the next), an across cell reference distribution would be more suitable if made up of remapping distances computed on the incorrect cell-pairings. Such a reference distribution would be more flexible, and could be separated based on hierarchal data. For example distributions can be computed for mismatched distances at the level of animals, sessions, contexts or other.

Such reference groupings are not seen for within cell reference quantiles. We provide such an example using random sampling of locations within the firing map (Figure 9). In doing so, we create a spatial reference distribution consisting of distances relative to randomly smapled positions. While simple in 1D maps, sampling techniques can vary for 2D maps. Given how the distribution of EMD distances across a map looks (Figure 1, Figure 2, Figure 4), it is appropriate to attempt an even sample across the map. To do this we can use approaches such as rectangular sampling for unmasked, square ratemaps or hexagonal sampling for masked, circular ratemaps.

In our example, while based on different counterfactuals and analyses, the different choices of across or within cell reference distributions provide similar results. This is likely owing to the simple task design however in practice, the choice of reference distributions is important and can vary given different hypotheses. It is also important to note the difference in raw quantile values. In the within cell reference, quantiles are generally larger. This is something also observed in 2D maps and can be explained by the symmetrical, distributional approach of the EMD, as well as the emergence of multiple fields or areas of noise. To reduce this, we can sample locations at a certain distance from our reference point. Additionally, in practice, it is often helpful to extract the location of different fields and compute these metrics separately for different fields on a map. The comparison of whole map to field metrics is also informative and can explain observed quantile distributions. Therefore we show how, in practice, cell references can be computed within and across cells depending on the use case, expected counterfactual and other topic-specific knowledge.

While we acknowledge that previous studies have used Pearson’s correlation coefficient to evaluate remapping using a reference template (Masuda et al. (2023)), or a set of gaussian templates (Nagelhus et al. (2023)), our results with synthetic and real data demonstrate that the EMD is more robust to such template choices. For example, in one study, a reference template computed from the activity of cells in the baseline context was used to compute similarity (Masuda et al. (2023)). However with the bin to bin correlation approach, such a template could vary in results given different parameters that shift, bin or smooth the correlated bins. In another study with 2D ratemaps, this required using multiple sets of gaussian templates spanning a range of hyperparameters (Nagelhus et al. (2023)). The distributional focus of the EMD and the ability to describe dissimilarity regardless of size, dimensions and binning demonstrates its superiority to Pearson’s r in practice. This is especially true for noise-heavy, multi-field and different-sized ratemaps that are either 1D or 2D and applies to within cell dissimilarity and reference relative dissimilarity.

## 4 DISCUSSION

Through these simulated and recorded cases of remapping, we demonstrate that the Earth Mover’s Distance (EMD) is more spatially sensitive in characterizing remapping than Pearson’s r correlation coefficient and other plausible metrics like the non-linear Spearman rank correlation coefficient. We find that both Pearson’s r and EMD are suitable for cases of remapping where fields are still overlapping from session to session. However, we demonstrate that Pearson’s r is unsuitable in describing remapping that results in non-overlapping receptive fields whereas EMD can numerically quantify remapping at any point in the ratemap. This EMD property is especially useful in experimental setups where arena shape is varied as they result in specific map areas, where a cell can reasonably move to, that cannot be appropriately quantified with Pearson’s r. For example, consider a field that remaps to the corner of a square arena after having been in a circular arena with diameter such that it is inscribed within the square. This field will have 0 correlation when tested on the circle arena followed by the square arena because of non-overlap at the 4 corners despite the fact that transformations to such regions are non-identical and can be distinguished from each other using the EMD. We show that EMD holds its sensitivity for circular/elliptical place fields and wide/narrow spaced grid fields. Additionally, we find that the symmetry offered by EMD allows us to describe more complex non-linear translations such as rotations and scaling of fields. In fact, we find that Pearson’s r is less suitable than the EMD in describing either scaling or rotations and does not offer the same sensitivity that the EMD provides. More importantly however, we show that this spatial sensitivity of the EMD to linear and non-linear transformations is robust to noise and field degeneration. By manipulating fixed fields across a range of standard deviations of added noise, we find that, for both normalized and unnormalized ratemaps, the EMD is more stable relative to the ‘true’ EMD score. That is, as we increase the standard deviation of noise, EMD varies less frequently and at later standard deviations than the Pearson’s r score. Notably, we see that the EMD performs similarly in the normalized and unnormalized cases also demonstrating a robustness to rate. While the raw EMD values change for the normalized and unnormalized ratemaps, the distribution across standard deviations evolves in the same ways, with fewer outliers in the normalized and smoothed-then-normalized cases than the unnormalized case, further demonstrating a type of robustness to rate. The application of a binary EMD where all spikes are considered with even weight by imputing 0 and 1 for outside and inside the field respectively further disentangles rate effects by exclusively describing the underlying dispersion of the field or map, regardless of firing rate. This highlights the use of such a metric in describing field or whole map distortions where the spatial distribution of spikes can change separately to rate and even the field centroid.

Such dispersion metrics can be thought of as purely spatial and can be extended to include the entire distribution of raw spike positions as part of a spike density EMD. This ‘pure spatial’ metric may be most useful in the case of linear tracks with identical navigation structure across animals but can also be extended to arena navigation studies where coverage and occupancy vary from animal to animal. In the latter, caution should be taken to account for coverage where arena sizes may be consistent but map exploration varies from animal to animal. Normalization of distances by coverage limits would reduce such effects. Additionally, care should be taken in interpreting dispersion metrics (binary and spike density) in the case of extremely biased occupancy where most of the spiking distribution will fall in the same area purely due to biased behavior. This argument also extends to the use of EMD to assess temporal similarity profiles as part of a ‘pure temporal’ metric (Grossberger et al. (2018), Sihn and Kim (2019)). These pure metrics may be best interpreted in such a way where especially stable distances (binary/density or temporal EMD) along with fair occupancy, and while accounting for coverage limits, would be a non-trivial result, as would differing trends among the rate, temporal and spatial components of remapping. In fact, given that rate remapping is well defined and temporal remapping of firing profiles can be quantified (Sotomayor-Gómez et al. (2023)), we further suggest the EMD and single point Wasserstein metric to describe the spatial component in firing rate maps enabling a much more detailed and flexible framework for remapping. Through various manipulations, this would allow for an understanding of how different components of neural coding interact by separately and concurrently considering the rate, spatial and temporal components.

We also see in these examples that the whole map EMD is more suitable than the field-restricted EMD in capturing remapping despite degeneration. The reason behind this is likely two fold. The first being that the field-restricted EMD does not capture noise across the entire spatial map. Since, in our example, randomly sampled normally distributed values are added at every position in the ratemap (NxN noise), the specific subset added to the indices of a given field may be particularly large/small relative to the rest of the distribution. As such, from step to step (each increase in std dev), restricting the EMD metric to the single field can cause larger deviations than would be seen with the whole map case. The second reason lies in the methods for detection of gaussian fields and the sensitivity of these approaches to noise and spatial degeneration. The best approach is often to select a contiguous region from a given spatial map, with a minimum size, where the firing rate is above a certain peak threshold (Fyhn et al. (2007), et al. (2021b)). Although this is suitable in the cases of low noise, degenerate and noisy spatial maps are often experimentally recorded and need more granular characterization of place field centroids and area. Traditional field characterization methods are sensitive to noise and often involve smoothing of the ratemap as part of this procedure (Fyhn et al. (2007), Grijseels et al. (2021b)). With minimal smoothing, localization of place fields in a noisy map results in multiple ‘noise’ fields detected with centroids deviating from the true field positions (Figure S9, Figure S10). With greater smoothing, localization of place fields on a noisy map can result in too wide an area being characterized as a field and multiple fields being incorrectly merged (Figure S9, Figure S10). In the presence of similar noise throughout the rate map, smoothing can even result in a contiguous region crossing detection thresholds despite being outside the fields of interest (Figure S9, Figure S10). Therefore the field EMD can be particularly susceptible to the accuracy of the detected fields and by extension to the underlying field detection process. However both the whole map EMD and field EMD still offer more stable and robust alternatives to Pearson’s r, especially in cases of high noise and for disease states.

While we find that the whole map to point map EMD does not replace the map blobs (fields) detection approach, we show that it can help inform classification of valid blobs by highlighting the region of lowest remapping through a gradient of remapping values. In the case of known single field cells (e.g. place cells), the map to point EMD will identify the region of lowest remapping which falls near the true field centroid and can therefore be used to filter out extracted noise blobs that may have crossed the area/size threshold but fall far from the region of lowest remapping. In the case of dual fields, the EMD can only separately identify fields at low noise levels and using the top 90% threshold as opposed to the top 20% use for Pearson’s r or 20% peak firing rates. In fact, in the case of dual fields, the EMD gravitates to the center point between the two fields. This is the point that requires the least amount of remapping. While it cannot be used for field detection in this case, it may yet be used to filter out blobs that fall far from this region. Therefore the whole map to point map EMD approach is suitable for localizing in the case of a known single field (e.g place cell) or to identify the region of lowest remapping in the case of multiple fields. In fact, the EMD is not as susceptible to noise as Pearson’s r and peak firing rates in that it performs more consistently and with less degradation in either the case of unnormalized ratemaps or smoothed then normalized ratemaps.

Moreover, the remapping quantiles that are generated across the entire map for single and multi-field cases can be used as a standard reference distribution for a given cell. That is, single point EMD (Wasserstein) scores can be converted into a quantile below which all values are smaller (easier point locations to remap to) and above which all values are larger (harder point locations to remap to). We see that these quantiles are a result of computing remapping scores between the whole map and every possible point on the rate map (Figure 4). In doing so, single point EMD values, and by extension individual fields, can be localized on a rate map through the ‘region’ of lowest remapping that they fall in. These quantiles may also enable comparisons across different cells where raw EMD values cannot be directly compared. For example, in practice, while arena size may be consistent, coverage and occupancy can vary, as can firing rates as a result of biased exploration. Despite the rate robustness of the EMD, distances can vary on different scales due to different behavior profiles. In such a scenario, one might observe a comparable trend in remapping within a cell’s own sessions, and even across the wider population where all cells increase or decrease EMD values across sessions. Yet, the raw EMD scores may not be comparable. For example, two cells, each from a different animal, could both be near a specific map-region but with very different raw distances required to move there because of exploration. Even with consistent coverage of arenas, several experimental contexts make use of changing arena shapes thus changing the total distance available in the EMD computation and creating the same situation for cells from the same animal across different sessions. Providing a quantile alongside a remapping value allows for a standard scale to describe how far/close the computed remapping score is relative to all the possible points of remapping in a rate map. If a position required the least remapping (e.g. exactly at the object location for an object cell or at the centroid of an individual field), then the quantile would be sufficiently small such that most or all of the other possible locations result in a larger remapping score. These quantiles are unique to the EMD and can be computed across noise standard deviations allowing for a quantification of remapping regions despite degeneration, instability and substantial behavioral/rate influences. These quantiles can also be used to describe spatial remapping when raw distances cannot be directly interpreted.

The EMD’s viability as a remapping metric is bolstered by the computational expediency of its simplified cases (Figure S11). The sliced EMD approximation can adequately estimate the true EMD value. Using the sliced EMD across varying numbers of projections, with 10**2 offering a favorable balance of speed and accuracy, renders the EMD an optimal choice for cell remapping analyses. Nonetheless, the sliced EMD may still be computationally demanding in instances involving exceptionally large rate maps (e.g., 256 x 256) and a substantial number of cells. The modified map to single point approach also allows for swift and adaptable application of the EMD. This is particularly important in describing stability relative to specific locations. While we acknowledge that previous studies have used Pearson’s correlation coefficient to evaluate remapping using a reference template (Masuda et al. (2023)), or a set of gaussian templates (Nagelhus et al. (2023)), our results with synthetic and real data demonstrate that the EMD is more robust to such template choices. For example, in one study, a reference template computed from the activity of cells in the baseline context was used to compute similarity (Masuda et al. (2023)). However with the bin to bin correlation approach, such a template could vary in results given different parameters that shift, bin or smooth the correlated bins. In another study with 2D ratemaps, this required using multiple sets of gaussian templates spanning a range of hyperparameters (Nagelhus et al. (2023)). Therefore, with the linear runtime of both the single point EMD approach and Pearson’s r, it can be both more efficient and simpler to use the EMD metric. Avoiding such choices is possible because of the distributional focus of the EMD and its ability to describe dissimilarity regardless of size, dimensions and binning.

The EMD on 2-dimensional maps is better equipped to describe the spatiotemporal patterns seen than Pearon’s r correlation coefficient. However, the temporal aspect of these maps could be further characterized by applying the EMD in a stepwise manner. In a basic example, one could take the spike train of a given cell and iterate over time windows of activity to produce a continuous string of EMD values describing remapping as it evolves over an experimental session. While this approach is highly unsuitable with Pearson’s r because of the need for same size distributions, and the over-susceptibility to spurious correlations at lower sample sizes, EMD has been shown in this paper to be more robust, and in previous work to support continuous application (Zhao et al. (2010)). Given that the EMD can be applied on raw spike positions (binary EMD) and does not require a rate map to describe the pure spatial and pure temporal components, we further propose it as a flexible tool for disentangling spatiotemporal components in a continuous-like setup. Such a setup could be used to describe remapping as it relates to specific temporal events/markers. With sufficiently long experimental sessions and high sample rates, EMD will support a characterization of remapping on smaller timescales than separate sessions and can enlighten intersession and intertrial remapping dynamics that are triggered or otherwise shaped by time. For example, one can consider an experimental setup where object location is rotated continuously or otherwise transformed during the session, as was seen in previous work (Shapiro et al. (1997), Knierim et al. (1995)). In such a case, applying EMD across a sliding window of spike activity can be highly informative in identifying how remapping evolves over time for a given cell and for the broader population. This can be especially useful given the varying results seen in morph experiments where partial and complete remapping are seen to occur in different ratios across different studies (Wills et al. (2005), Leutgeb et al. (2005a), Colgin et al. (2010)). Using EMD we can better describe the quantity of remapping over time, the periods or triggers before/after which the gradient of remapping increases or decreases, and the amount of remapping across the population of cells (or ensembles of cells) at specific points in time.

In summary, the Earth Mover’s Distance (EMD) offers a more comprehensive and spatially sensitive approach to characterizing remapping in comparison to the Pearson’s r correlation coefficient. EMD’s ability to handle non-overlapping receptive fields and intricate non-linear transformations, such as rotations and scaling, renders it a powerful tool for understanding the complexities of spatial navigation and remapping. Although Pearson’s r might remain useful in specific cases with linear relationships and overlapping fields, EMD’s versatility makes it applicable to a broader range of scenarios. EMD estimators such as sliced EMD are computationally expensive with respect to correlation metrics. However, most modern computers can easily handle the additional computational load and this should not be a hindrance to the adoption of EMD in most use-cases. The application of EMD in spatial remapping research has far-reaching implications in the study of memory and neurodegenerative disorders, such as Alzheimer’s Disease (Jun et al. (2020), Ridler et al. (2020), Fu et al. (2017)). By providing a more detailed analysis of spatial remapping, EMD can shed light on the intricate relationships between spatial representations, memory formation, and the influence of various factors on these processes. The enhanced understanding of remapping dynamics facilitated by EMD may contribute to the identification of potential therapeutic targets for memory-related disorders, thereby opening new avenues for Alzheimer’s Disease research and treatment.

## Supporting information

Supplementary Figures (Spearman)

## AUTHOR CONTRIBUTIONS

A.A., O.S., and S.A.H. conceptualized the project. O.S. proposed the theory and use of optimal transport metrics, and defined the statistical methods. R.R. and G.A.R. curated and provided all real data examples. A.A. designed the experiments with guidance from O.S. and S.A.H., implemented the simulations and conducted the data analysis. A.A. crafted all figures and wrote the primary manuscript. O.S. wrote the methods. The document was reviewed and edited by A.A., O.S., R.R., G.A.R. and S.A.H.

## ACKNOWLEDGMENTS

We would like to thank the Hussaini Lab for their advice and assistance in developing this framework and for their helpful comments throughout the writing of this manuscript. We also thank the reviewers for their helpful critiques and feedback towards strengthening this work.

## DATA AVAILABILITY STATEMENT

The jupyter notebooks used to synthesize data and generate the figures can be found at the Hussaini Lab github page in the Neuroscikit project under prototypes/cell remapping /remapping paper.

