## Supplementary Data & Figures (I) for "Beyond Correlation: Optimal Transport Metrics For Characterizing Representational Stability and Remapping in Neurons Encoding Spatial Memory"

### Supplementary Material

#### 1 SUPPLEMENTARY DATA

Additional examples of synthesized field data pre-interpolation. Data are generated using basic place field manipulations to vary location, size and shape. Examples of corner/directional fields, field scaling and field tracking using EMD and centroid distances for the 4 cardinal directions and angles.

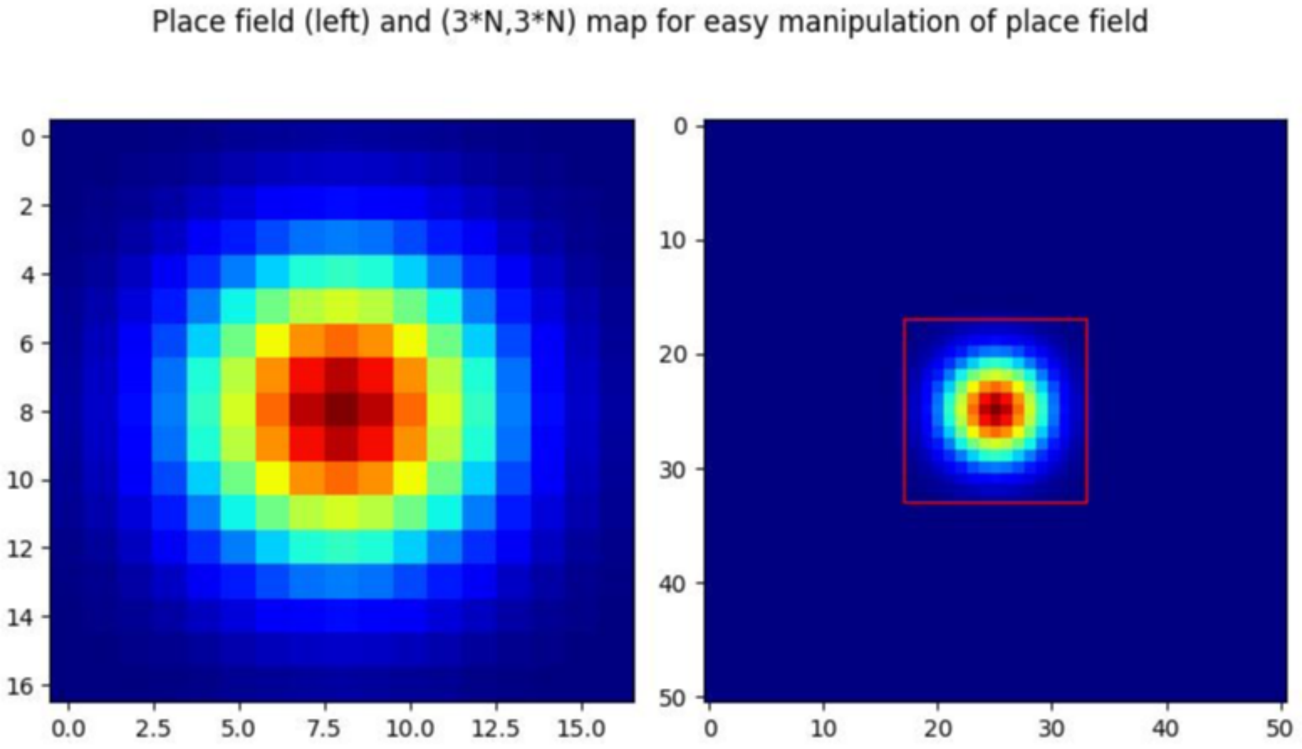


**Figure S1. Synthetic gaussian field.** Example of a (17,17) gaussian field with $\sigma = 3$ without interpolation to (257,257) (left). A wider (3*17,3*17) map is shown which allows for shifting of the gaussian field by its centroid (right). The red square denotes all possible locations a centroid can move to.

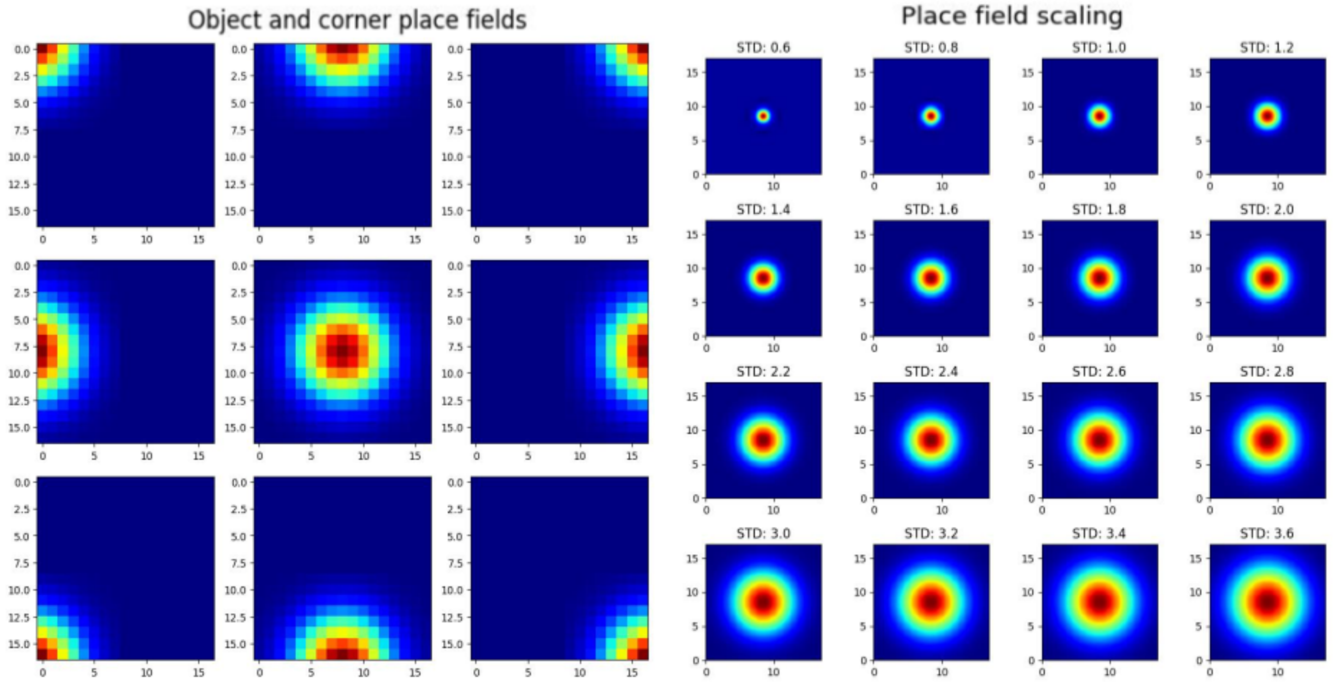


**Figure S2. Non-linear field transformations.** Corner/boundary fields are shown where consecutive 90 degree transformations create 4 rotated positions for each field in the square arena (left). Example of scaling by varying field $\sigma$ from 0.6 to 3.6 (right).

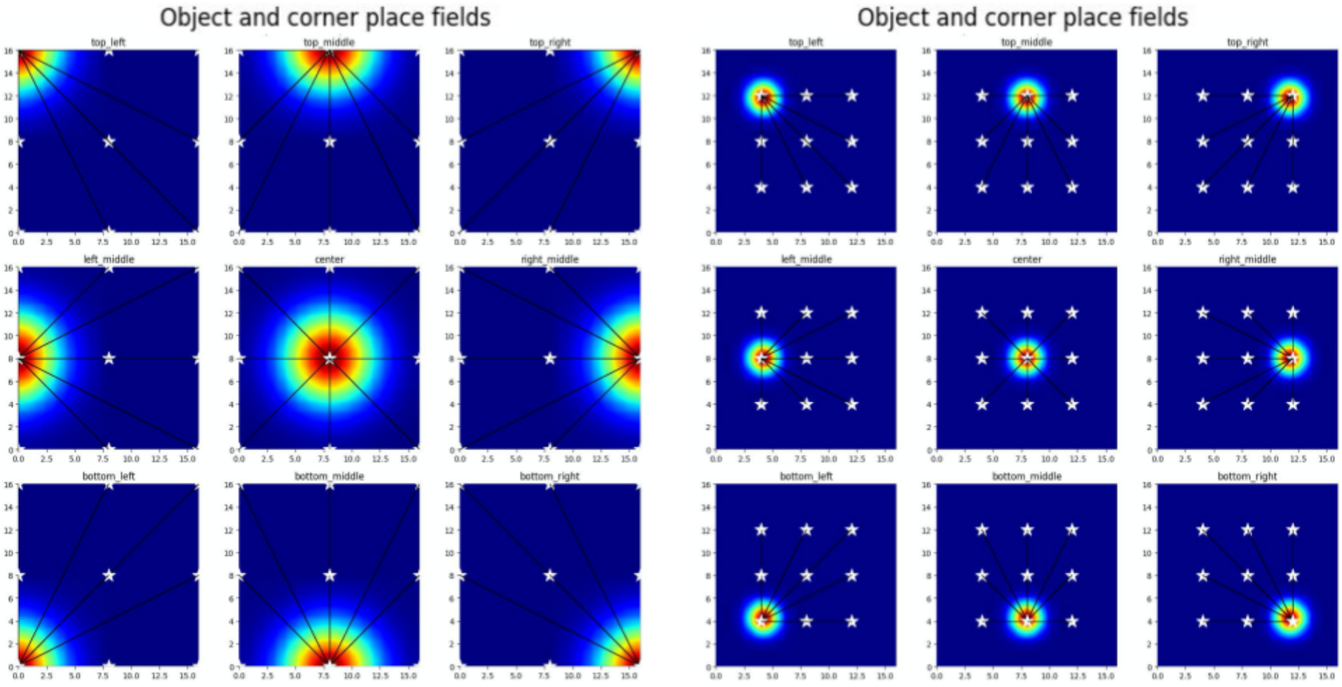


**Figure S3. Object/Trace transformations.** Example of fields to be used for rotational remapping and object/trace mechanisms. Plots show fields at the 4 corners, the 4 cardinal directions (N,S,W,E) and the center. Object locations are denoted by white stars and connected to the field centroid with a black line (centroid distance).

#### 2 SUPPLEMENTARY TABLES AND FIGURES

##### 2.1 Figures

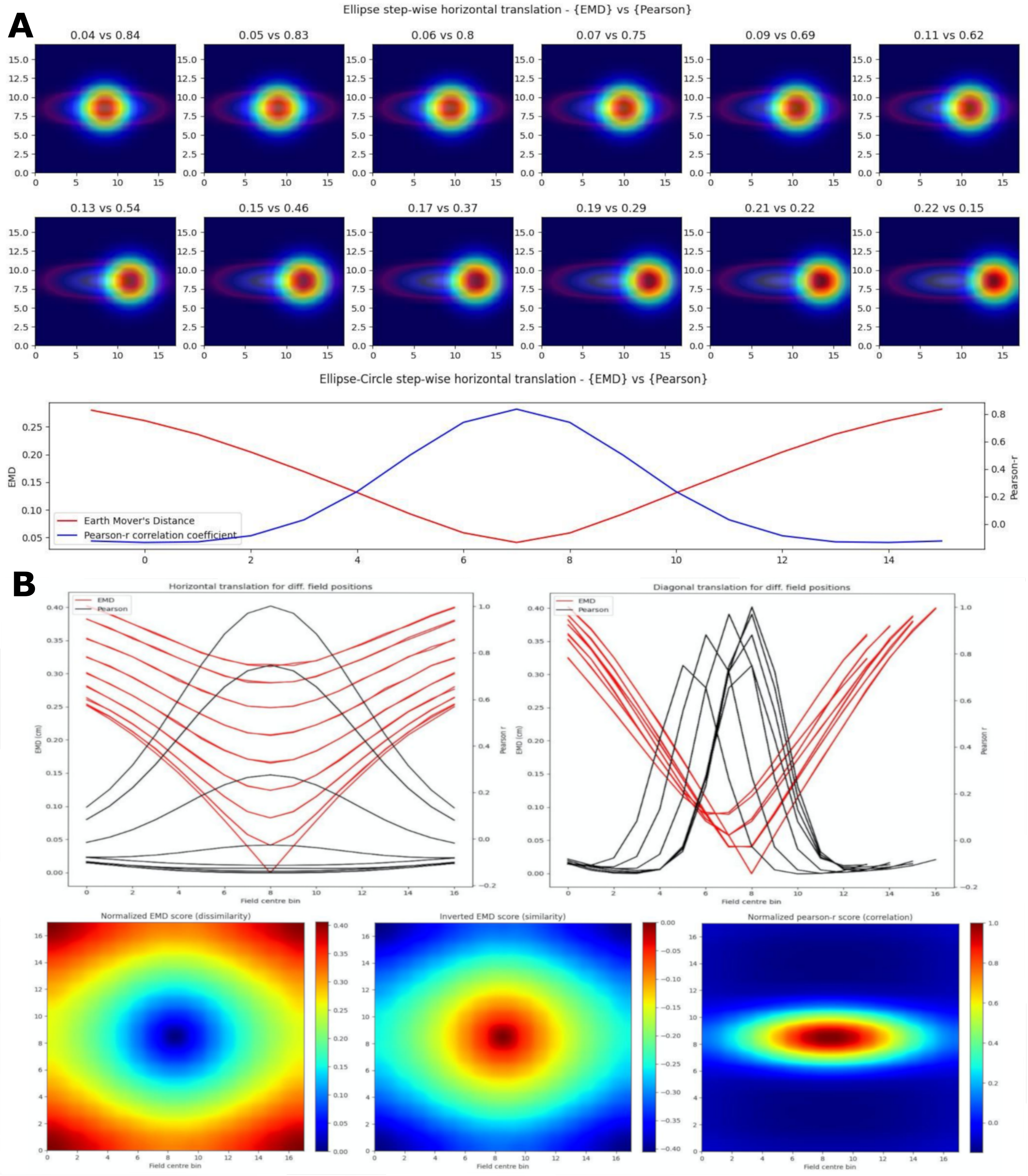

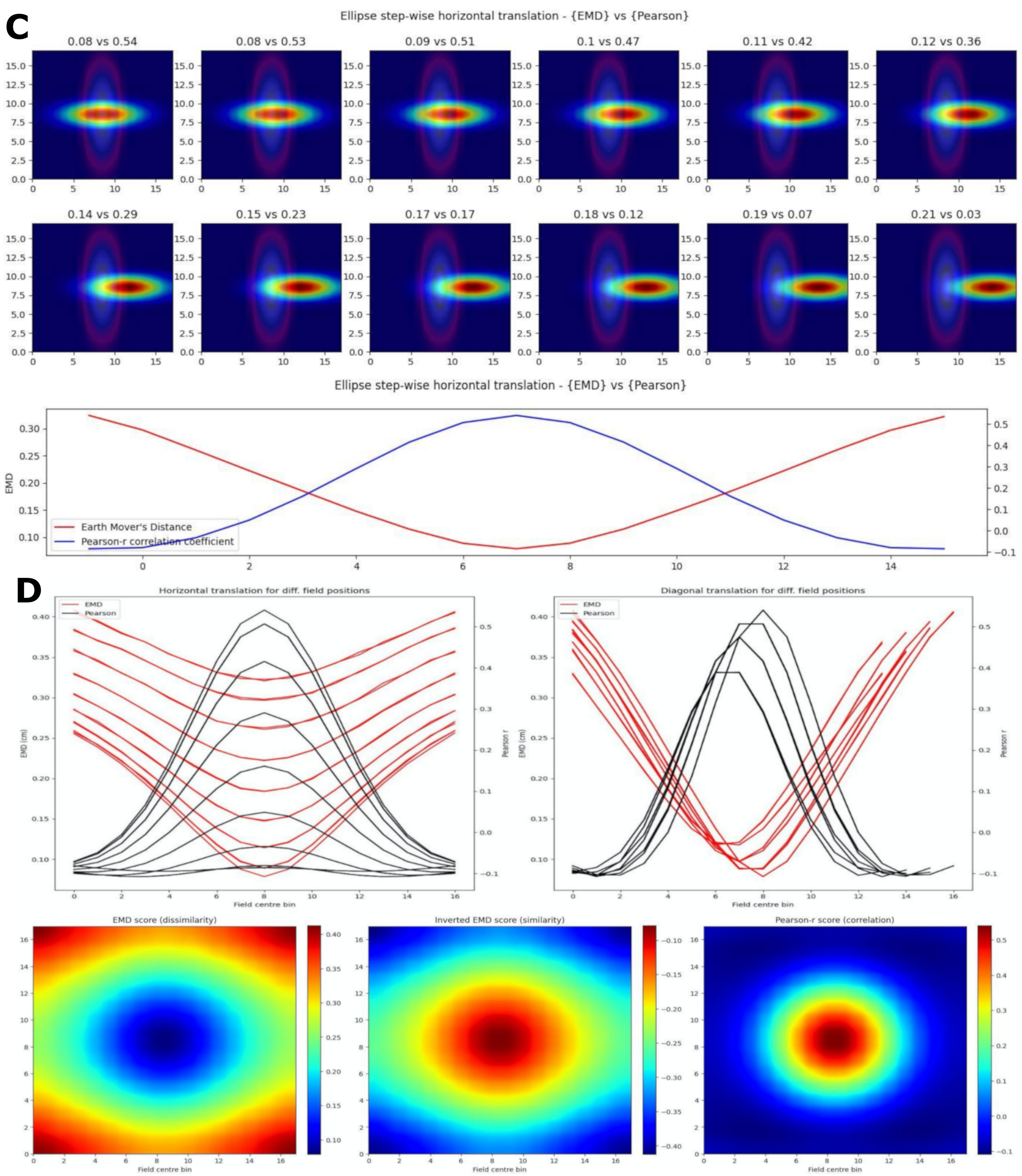


**Figure S4. Non-identical place field translation**. Stepwise horizontal linear translation of non-identical, overlapping place fields ($N = 17$, $\sigma = 3$) moving from the center to the right (**A, C**). EMD score is shown on the left while Pearson's r is shown on the right. 12 steps are shown and scores are rounded for display (top panel). Scores from remapping tested at all possible centroids in a single row on the rate map (bottom panel). EMD and Pearson's r scores tested at all possible centroids in the rate map (N*N) (**B, D**). Horizontal and diagonal translations across the rate map are shown for all rows ($N = 17$) (top panel). Heatmap showing the gradient of EMD scores both raw and inverted to match Pearson's r color scheme (bottom panel).

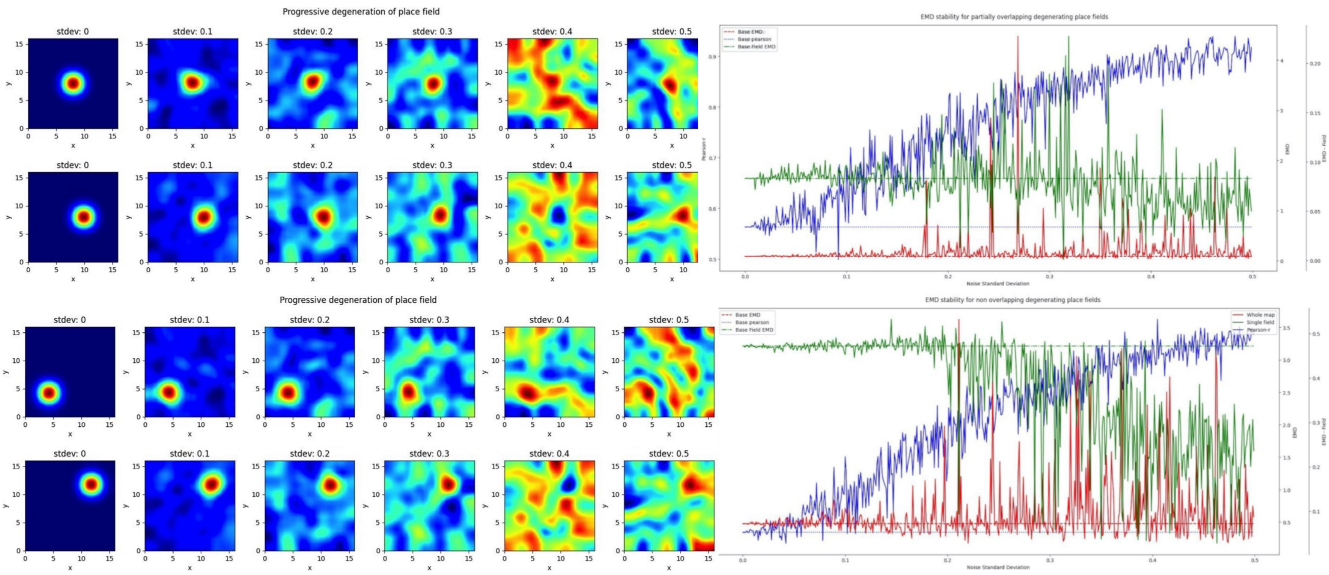


**Figure S5.** Repetition of Figure 3 but for normalization and smoothing post-added noise. The same noise distribution that was added to the unnormalized case is added at each step. Examples are shown for overlapping (top panel) and non-overlapping fields (bottom panel).

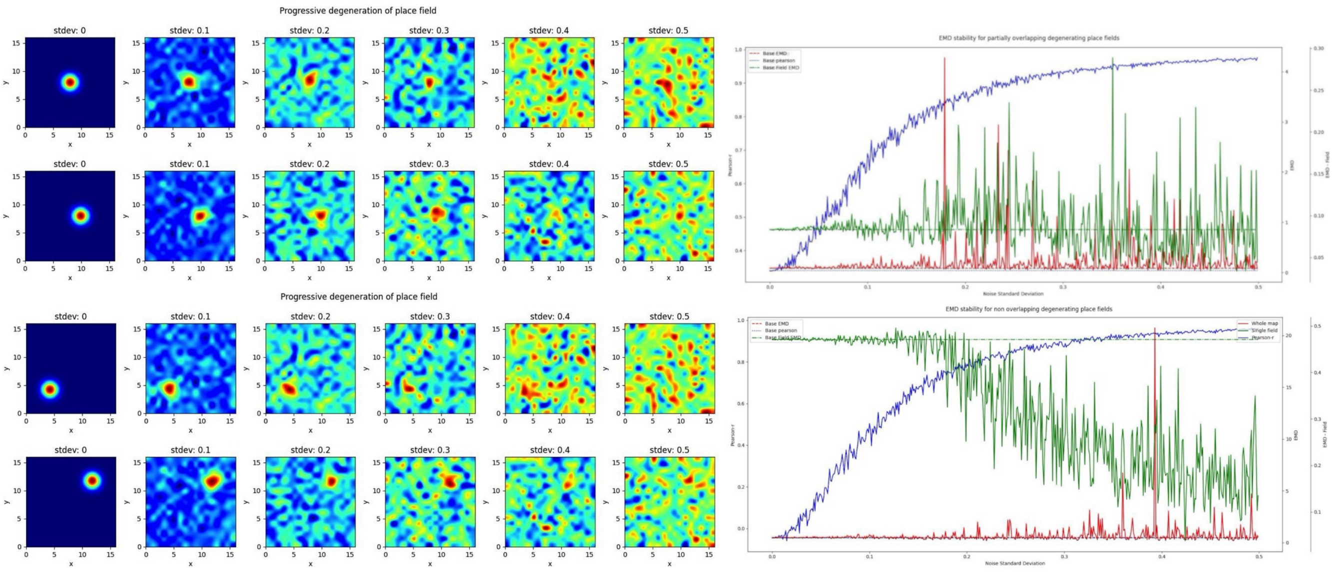


**Figure S6.** Repetition of Figure 3 but for normalization only post-added noise. The same noise distribution that was added to the unnormalized case is added at each step. Examples are shown for overlapping (top panel) and non-overlapping fields (bottom panel).

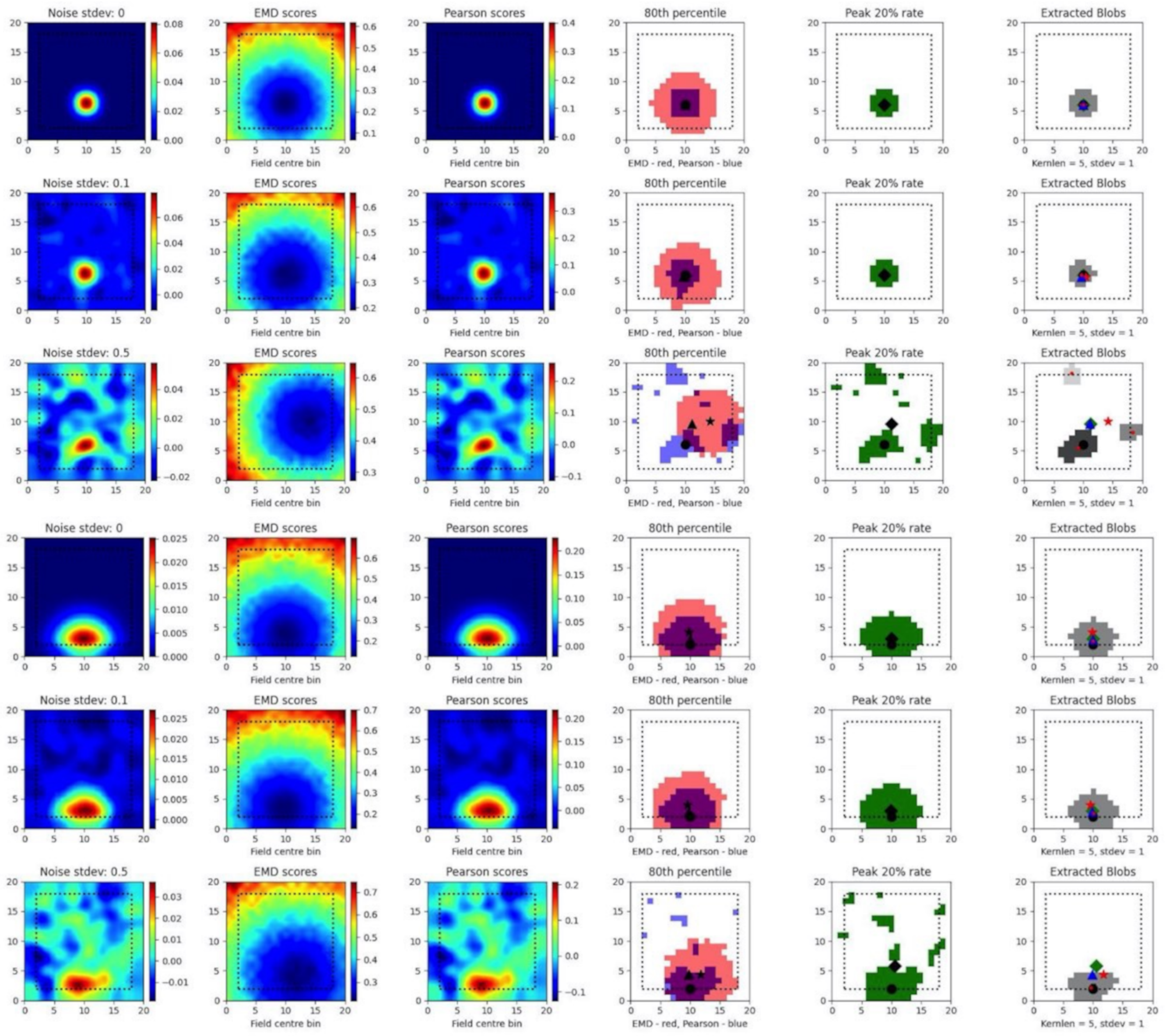


**Figure S7. Single field localization**. Additional field localization plots from Figure 4). Here, localization plots are shown across three different noise levels (rows: no noise 0, low noise 0.1 and high noise 0.5)

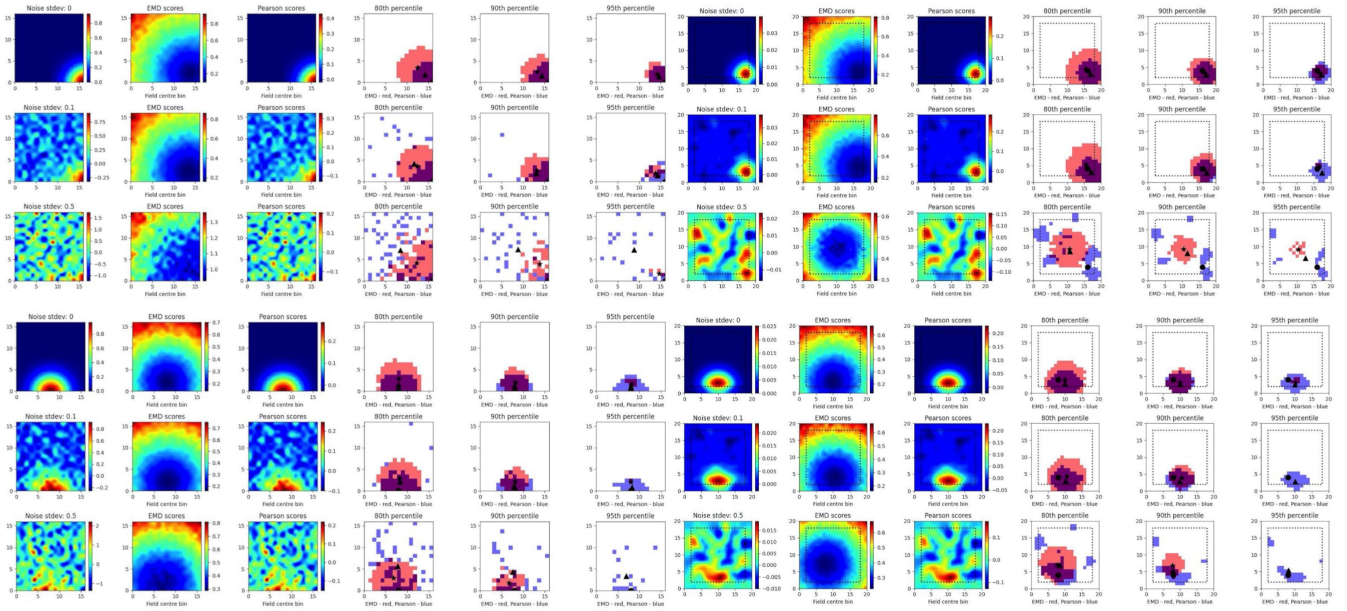


**Figure S8. Localization with padding and smoothing**. These plots come in pairs where the first 3 rows (no noise 0, low noise 0.1 and high noise 0.5) are the unnormalized, unsmoothed and unpadded version of Fig4 while the next 3 rows show the recovery of scores (in some cases) post padding, normalizing and smoothing. The last 3 columns are for the specificity of scores and show the 80th, 90th and 95th percentile of EMD and Pearson's r scores.
