## Supplementary Data & Figures (II) for "Beyond Correlation: Optimal Transport Metrics For Characterizing Representational Stability and Remapping in Neurons Encoding Spatial Memory"

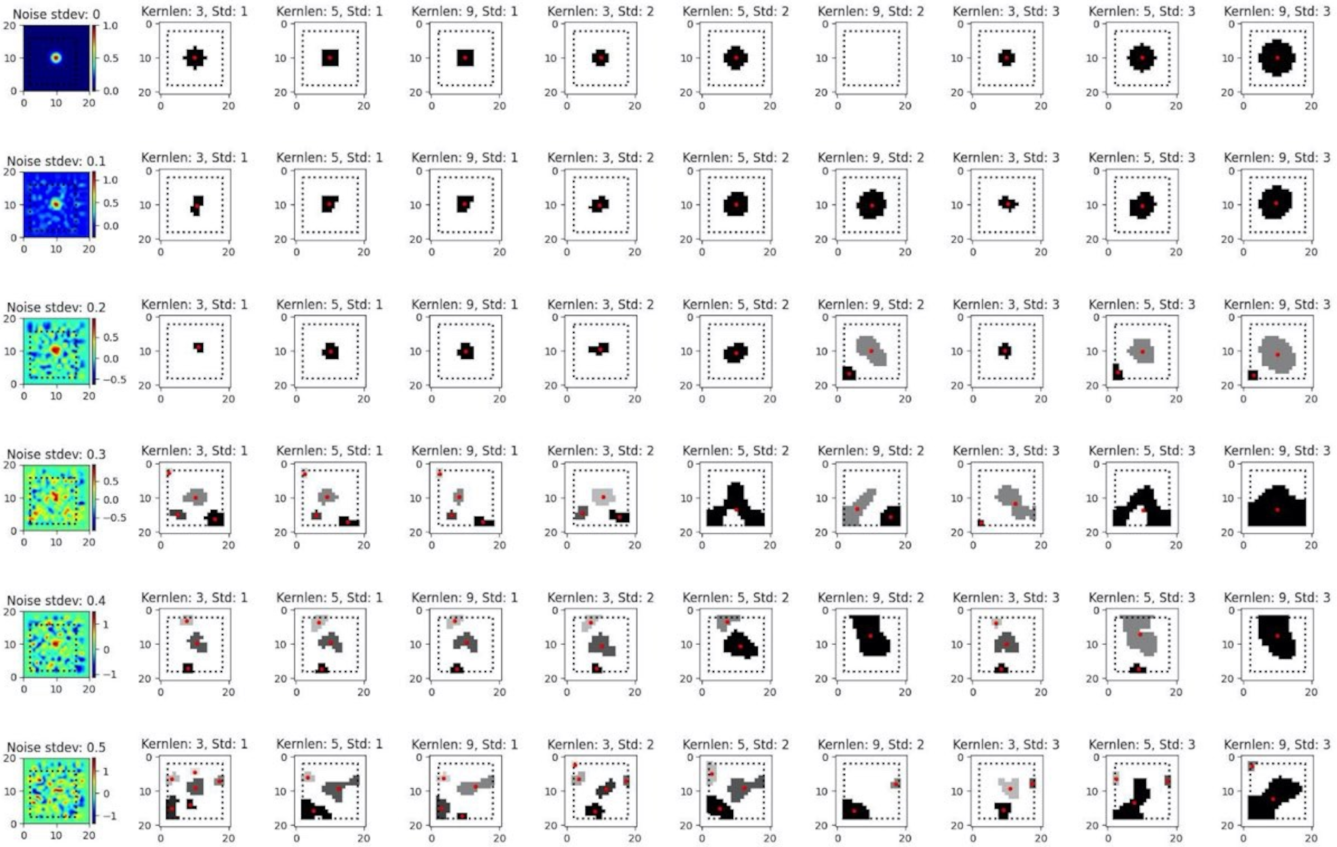


**Figure S9. Field extraction susceptibility to noise and smoothing - Single field**. Field extraction algorithm tested across six noise levels ($\sigma = 0, 0.1, 0.2, 0.3, 0.4, 0.5$), 3 kernel sizes (3,5,9) and 3 kernel standard deviations (1,2,3) with resulting extracted blobs shown along with centroids (red dots). The first column shows the ratemap immediately post added noise with no additional processing.

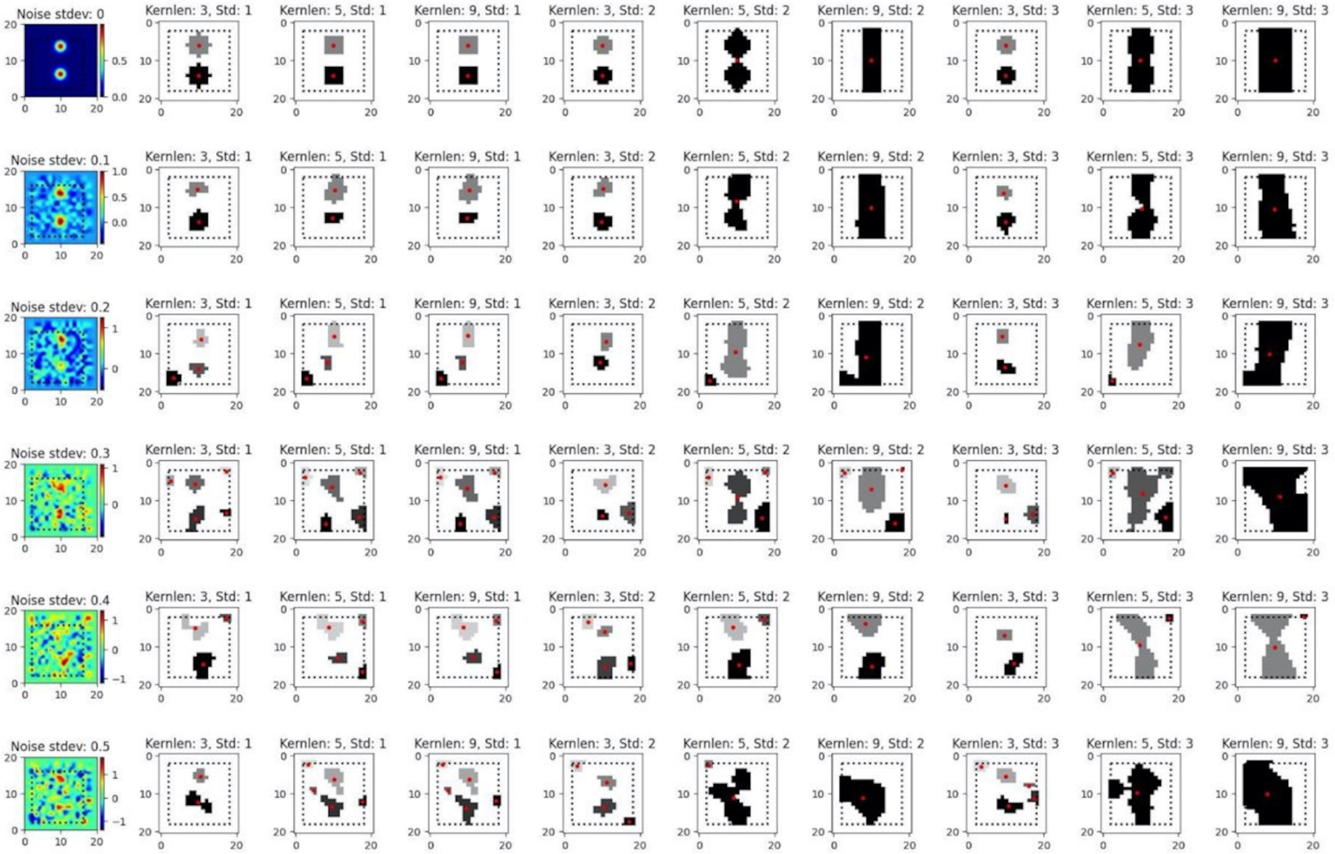


**Figure S10. Field extraction susceptibility to noise and smoothing - Dual field**. Field extraction algorithm tested across six noise levels ($\sigma = 0, 0.1, 0.2, 0.3, 0.4, 0.5$), 3 kernel sizes (3,5,9) and 3 kernel standard deviations (1,2,3) with resulting extracted blobs shown along with centroids (red dots). The first column shows the ratemap immediately post added noise with no additional processing.

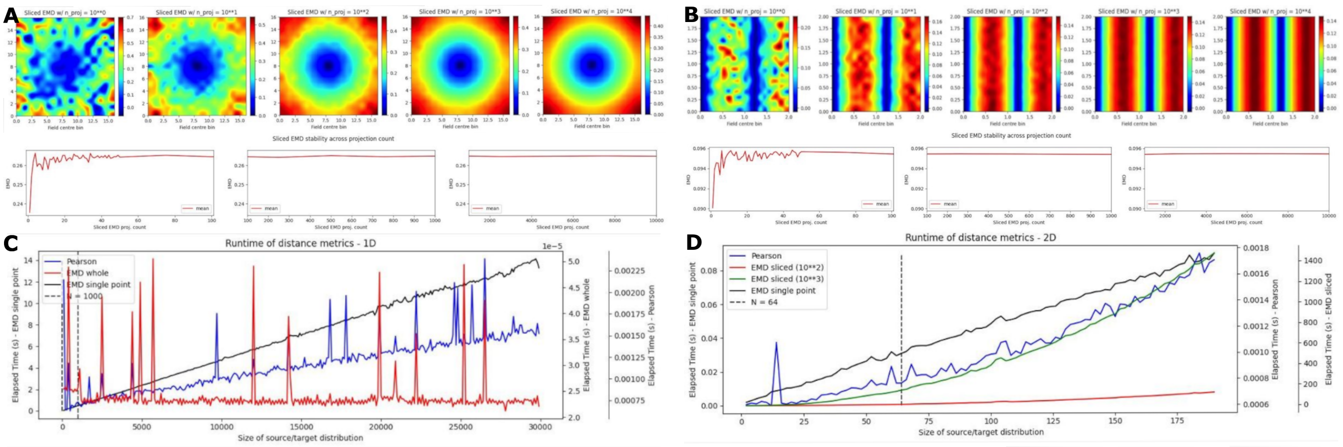


| **Optimal Transport Metric** | **Cost 1D Map** | **Cost 2D Map** |
|---|---|---|
| Whole Map EMD | $O(n^3)$ | $O((mn)^3)$ |
| Sliced EMD | $N/A$ | $O(Ln\log(n))$ |
| Single Point Wasserstein | $O(n)$ | $O(n)$ |
| Spearman Rank | $O(n\log(n))$ | $O(n\log(n))$ |
| Pearson Correlation | $O(n)$ | $O(n)$ |

**Figure S11. Numerical Cost**. Repetition of heatmap procedure for place fields in Figure 1 across different numbers of projections using the sliced EMD (**A**). Default settings is at 10**2 after which values remain notably more stable. Heatmaps are shown for 10**(0,1,2,3,4) projections with distribution plots across incremental steps shown below. Repetition of heatmap procedure for grid fields in Figure 2 across different numbers of projections using the sliced EMD (**B**). Example of how compute time evolves using remapping metrics on a 1D map (**C**). Example of how compute time evolves using remapping metrics on a 2D map (**D**). Table detailing numerical cost of different remapping metrics using big O notation. Cost is shown for 1 dimension and 2 dimensional acitvity maps.

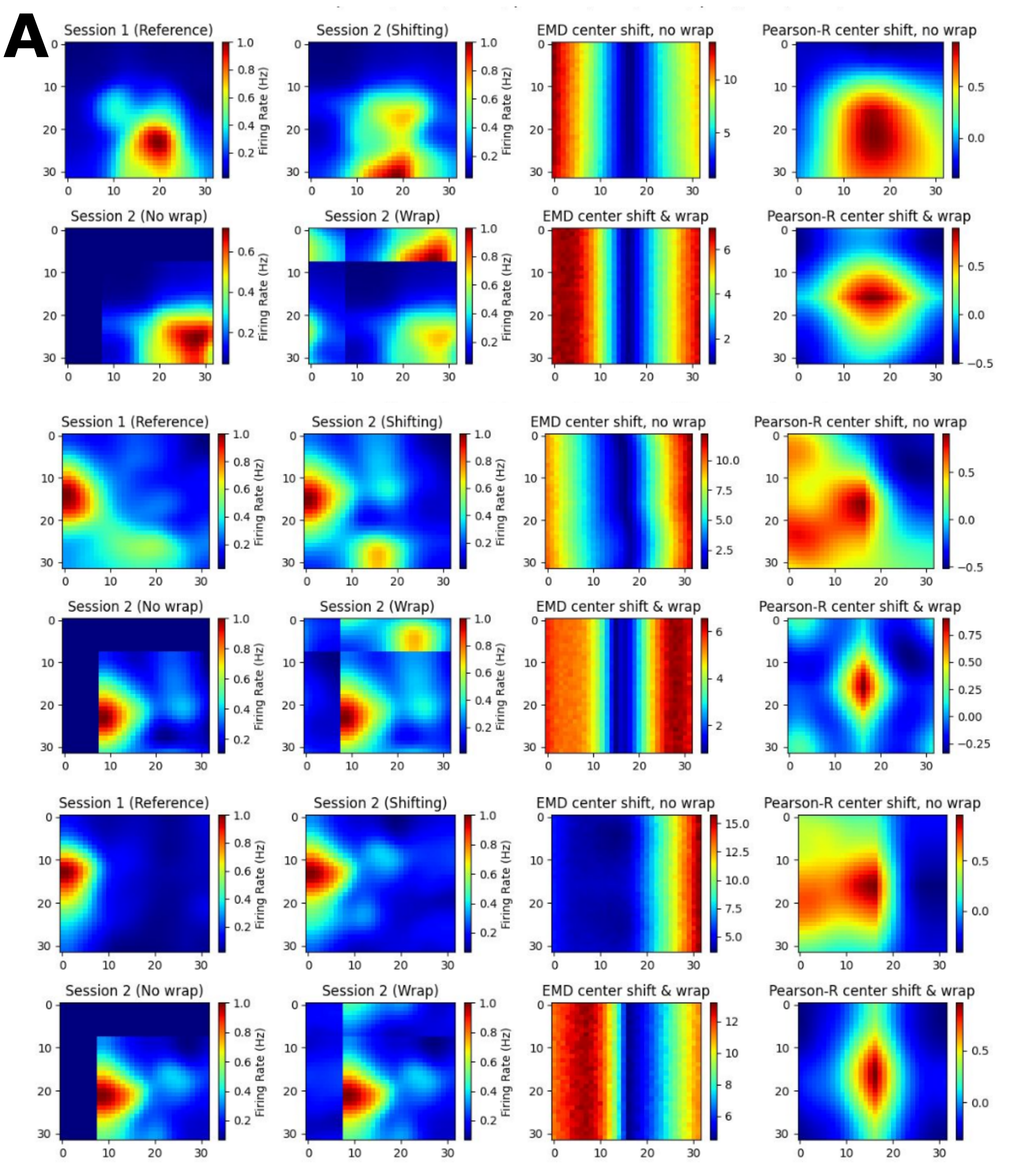


**Figure S12. Additional MEC cell examples**. Additional examples of ratemaps from the MEC of AD mouse models for a reference session and a shifting session. Gradient of Pearson's r scores tested at all possible map shift centers (N*N) is shown to the far right of each cell example with the EMD gradient immediately to the left of it. For each MEC example, the top row demonstrates a shift with no wrap (0 padding) while the bottom row demonstrates a shift with wrapping.

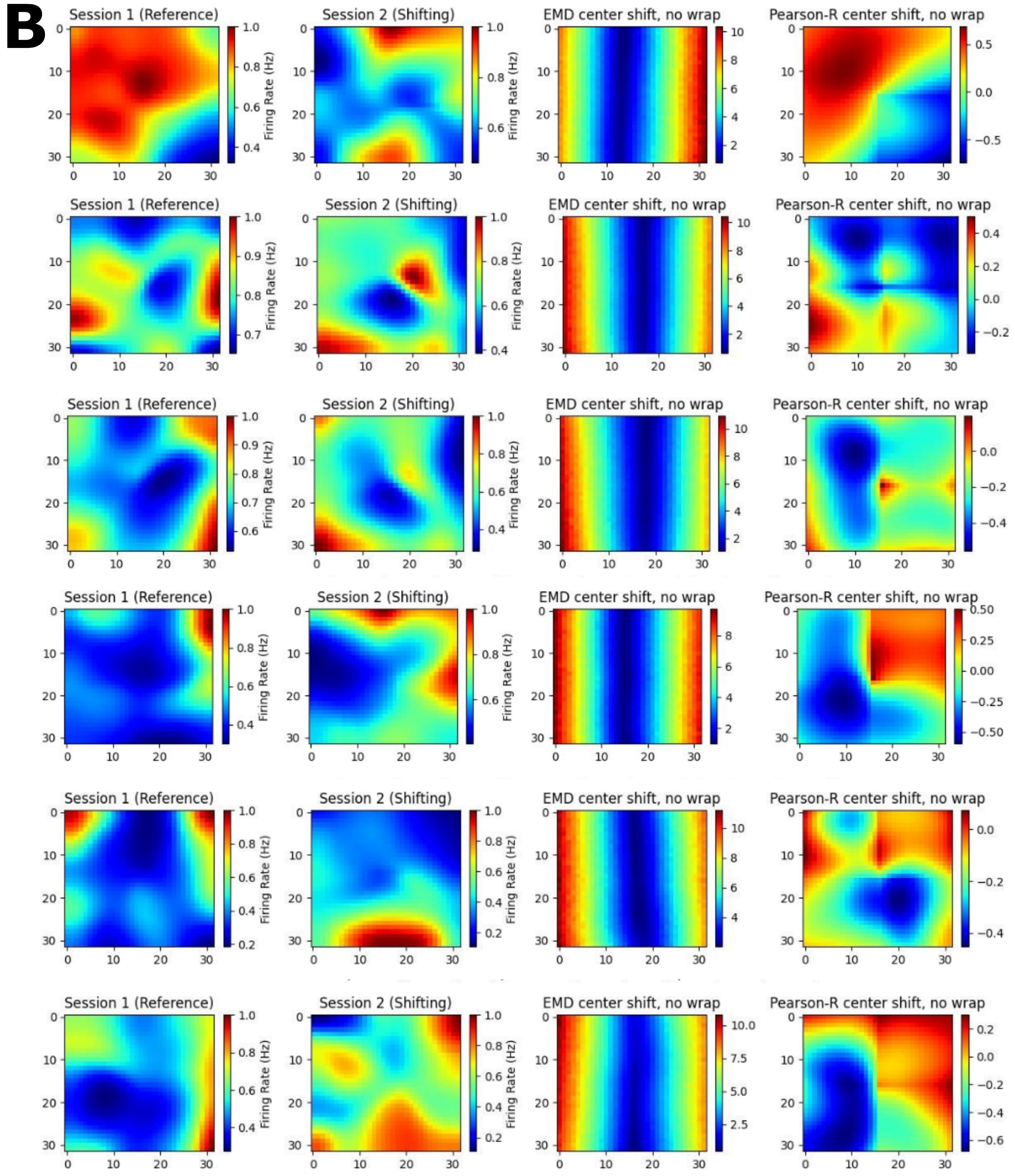


**Figure S13. Additional HPC cell examples**. Additional examples of ratemaps from the HPC of AD mouse models for a reference session and a shifting session. Gradient of Pearson's r scores tested at all possible map shift centers (N*N) is shown to the far right of each cell example with the EMD gradient immediately to the left of it. For each HPC example, only the no wrap row is provided, and examples of matched cells across circular to rectangular arena transitions are included.

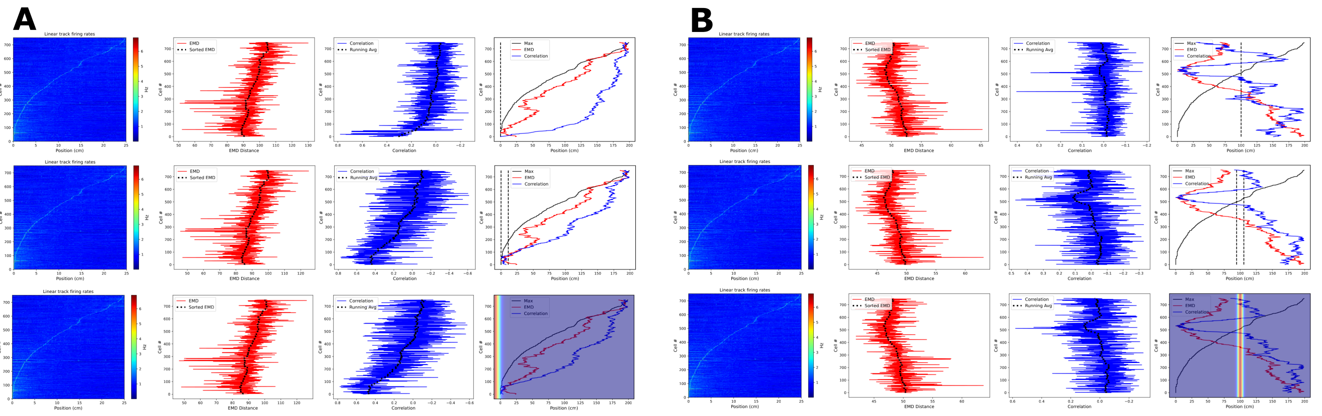


**Figure S14. Alternative place cell population decoding**. Place cell population decoding using a reference template and a population of 1D fluorescence values ($\Delta F/F_0$) across a 200cm linear track. The first plot in a row shows a population map of fluorescence. The second plot shows the distribution of EMD values, and the running average. The third plot shows the population of correlation values, and the running average. The last plot shows the peak firing rate trend overlaid with the running averages of each score distribution. Two sets of examples are provided with reference points at the start (A) and in the center (B). In each set, the first row of plots uses a reference map with all the activity in a single bin. The second row uses a map with the activity spread out across 10 bins. The last row uses a 1D gaussian template with sigma = 5. Reference points, and windows, are plotted with a dashed black line in the final plot of each row. Reference gaussians are overlaid in the final plot of each row.
