## Supplementary Figures (Spearman) for "Beyond Correlation: Optimal Transport Metrics For Characterizing Representational Stability and Remapping in Neurons Encoding Spatial Memory"

### 3 SPEARMAN COMPARISON

A sample of figures were re-generated with the spearman-$\rho$ function instead of the Pearson's r function. While plot labels may indicate pearson, every figure below this point has been re-generated with spearman-$\rho$.

#### 3.1 Figures

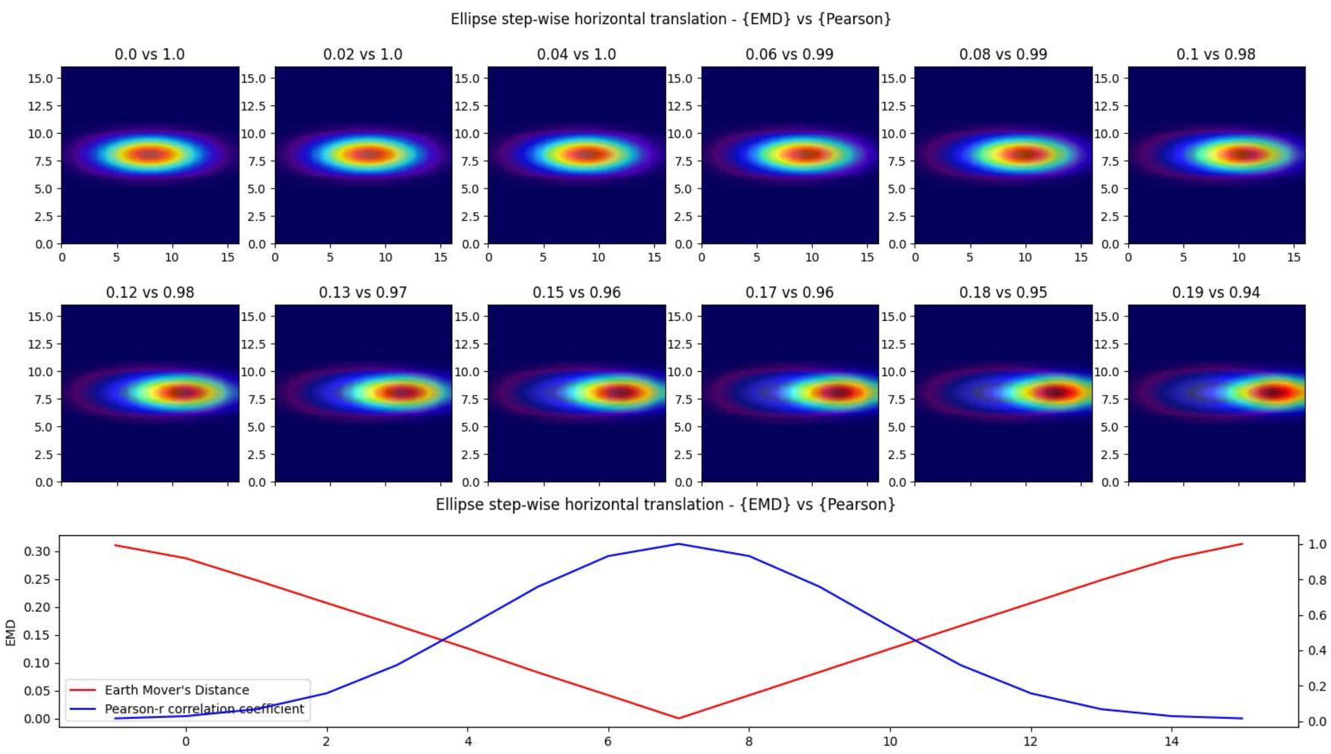
Ellipse step-wise horizontal translation - {EMD} vs {Pearson}
0.0 vs 1.0
0.02 vs 1.0
0.04 vs 1.0
0.06 vs 0.99
0.08 vs 0.99
0.1 vs 0.98
0.12 vs 0.98
0.13 vs 0.97
0.15 vs 0.96
0.17 vs 0.96
0.18 vs 0.95
0.19 vs 0.94
Ellipse step-wise horizontal translation - {EMD} vs {Pearson}
EMD
Earth Mover's Distance
Pearson-r correlation coefficient

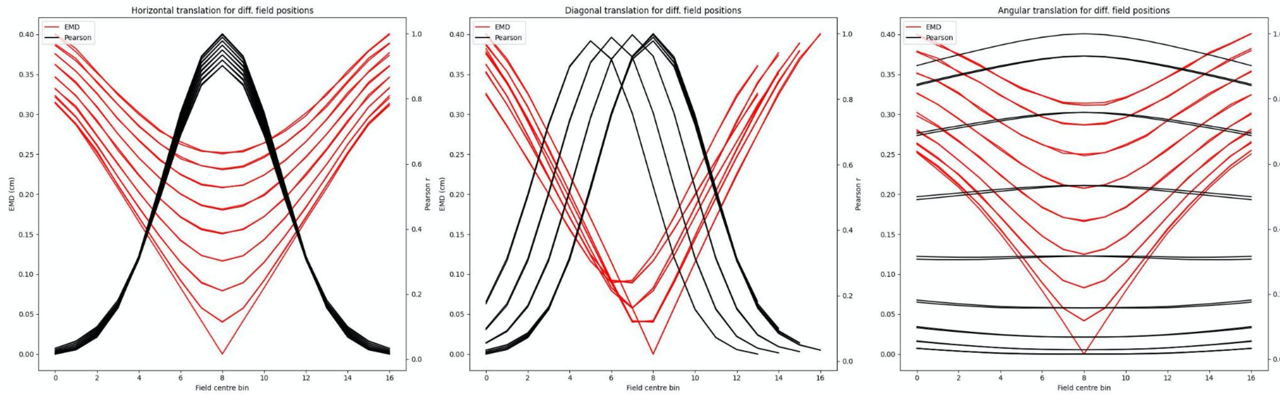
Horizontal translation for diff. field positions
Diagonal translation for diff. field positions
Angular translation for diff. field positions
EMD
Pearson
EMD (cm)
Pearson r
Field centre bin

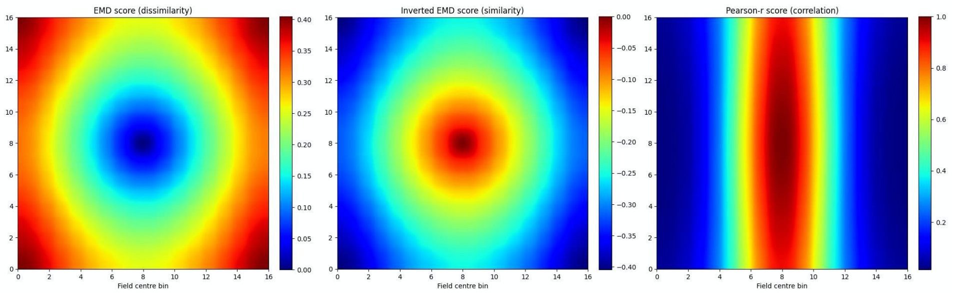
EMD score (dissimilarity)
Inverted EMD score (similarity)
Pearson-r score (correlation)
Field centre bin

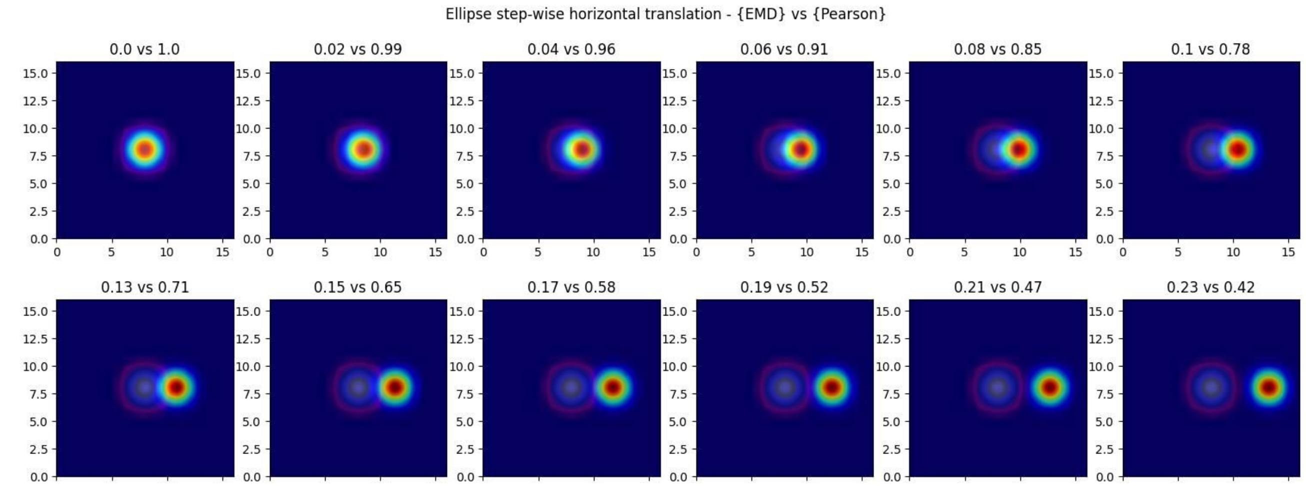
Ellipse step-wise horizontal translation - {EMD} vs {Pearson}
0.0 vs 1.0
0.02 vs 0.99
0.04 vs 0.96
0.06 vs 0.91
0.08 vs 0.85
0.1 vs 0.78
0.13 vs 0.71
0.15 vs 0.65
0.17 vs 0.58
0.19 vs 0.52
0.21 vs 0.47
0.23 vs 0.42

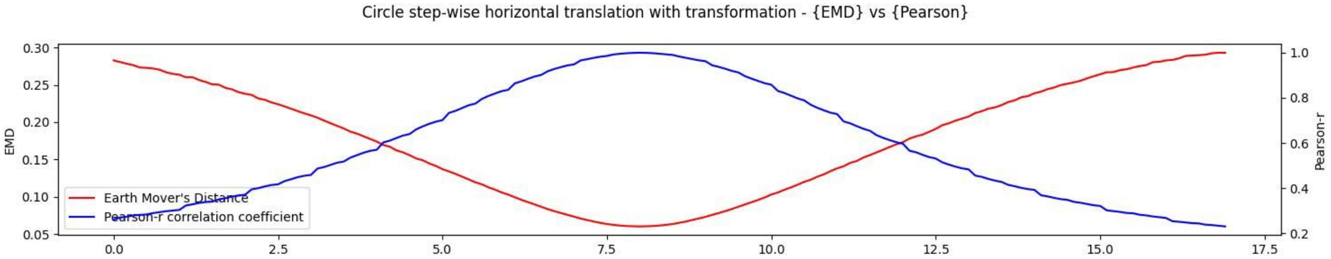
Circle step-wise horizontal translation with transformation - {EMD} vs {Pearson}
EMD
Pearson-r
Earth Mover's Distance
Pearson-r correlation coefficient

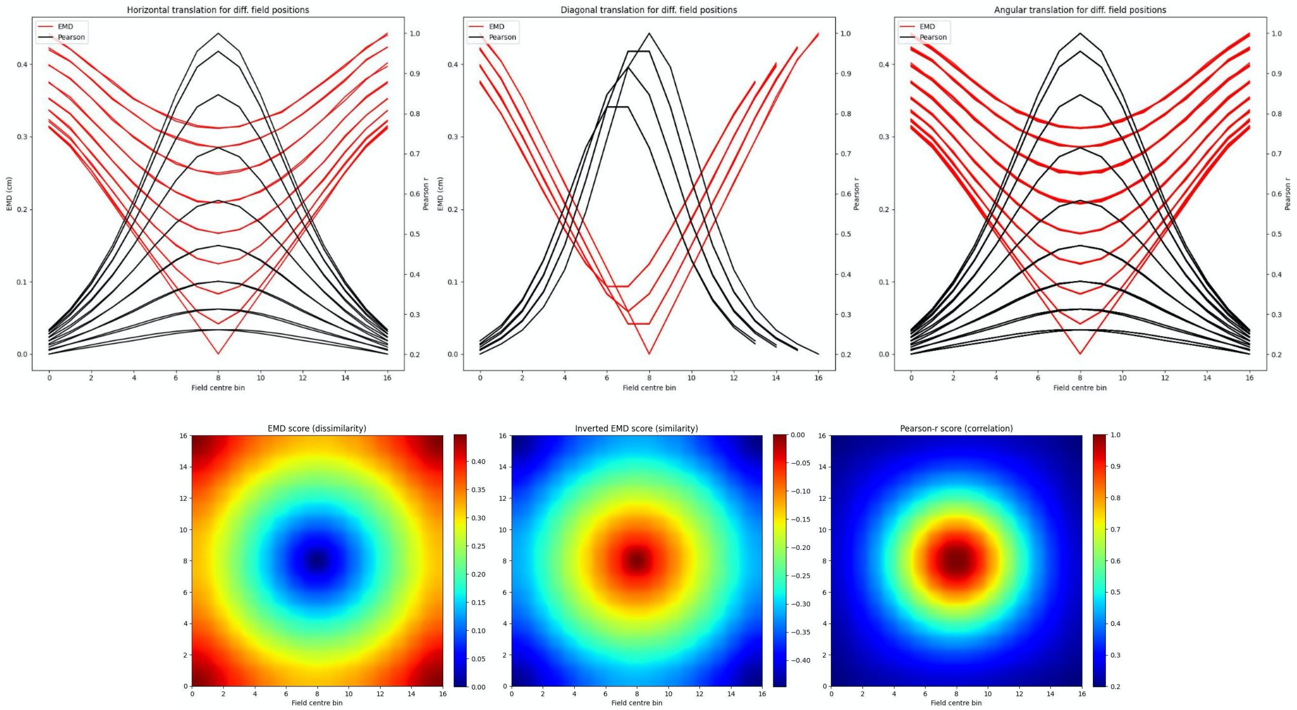


**Figure S12. Identical place field translation**. Stepwise horizontal linear translation of identical, overlapping place fields ($N = 17$, $\sigma = 3$) moving from the center to the right (**A, C**). EMD score is shown on the left while Pearson's r is shown on the right (top panel - EMD vs Pearson). 12 steps are shown and scores are rounded for display. Scores from remapping tested at all possible centroids along a single row on the rate map (bottom panel). EMD and Pearson's r scores tested at all possible centroids in the rate map (N*N) (**B, D**). Scores for horizontal and diagonal translations along the rate map are shown for all rows ($N = 17$) (top panel). Heatmap showing the gradient of scores for both raw and inverted EMD (left and center) and for Pearson's r (right) (bottom panel).

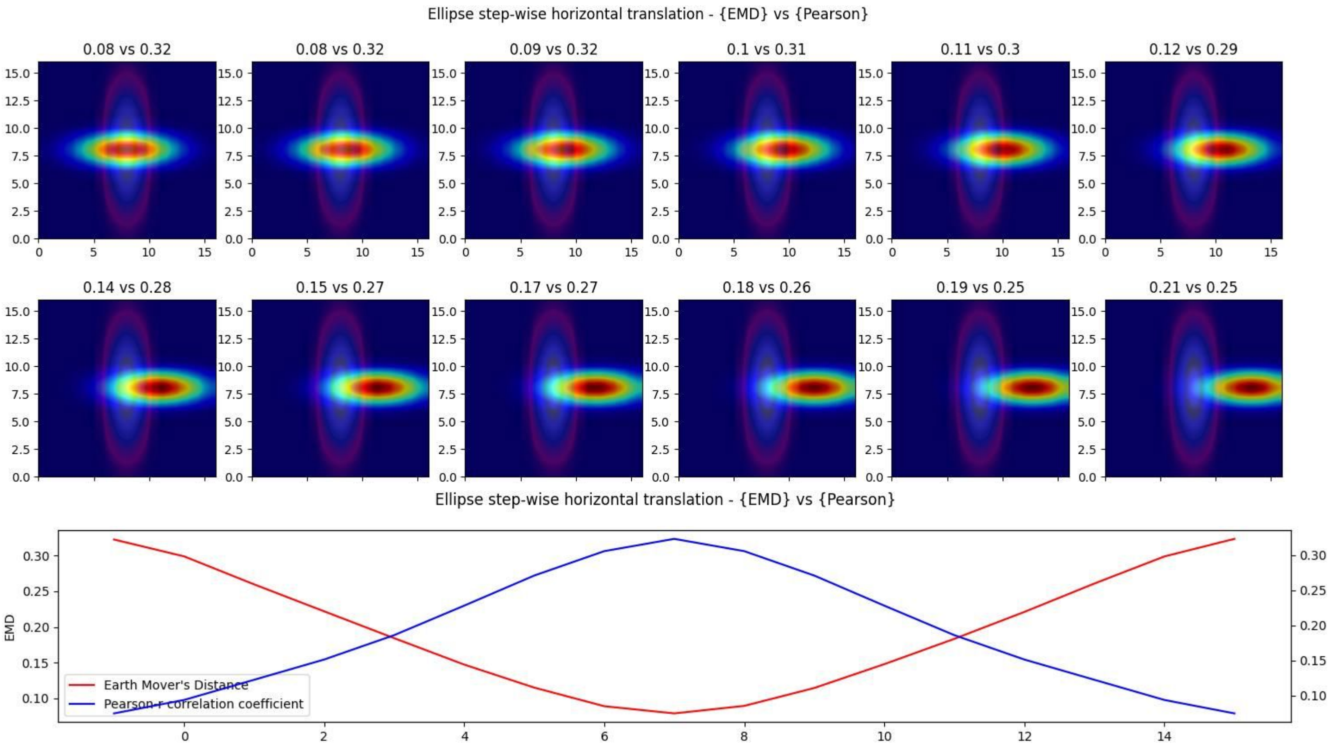
Ellipse step-wise horizontal translation - {EMD} vs {Pearson}
0.08 vs 0.32
0.08 vs 0.32
0.09 vs 0.32
0.1 vs 0.31
0.11 vs 0.3
0.12 vs 0.29
0.14 vs 0.28
0.15 vs 0.27
0.17 vs 0.27
0.18 vs 0.26
0.19 vs 0.25
0.21 vs 0.25
Ellipse step-wise horizontal translation - {EMD} vs {Pearson}
EMD
Earth Mover's Distance
Pearson-r correlation coefficient

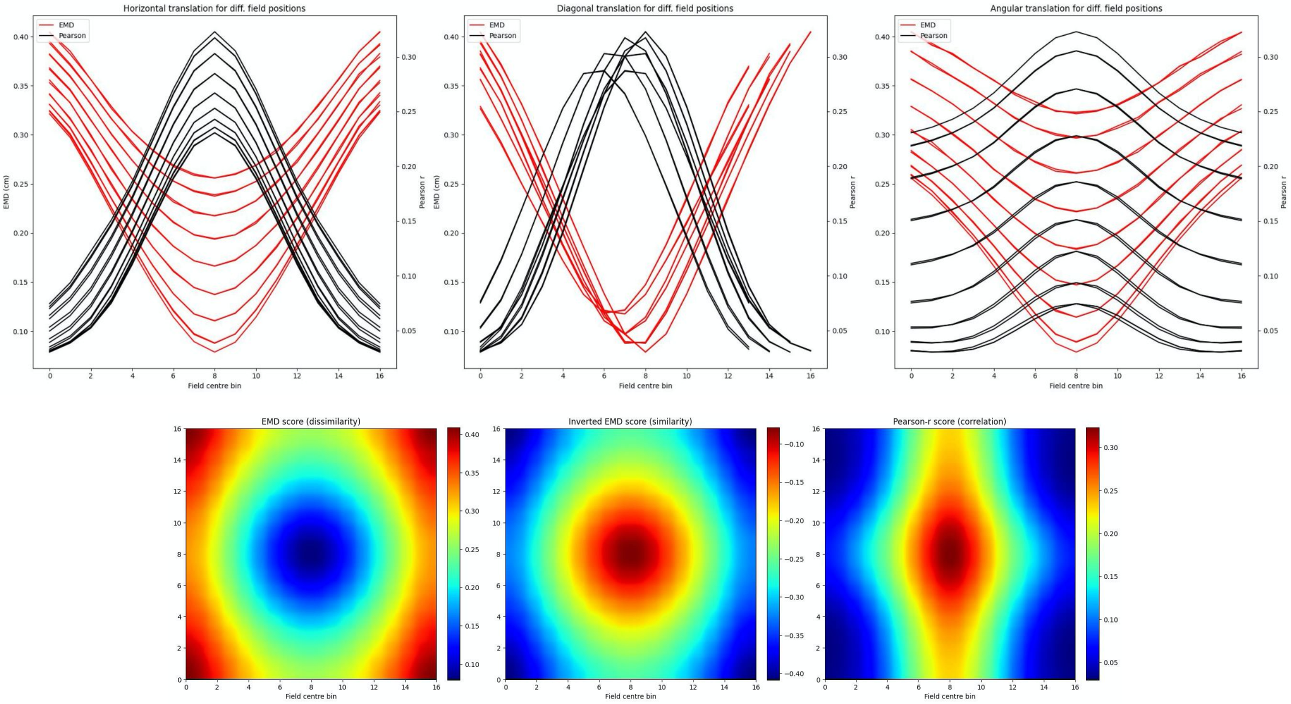
Horizontal translation for diff. field positions
Diagonal translation for diff. field positions
Angular translation for diff. field positions
EMD
Pearson
EMD (cm)
Pearson r
Field centre bin
EMD score (dissimilarity)
Inverted EMD score (similarity)
Pearson-r score (correlation)

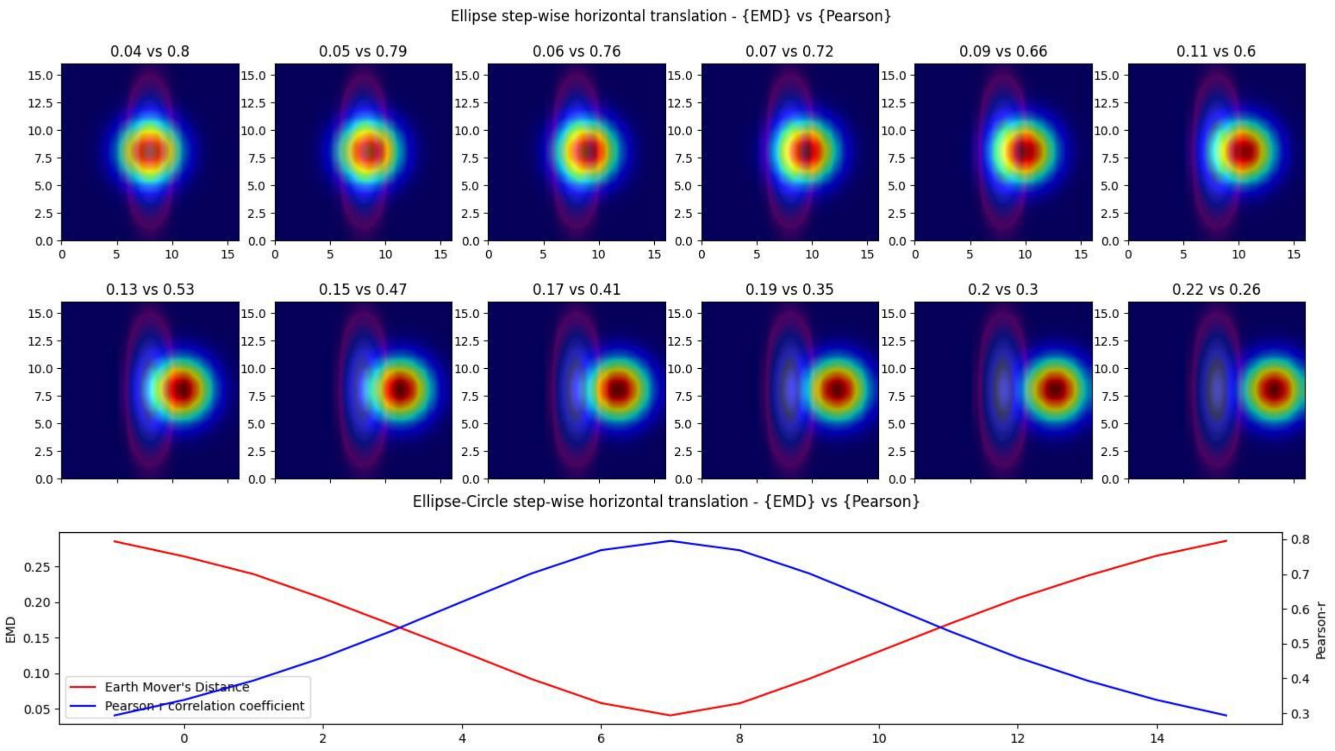
Ellipse step-wise horizontal translation - {EMD} vs {Pearson}
0.04 vs 0.8
0.05 vs 0.79
0.06 vs 0.76
0.07 vs 0.72
0.09 vs 0.66
0.11 vs 0.6
0.13 vs 0.53
0.15 vs 0.47
0.17 vs 0.41
0.19 vs 0.35
0.2 vs 0.3
0.22 vs 0.26
Ellipse-Circle step-wise horizontal translation - {EMD} vs {Pearson}
EMD
Pearson-r
Earth Mover's Distance
Pearson r correlation coefficient

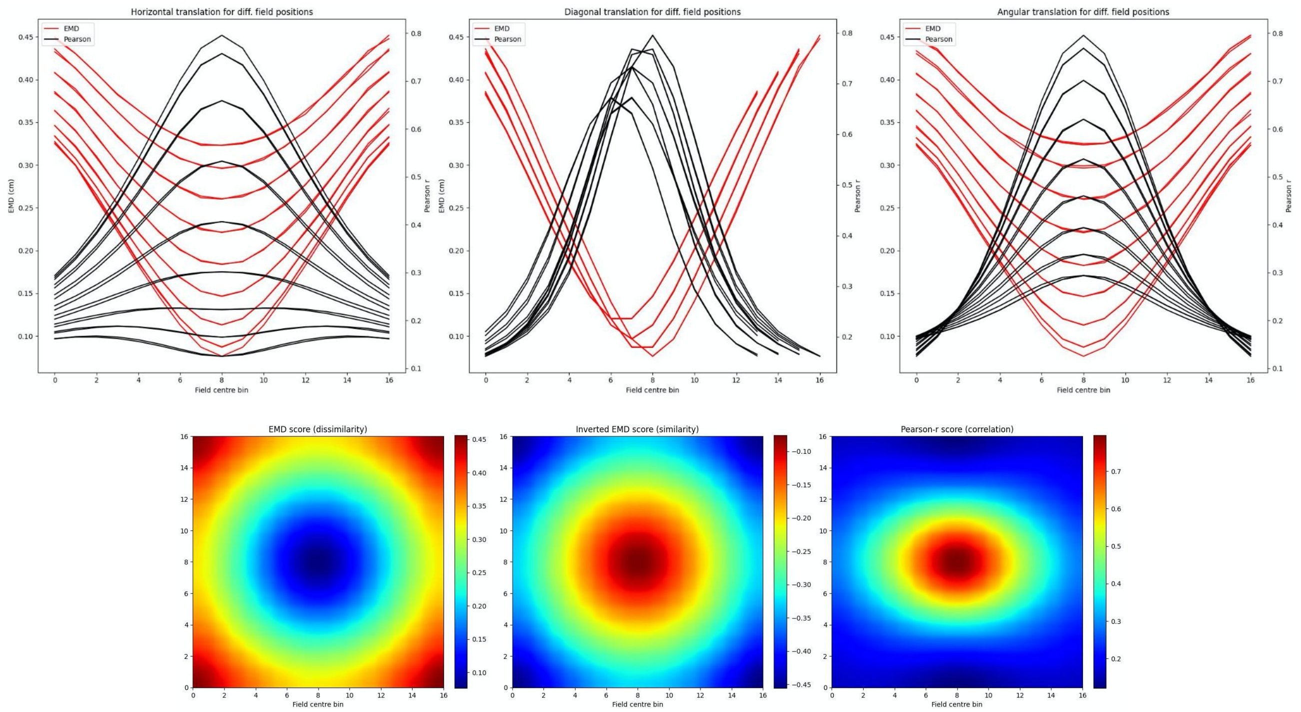


**Figure S13. Non-identical place field translation**. Stepwise horizontal linear translation of non-identical, overlapping place fields ($N = 17$, $\sigma = 3$) moving from the center to the right (**A, C**). EMD score is shown on the left while Pearson's r is shown on the right. 12 steps are shown and scores are rounded for display (top panel). Scores from remapping tested at all possible centroids in a single row on the rate map (bottom panel). EMD and Pearson's r scores tested at all possible centroids in the rate map (N*N) (**B, D**). Horizontal and diagonal translations across the rate map are shown for all rows ($N = 17$) (top panel). Heatmap showing the gradient of EMD scores both raw and inverted to match Pearson's r color scheme (bottom panel).

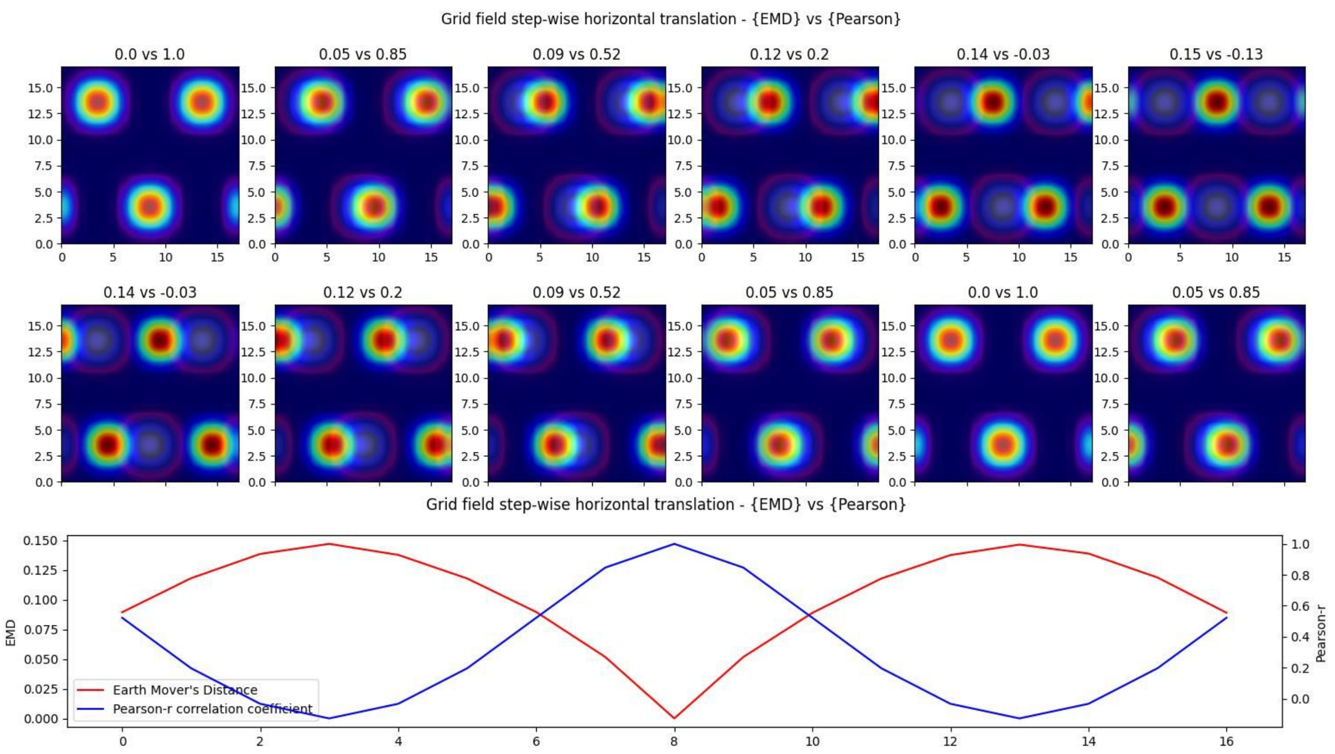
Grid field step-wise horizontal translation - {EMD} vs {Pearson}
0.0 vs 1.0
0.05 vs 0.85
0.09 vs 0.52
0.12 vs 0.2
0.14 vs -0.03
0.15 vs -0.13
0.14 vs -0.03
0.12 vs 0.2
0.09 vs 0.52
0.05 vs 0.85
0.0 vs 1.0
0.05 vs 0.85
Grid field step-wise horizontal translation - {EMD} vs {Pearson}
EMD
Pearson-r
Earth Mover's Distance
Pearson-r correlation coefficient

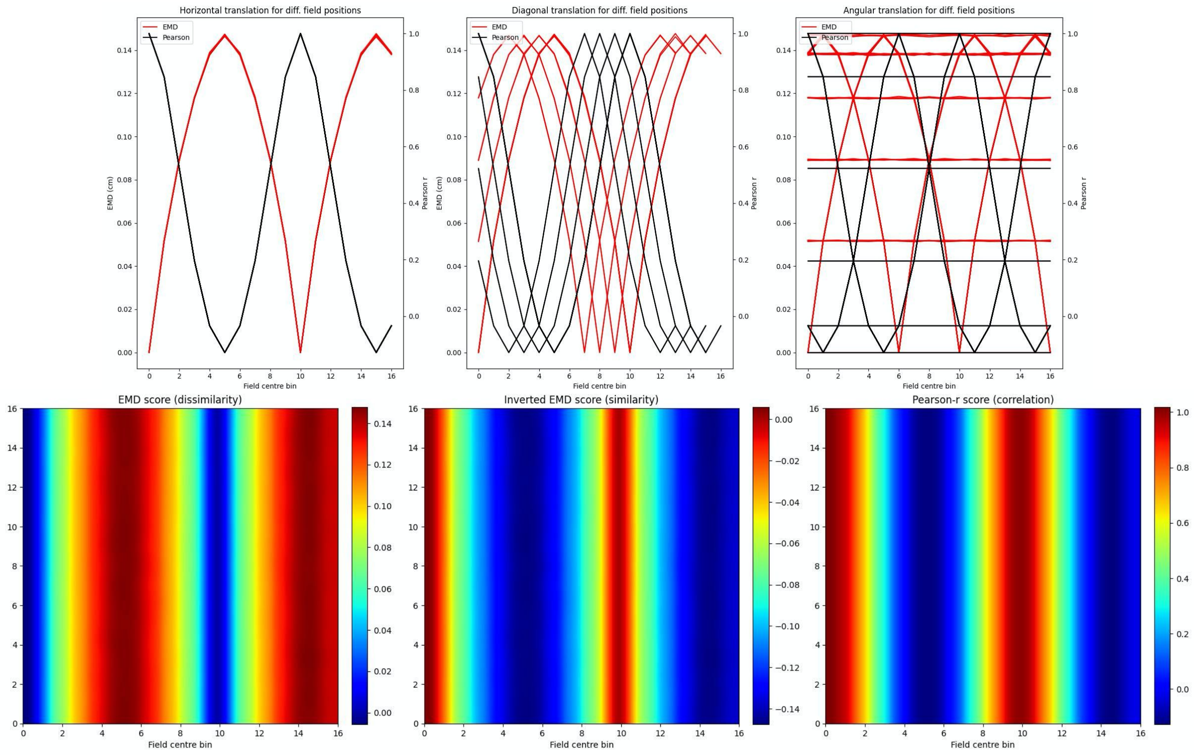
Horizontal translation for diff. field positions
Diagonal translation for diff. field positions
Angular translation for diff. field positions
EMD
Pearson
EMD (cm)
Pearson r
Field centre bin
EMD score (dissimilarity)
Inverted EMD score (similarity)
Pearson-r score (correlation)

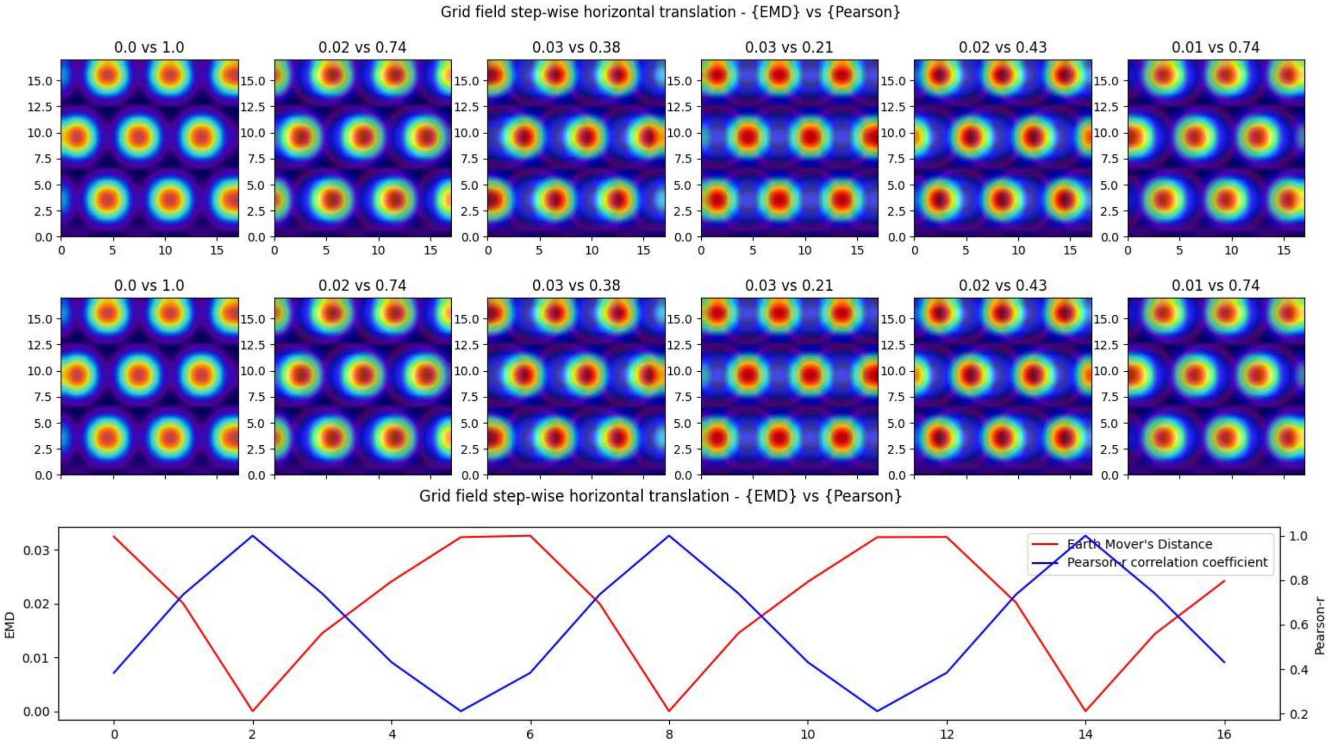
Grid field step-wise horizontal translation - {EMD} vs {Pearson}
0.0 vs 1.0
0.02 vs 0.74
0.03 vs 0.38
0.03 vs 0.21
0.02 vs 0.43
0.01 vs 0.74
Grid field step-wise horizontal translation - {EMD} vs {Pearson}
EMD
Pearson-r
Earth Mover's Distance
Pearson-r correlation coefficient

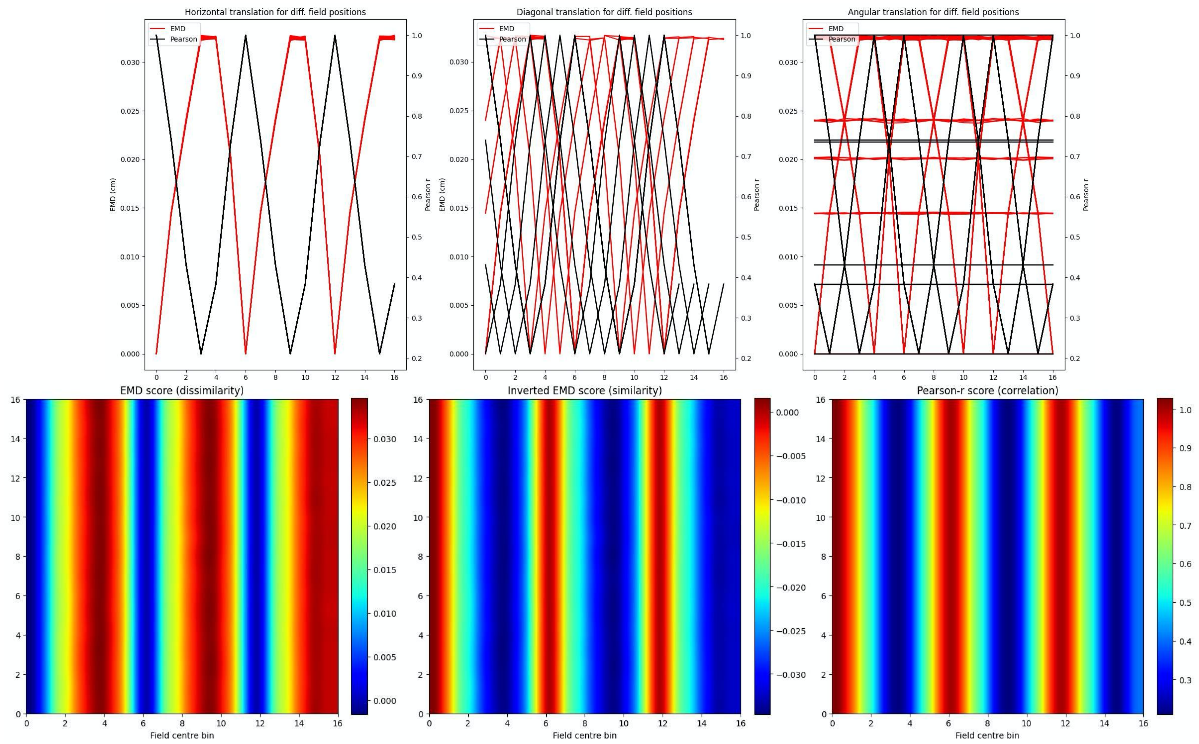


**Figure S14. Identical grid field translation**. Stepwise horizontal linear translation of identical, overlapping grid fields ($N = 3$, $\sigma = 1$) moving from the top left corner to the right and/or downwards on a rate map ($N = 17$) (**A, C**). EMD score is shown on the left while Pearson's r is shown on the right. 12 steps are shown and scores are rounded for display (top panel - EMD vs Pearson). Grid maps were sliced from a larger map with sufficient fields and bins to support N*N steps. Initial grid maps were chosen by taking a slice from the wider map. Scores from remapping tested across N*N different shifts from the initial grid map (0 to N combinations) (bottom panel). EMD and Pearson's r scores tested at N*N different centroid positions on the wider grid (**B, D**). Scores for horizontal and diagonal translations along the rate map are shown for all rows ($N = 17$) (top panel). Heatmap showing the gradient of scores for both raw and inverted EMD (left and center) and for Pearson's r (right) (bottom panel).

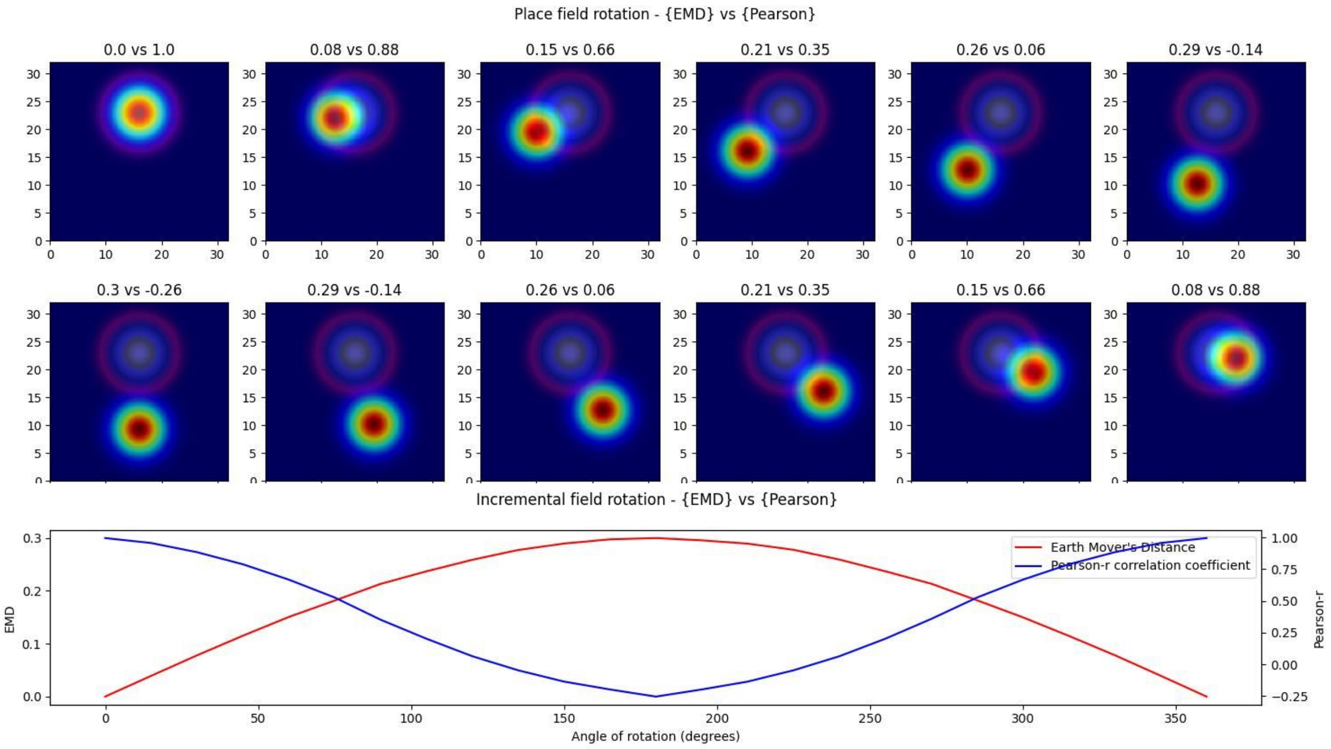
Place field rotation - {EMD} vs {Pearson}
0.0 vs 1.0
0.08 vs 0.88
0.15 vs 0.66
0.21 vs 0.35
0.26 vs 0.06
0.29 vs -0.14
0.3 vs -0.26
0.29 vs -0.14
0.26 vs 0.06
0.21 vs 0.35
0.15 vs 0.66
0.08 vs 0.88
Incremental field rotation - {EMD} vs {Pearson}
Earth Mover's Distance
Pearson-r correlation coefficient
EMD
Pearson-r
Angle of rotation (degrees)

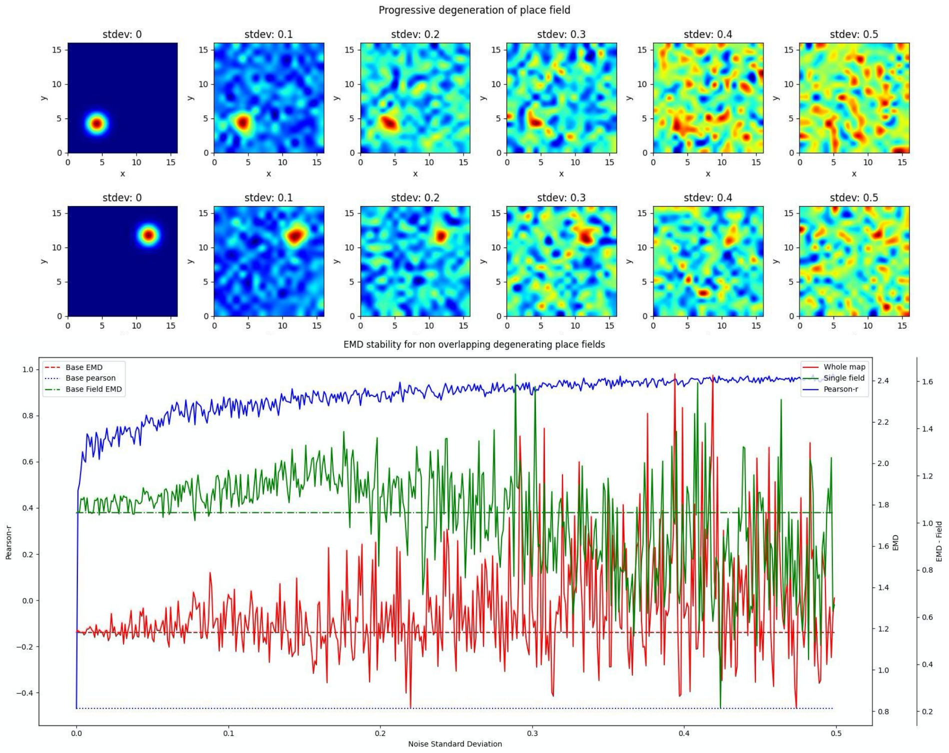
Progressive degeneration of place field
stdev: 0
stdev: 0.1
stdev: 0.2
stdev: 0.3
stdev: 0.4
stdev: 0.5
EMD stability for non overlapping degenerating place fields
Base EMD
Base pearson
Base Field EMD
Whole map
Single field
Pearson-r
Pearson-r
EMD
EMD - Field
Noise Standard Deviation

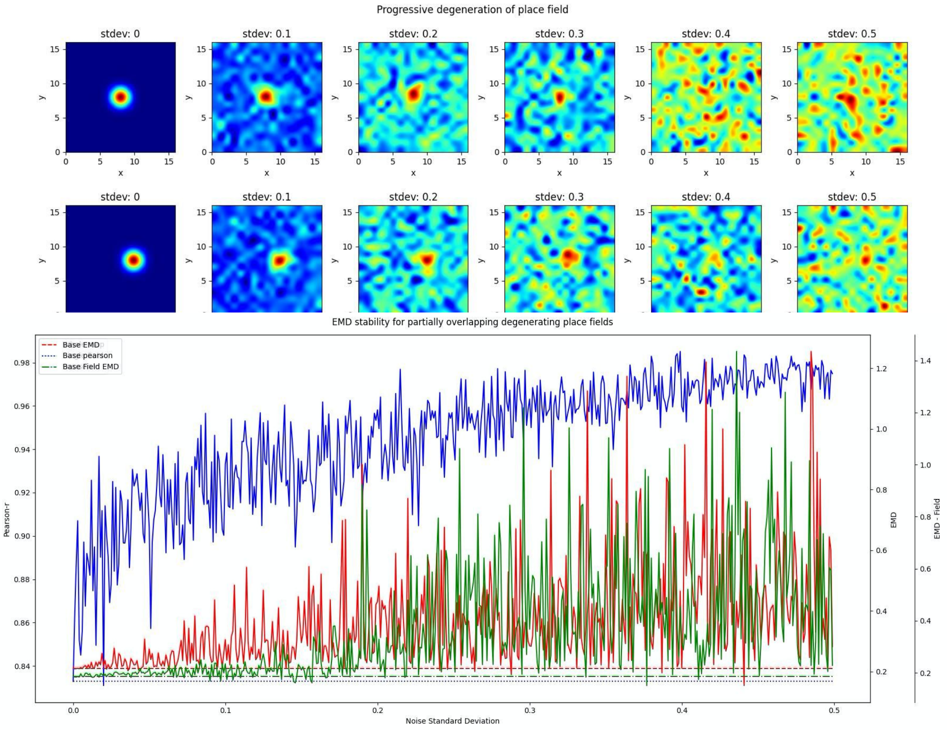


**Figure S15. Incremental field degeneration**. Stepwise nonlinear translation of overlapping fields ($N = 33$) relative to a fixed field at $\sigma = 3$. 12 steps are shown with 6 scaling down and 6 scaling up relative to the fixed field (**A**). EMD score is shown on the left while Pearson's r is shown on the right (left panel). Scores from remapping tested across a range of standard deviations for the scaling field (right panel). Incremental field degeneration for a pair of fields, non-overlapping and overlapping (**B,C**). Left panels show the stepwise degradation in the rate map due to randomly sampled normally distributed noise with varying standard deviations. Noise standard deviations are shown above the rate map plots. The distribution plots show the computed remapping score between the pair of fields for the overlapping and non-overlapping cases. Both cases have values for the EMD (red), field EMD (green) and Pearson's r scores (blue) displayed (right panels).

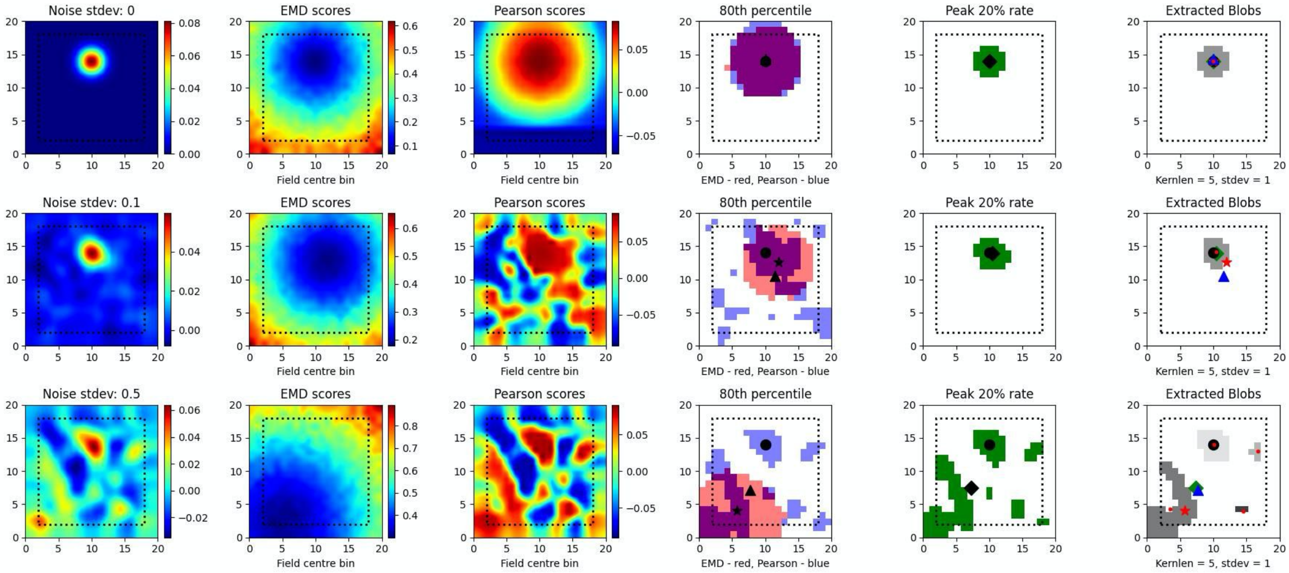
Noise stdev: 0
EMD scores
Pearson scores
80th percentile
Peak 20% rate
Extracted Blobs
Noise stdev: 0.1
Noise stdev: 0.5
Field centre bin
EMD - red, Pearson - blue
Kernlen = 5, stdev = 1

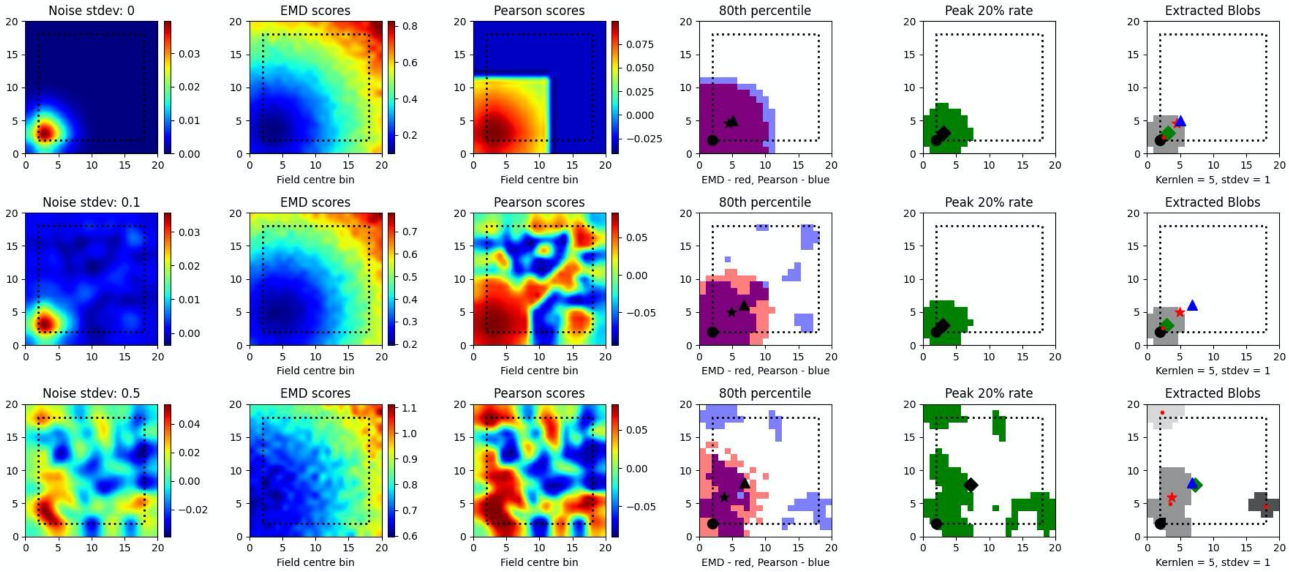
Noise stdev: 0
EMD scores
Pearson scores
80th percentile
Peak 20% rate
Extracted Blobs
Noise stdev: 0.1
Noise stdev: 0.5
Field centre bin
EMD - red, Pearson - blue
Kernlen = 5, stdev = 1

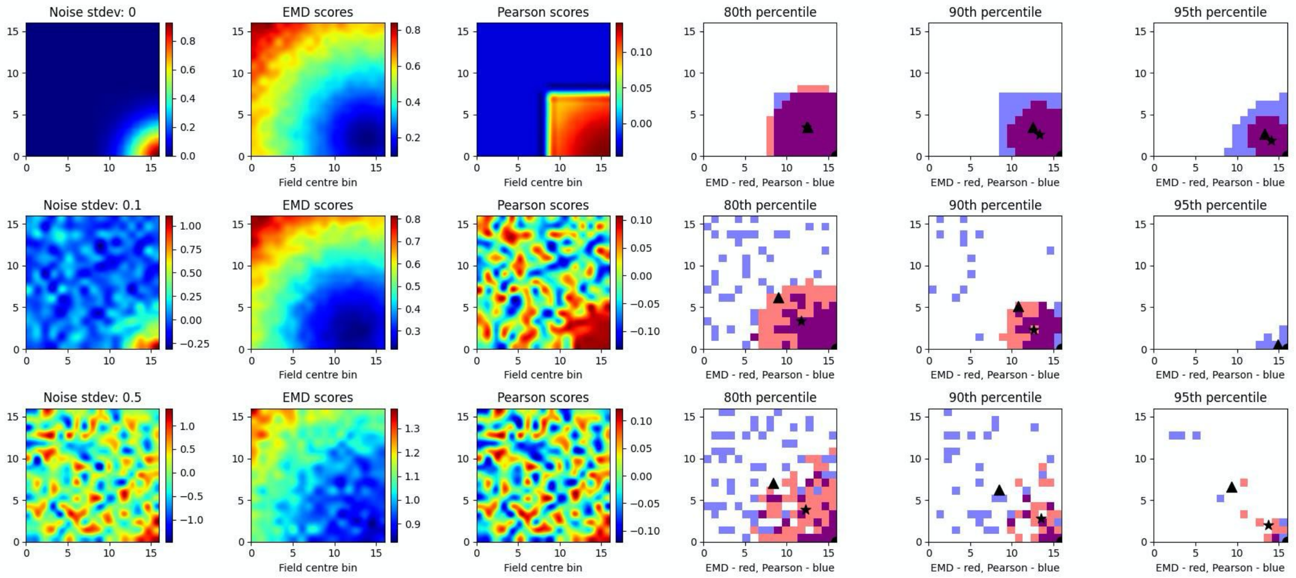
Noise stdev: 0
EMD scores
Pearson scores
80th percentile
90th percentile
95th percentile
Noise stdev: 0.1
Noise stdev: 0.5
Field centre bin
EMD - red, Pearson - blue

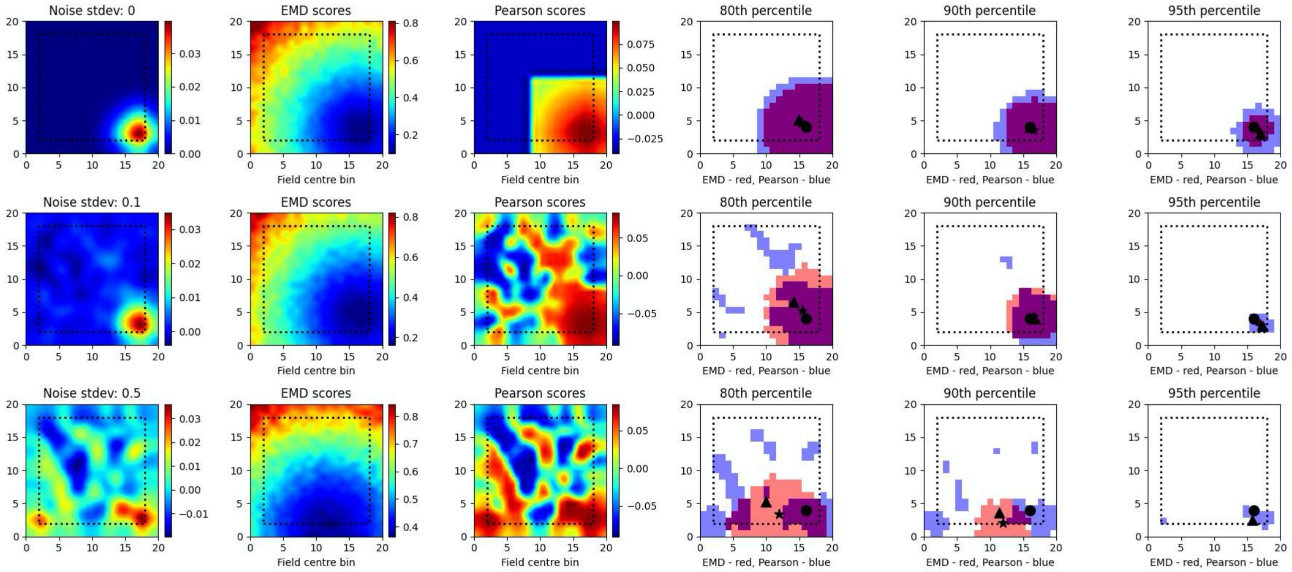
Noise stdev: 0
EMD scores
Pearson scores
80th percentile
90th percentile
95th percentile
Noise stdev: 0.1
Noise stdev: 0.5
Field centre bin
EMD - red, Pearson - blue

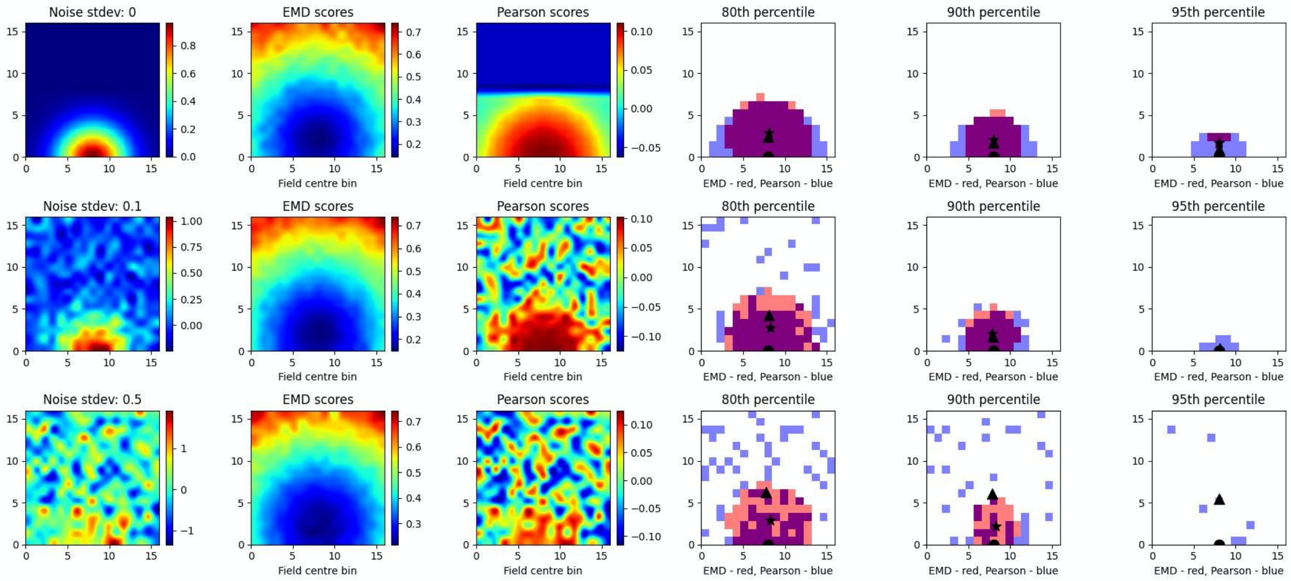


**Figure S16. Single field localization**. Field localization plots across two different noise levels (rows: low noise 0.1 and high noise 0.5). For each row in a plot, the first column shows the ratemap post added noise with padding, smoothing and normalizing. The second column shows the EMD distribution on the padded rate map with scores being relative to a fake point map with all the density placed in the bin at which the EMD score is found. The third column shows the same map to point computation for Pearson's r scores. The fourth column shows the 80th percentile scores for the EMD (red) and Pearson's r distributions (blue). The fifth column shows the top 20% firing rates in the cell. The last column (sixth) holds the extracted blobs (fields) from the padded ratemap with the centroid of each blob shown in red. The circle represents the true field centroid. The star represents the centroid computed on the peak EMD scores. The triangle is the centroid computed from the peak Pearson's r scores. The diamond is the centroid from the peak firing rates. The red dots are the centroids of a given field

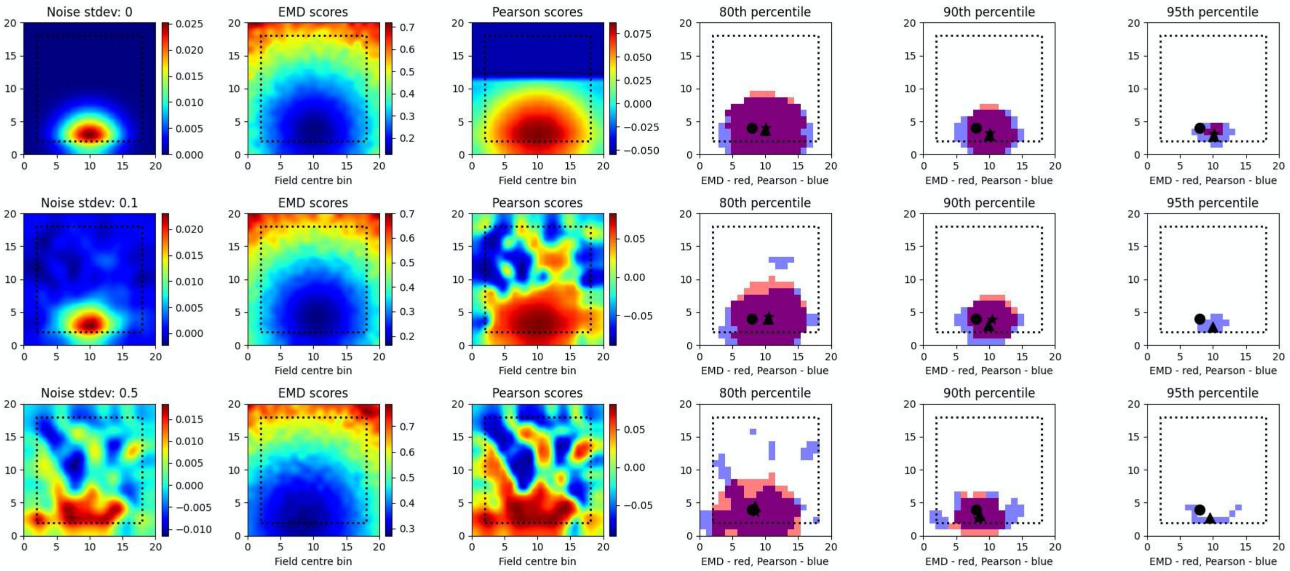
Noise stdev: 0
EMD scores
Pearson scores
80th percentile
90th percentile
95th percentile
Field centre bin
EMD - red, Pearson - blue
Noise stdev: 0.1
Noise stdev: 0.5

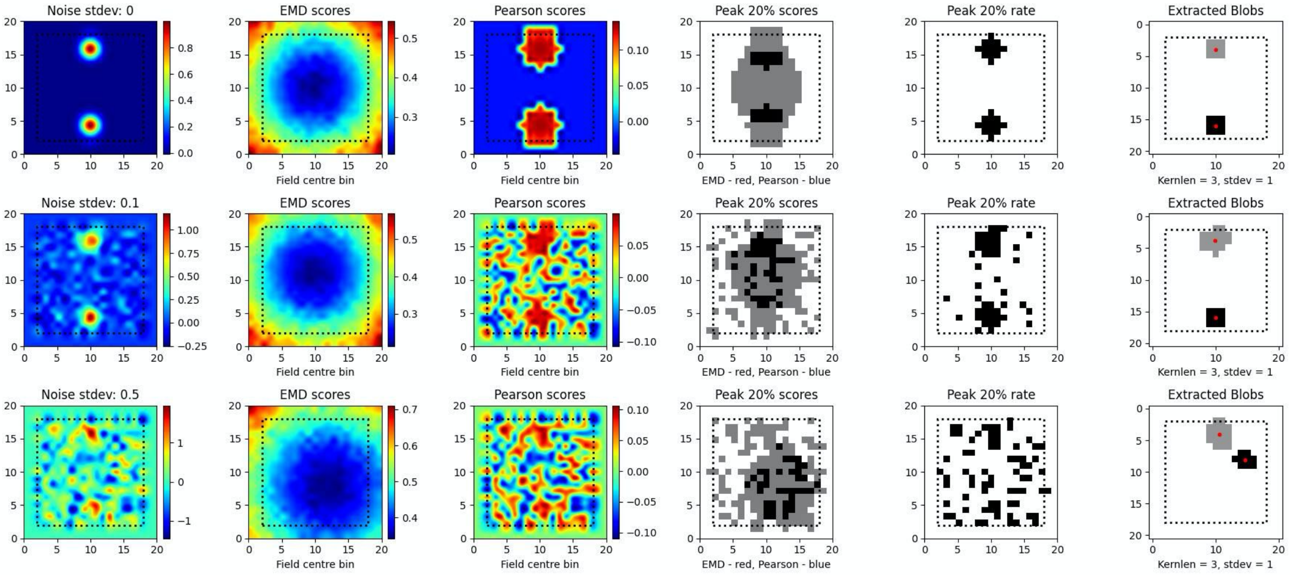
Noise stdev: 0
EMD scores
Pearson scores
Peak 20% scores
Peak 20% rate
Extracted Blobs
Field centre bin
EMD - red, Pearson - blue
Kernlen = 3, stdev = 1
Noise stdev: 0.1
Noise stdev: 0.5

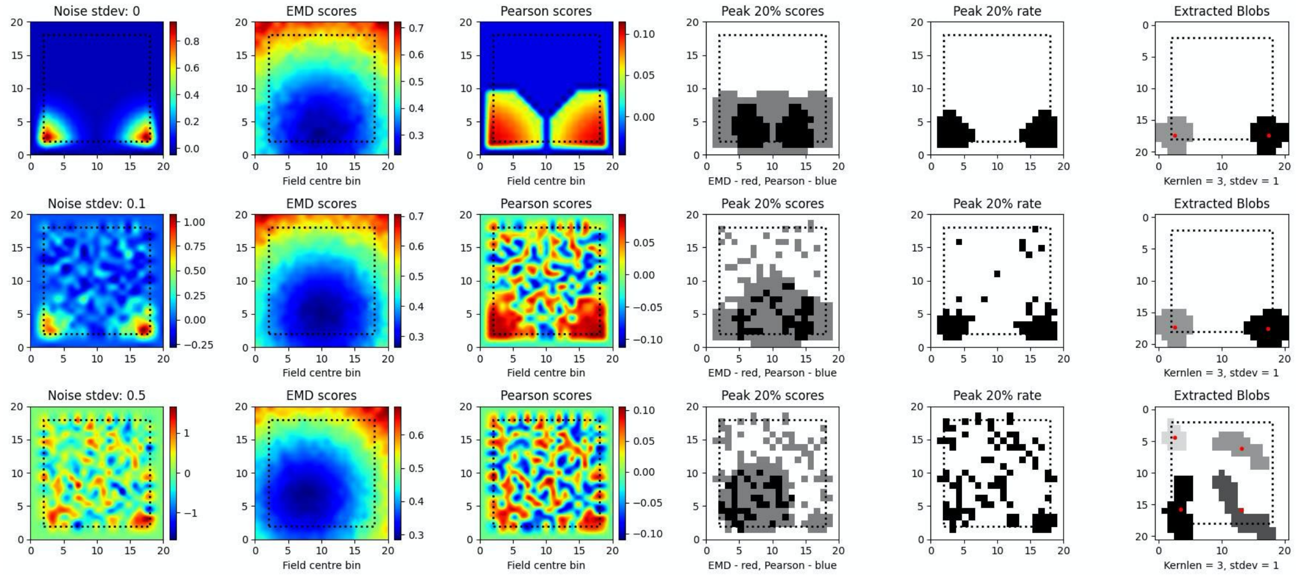
Noise stdev: 0
EMD scores
Pearson scores
Peak 20% scores
Peak 20% rate
Extracted Blobs
Field centre bin
EMD - red, Pearson - blue
Kernlen = 3, stdev = 1
Noise stdev: 0.1
Noise stdev: 0.5

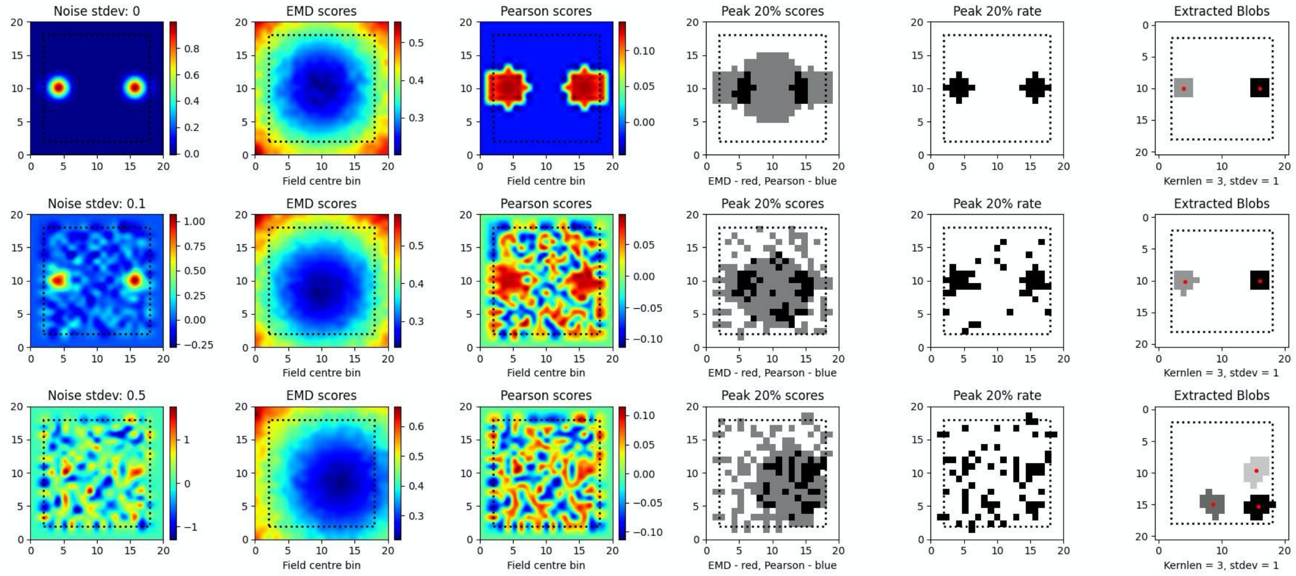
Noise stdev: 0
EMD scores
Pearson scores
Peak 20% scores
Peak 20% rate
Extracted Blobs
Field centre bin
EMD - red, Pearson - blue
Kernlen = 3, stdev = 1
Noise stdev: 0.1
Noise stdev: 0.5

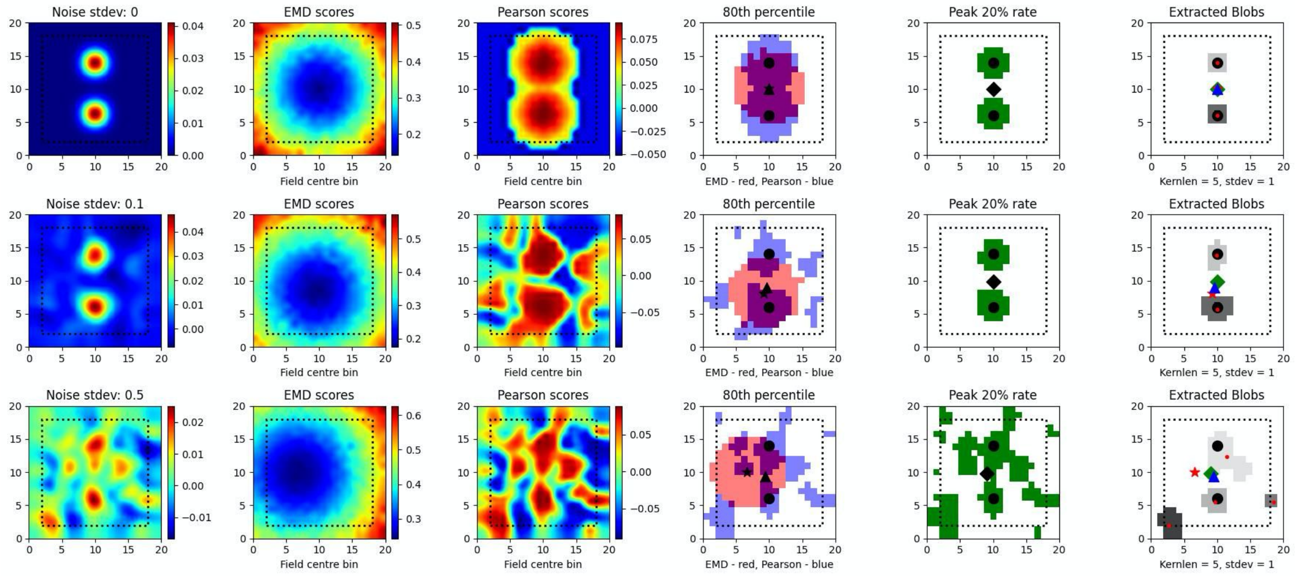
Noise stdev: 0
EMD scores
Pearson scores
80th percentile
Peak 20% rate
Extracted Blobs
Field centre bin
EMD - red, Pearson - blue
Kernlen = 5, stdev = 1
Noise stdev: 0.1
Noise stdev: 0.5

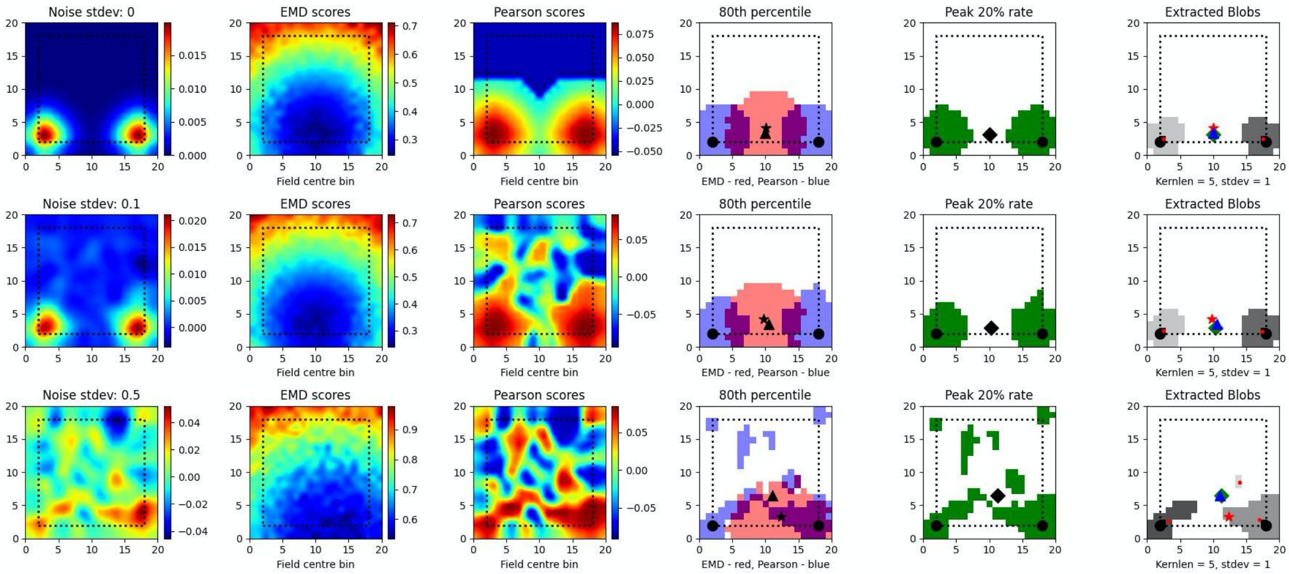


**Figure S17. Dual field localization**. Field localization plots across two different noise levels (rows: low noise 0.1 and high noise 0.5). For each row in a plot, the first column shows the ratemap post added noise with padding, smoothing and normalizing. The second column shows the EMD distribution on the padded rate map with scores being relative to a fake point map with all the density placed in the bin at which the EMD score is found. The third column shows the same map to point computation for Pearson's r scores. The fourth column shows the 80th percentile scores for the EMD (red) and Pearson's r distributions (blue). The fifth column shows the top 20% firing rates in the cell. The last column (sixth) holds the extracted blobs (fields) from the padded ratemap with the centroid of each blob shown in red. The circle represents the true field centroid. The star represents the centroid computed on the peak EMD scores. The triangle is the centroid computed from the peak Pearson's r scores. The diamond is the centroid from the peak firing rates. The red dots are the centroids of a given field

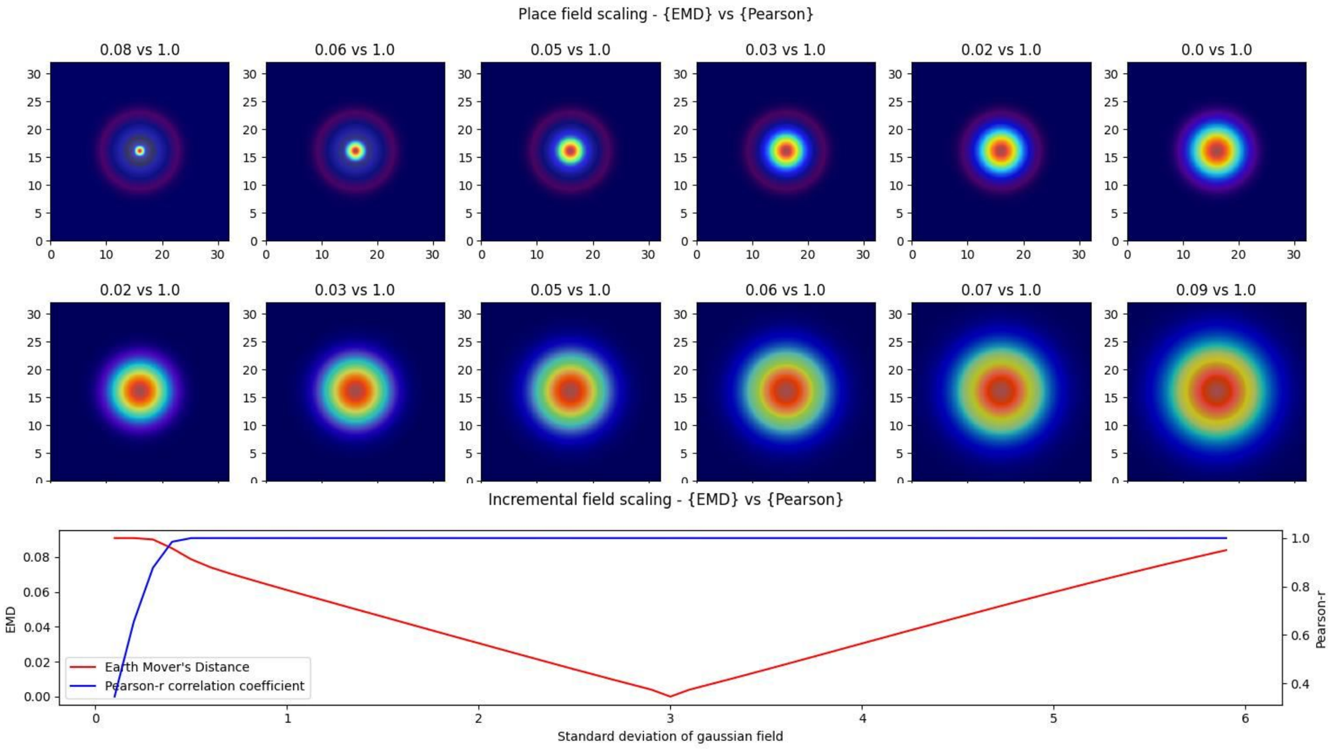
Place field scaling - {EMD} vs {Pearson}
0.08 vs 1.0
0.06 vs 1.0
0.05 vs 1.0
0.03 vs 1.0
0.02 vs 1.0
0.0 vs 1.0
0.02 vs 1.0
0.03 vs 1.0
0.05 vs 1.0
0.06 vs 1.0
0.07 vs 1.0
0.09 vs 1.0
Incremental field scaling - {EMD} vs {Pearson}
EMD
Pearson-r
Earth Mover's Distance
Pearson-r correlation coefficient
Standard deviation of gaussian field

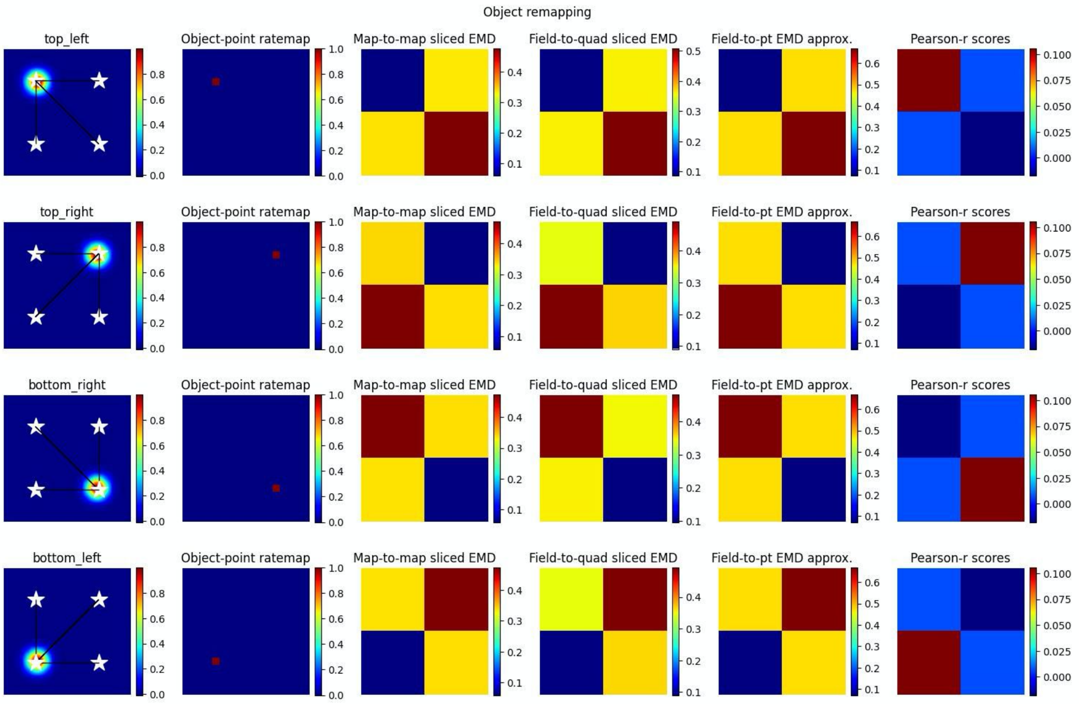


**Figure S18. Complex non-linear field remapping**. Stepwise nonlinear translation of overlapping fields ($N = 33$) relative to a fixed field at $\sigma = 3$. 12 steps are shown with 6 scaled down and 6 scaled up relative to the fixed field (A). EMD score is shown on the left while Pearson's r is shown on the right (top panel). Scores from remapping tested across a range of rotation angles (bottom panel). Four corner point driven remapping with top left, top right, bottom right and bottom left tested. Fields were positioned so as to be fully encompassed by the rate map area. The first column shows the field location, four possible object/point/stimulus locations (stars), and distances from the field centroid to each of the four positions. The second column shows the whole map to whole map EMD scores with the full rate map and a fake pointmap (1 at object location, 0 everywhere else). The third shows a field restricted EMD between a field and a quadrant of multiple bins. The fourth column shows an approximation to the whole map sliced EMD using only the field and the single point object point (single point Wasserstein). The last column holds the Pearson's r scores. Heatmaps demonstrate the scores in the four possible corners.
